# Imputation and polygenic score performance of low coverage whole-genome sequencing and genotyping arrays in diverse human populations

**DOI:** 10.1101/2025.07.18.665609

**Authors:** Phi Truong Nguyen, Thuy Vy Nguyen, Dat Thanh Nguyen, Thuy Duong Ho Huynh

**Affiliations:** KTest Vietnam, Ho Chi Minh City, Vietnam; Department of Genetics, Faculty of Biology and Biotechnology, University of Science, VNUHCM, Ho Chi Minh City, Vietnam; Centre for Integrative Genetics, Faculty of Biosciences, Norwegian University of Life Sciences, As, Norway; Center for Precision Psychiatry, Division of Mental Health and Addiction, University of Oslo, Oslo, Norway

## Abstract

Genome-wide association studies and polygenic score analysis rely on large-scale genotypic data, traditionally obtained through SNP arrays and imputation. However, low coverage whole-genome sequencing has emerged as a promising alternative. This study presents a comprehensive comparison of imputation accuracy and polygenic score performance between eight high-performance genotyping arrays and six low coverage whole-genome sequencing coverage levels (0.5–2x) across diverse populations. We analyze data from 2,504 individuals in the 1000 Genomes Project using a 10-fold cross-imputation strategy to evaluate imputation accuracy and polygenic score performance for four complex traits. Our results demonstrate that low-pass whole-genome sequencing performs competitively with population-specific arrays in both imputation accuracy and polygenic score estimation. Interestingly, low coverage whole-genome sequencing shows superior performances compared to arrays in underrepresented populations and for rare and low-frequency variants. Our findings suggest that low coverage whole-genome sequencing offers a flexible and powerful alternative to genotyping arrays for large-scale genetic studies, particularly in diverse or underrepresented populations.

## 1 Introduction

Genome-wide association studies (GWAS) have identified thousands of genetic variants linked to complex human traits. By 2023, more than 400,000 associations had been reported from over 6,000 studies in humans^[1]^. Research has shown that many complex traits exhibit a polygenic architecture, where numerous genetic variants with small effects contribute to phenotypic variation^[2,3,4]^. One key application of GWAS is estimating an individual’s genetic predisposition to specific phenotypes^[5,6,7,8]^. This predisposition is commonly quantified as a polygenic score (PGS), which aggregates an individual’s risk alleles weighted by their effect sizes derived from GWAS summary statistics^[9]^. With its potential to enhance precision medicine by improving disease stratification, identifying high-risk individuals, refining diagnoses, and predicting therapeutic outcomes, PGS has become a rapidly growing research area^[10,11,12,7,8]^.

Both GWAS and PGS applications typically require large-scale, genome-wide genotypic data. With advantages such as cost-effectiveness and low computational requirements^[13]^, SNP arrays followed by genotype imputation have been the dominant approach for obtaining genetic data in large-scale studies^[14,15]^. Despite their effectiveness, SNP arrays have several limitations. First, population-specific designs are necessary to maximize performance^[16]^. For instance, the UK Biobank Axiom Array^[17]^, Japonica NEO Arrays^[18]^, Axiom KoreanChip^[19]^, and FinnGen array are specifically designed to optimize imputation performance within their respective target populations. Second, the imputation accuracy of rare variants is limited^[20]^, as SNP arrays typically include a constrained set of variants. Consequently, variants with a minor allele frequency (MAF) below 1% are often omitted during array design^[21,22]^.

With decreasing sequencing costs, low coverage whole-genome sequencing or low-pass sequencing (LPS) has become a promising alternative to SNP arrays. Several studies highlighted its potential, including one using 1.7x coverage that identified genetic loci associated with major depressive disorder^[23]^. Another study demonstrated that 1x LPS could uncover associations missed by SNP array imputation^[24]^. Additionally, LPS with 0.5–1x coverage performs comparably to standard low-density arrays, while higher coverage (4×) improves the detection of novel variants in underrepresented populations^[25]^. Moreover, LPS provides similar accuracy at a comparable cost while reducing biases associated with SNP array design^[26]^.

As LPS gains traction, a comprehensive comparison with genotyping arrays is essential to help researchers choose the most suitable approach for specific study objectives and populations. While previous studies have examined LPS in relation to SNP arrays, their scope has been limited by a narrow selection of arrays or populations^[26,27,28,25,29,30]^. Here, we conduct an extensive in silico evaluation of imputation accuracy and PGS performance across eight high-performance human genotyping arrays and six LPS coverage levels ranging from 0.5-2x in diverse populations. Our analysis aims to provide researchers with practical guidance for optimizing genotyping strategies in genetic research.

## 2 Materials and Methods

### 2.1 Datasets and computational pipelines

An overview of the analytical pipeline is presented in Figure 1. In brief, we utilize both mapped sequences in CRAM format and phased genomic data in Variant Call Format (VCF) from 2,504 unrelated individuals in the 1000 Genomes Project dataset, which was re-sequenced at high coverage by the New York Genome Center (1KGPHC)^[31]^. To minimize noise during imputation and evaluation, we filter the VCF files to retain only biallelic SNPs with an allele count *≥* 2. We use the mapped sequences to simulate six LPS coverages (0.5, 0.75, 1.0, 1.25, 1.5, and 2.0x) and the SNP data to generate pseudo-arrays for eight different genotyping SNP chips. The obtained LPS and array data are then subjected to genotype imputation using a cross-validation approach, with performance evaluation performed by comparing the imputed data with the whole genome sequencing (WGS) 30x data (Fig 1).

**Figure 1:**
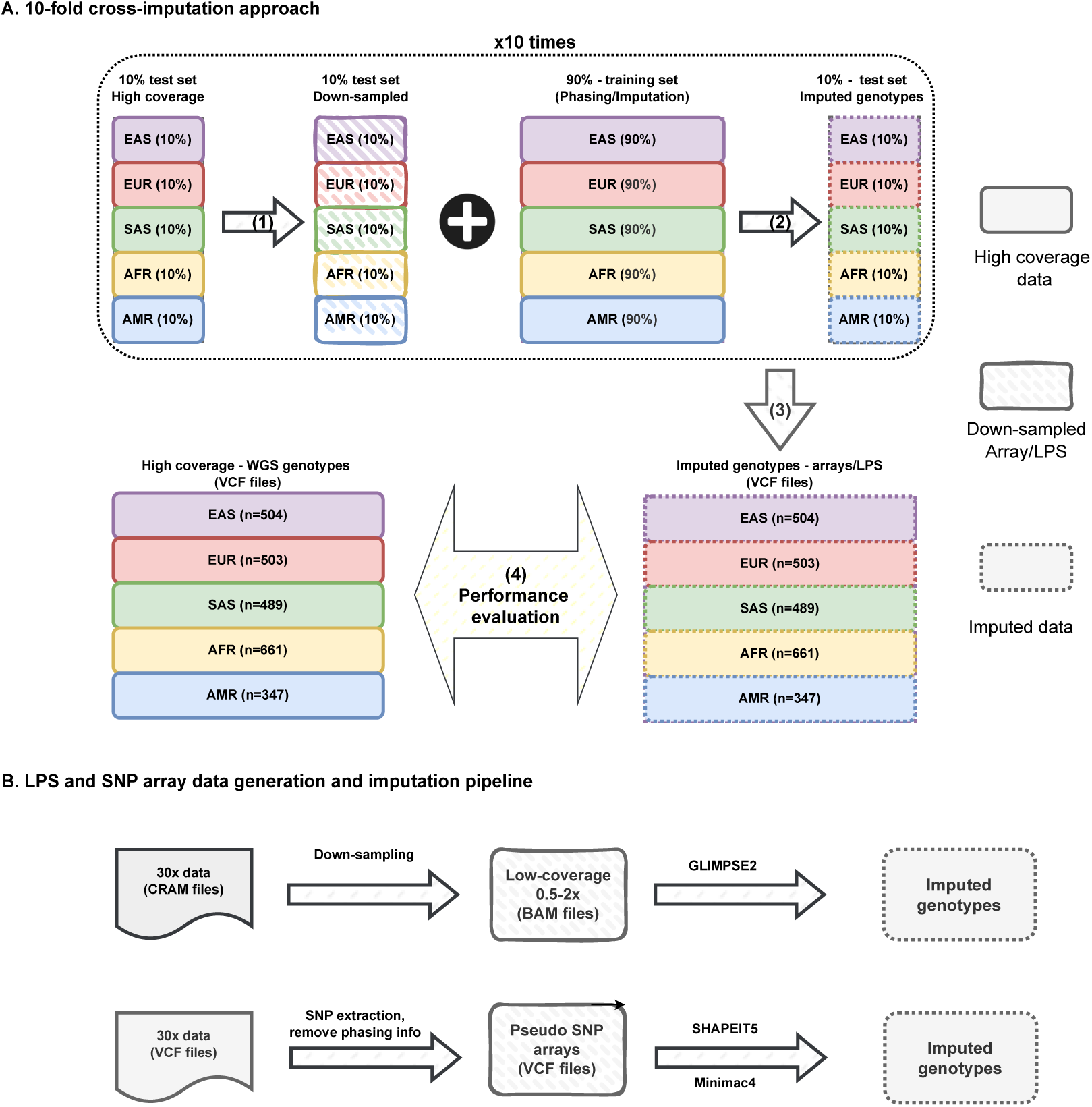
Overview of the analytical pipeline. A) 10-fold cross-imputation approach; (1) 10% of the samples are downsampled (BAM files) or filtered to retain only array variants (VCF files) to generate pseudo LPS and pseudo array data; (2) these data are imputed using the remaining 90% of the samples as the reference panel; (3) the imputed data from all batches are combined and then split by population; (4) performance is evaluated using high-coverage genotyping data as the ground truth. B) Data generation and imputation pipeline for LPS and SNP array data.

For LPS data simulation, we choose to sample high-coverage data from the mapped sequences rather than the raw sequencing reads, following approaches used in previous studies^[26,27,29]^. This decision helps to avoid the high computational cost associated with re-aligning large volumes of sequencing data. Since the mapped data has already undergone masking of duplicated reads, we firstly adjust the target coverage values for sampling by accounting for a 9% duplication rate, as reported in the original dataset^[31]^. To make simulation reflect real sequencing procedures, which often result in LPS coverage levels that deviate slightly from theoretical expectations^[28]^, we further incorporate variability in targeted coverage into our simulation. Specifically, we sample the coverage values from a normal distribution with a mean equal to the adjusted coverage and a standard deviation of 0.1. To avoid extreme cases, the sampling is constrained to the 0.1–0.9 quantile range of the distribution before performing down-sampling. Additionally, a minimum coverage threshold of 0.1x is applied as a pseudo-quality control step to exclude excessively low-coverage data. Of note, this approach is different from previous studies that typically sample mapped reads to match theoretical target coverage values^[26,27,29]^.

For SNP arrays, we select eight genotyping platforms based on marker density, population-specific optimization, and overall performance, as reported previously^[20]^. These arrays include the Axiom UK Biobank Array (820k markers), Axiom JAPONICA Array (667k markers), Axiom Precision Medicine Research Array (900k markers), Axiom Precision Medicine Diversity Array (901k markers), Infinium Global Screening Array v3.0 (648k markers), Infinium CytoSNP-850K v1.2 (2,364k markers), Infinium Omni2.5 v1.5 Array, and Infinium Omni5 v1.2 Array (4,245k markers). We use hg38-harmonized array manifests to extract typed variants with vcftools v0.1.17 ^[32]^ and remove phasing information to generate pseudo-SNP array data, following established methodologies^[20]^.

Once the simulated datasets for both LPS and SNP arrays are generated, we perform genotype imputation using a 10-fold cross-validation framework. To ensure balanced representation across continental groups, we stratify batch splitting so that samples are evenly distributed across five superpopulations: East Asian (EAS), European (EUR), South Asian (SAS), African (AFR), and American (AMR). This results in four batches with 251 samples and six batches with 250 samples each. In each iteration, nine batches (90% of samples) serve as the reference panel, while the remaining batch (10% of samples) is used as the target input in either LPS or SNP array data. For SNP arrays, we perform phasing using SHAPEIT5 ^[33]^, followed by imputation with Minimac4 ^[34]^. For LPS data, we conduct both phasing and imputation using GLIMPSE2^[35]^. Finally, imputed genotypes from all ten batches are merged by superpopulation for performance evaluation, as detailed in the following section.

### 2.2 Imputation and PGS performance assessment

Imputation increases the number of SNPs available for association testing, which is critical for GWAS, while for PGS, accurate imputation is essential since scores are derived by summing the product of risk allele counts (0, 1, or 2) and their estimated effect sizes. As a result, the accuracy of imputation is pivotal for statistical power^[36]^ and the reliability of PGS predictions. In line with previous studies^[20,37,28]^, we prioritize the use of the SNP-wise imputation *r*^2^ metric for several key reasons: (1) its direct relevance to both GWAS and PGS performance at individual variant levels^[38,39,40,28]^, (2) its ability to account for imputation uncertainty by utilizing expected allele dosages instead of relying solely on the most probable genotypes^[41]^, (3) its reduced sensitivity to allele frequency compared to concordance-based metrics^[41]^, making it better suited for assessing rare variants.

We use genotypes from WGS 30x datasets as the golden standard to evaluate imputation performance. Imputation accuracy is measured by calculating the SNP-wise squared Pearson correlation (*r*^2^) between the imputed dosages and WGS genotypes. For imputation coverage, a variant is defined to be covered if the *r*^2^ value for this variant is above or equal to 0.8. We focus on evaluating performance of SNPs with MAF belong to the range of (0.01–0.5] that is the commonly used threshold in GWAS and PGS analyses^[42,9]^. In addition, we also categorize these metrics into three MAF bins (0 *−* 0.01], (0.01 *−* 0.05], and (0.05 *−* 0.5] to assess imputation performance in various allele frequencies.

Regarding PGS performance assessment, we use the same method described previously^[20]^ that compute PGS scores using a standard P+T (Pruning and Thresholding) approach implemented in PRSice-2 ^[43]^. Compared to the use of pre-tuned PGS models^[28,44]^, this approach better reflects real-world PGS analysis, which involves testing multiple parameter settings to identify the optimal model^[9]^. Additionally, it avoids potential bias associated with pre-built PGS models, which may be over-fitted to specific arrays used during model training^[20]^.

Prior to PGS computation, we follow quality control procedures as recommended in the protocol for PGS analysis by Choi et al^[9]^. Using summary statistics from previous GWAS meta-analyses for body mass index (BMI), height^[45]^, type 2 diabetes^[46]^, and metabolic syndrome^[47]^, we calculate the PGS for each individual *i* as follows:

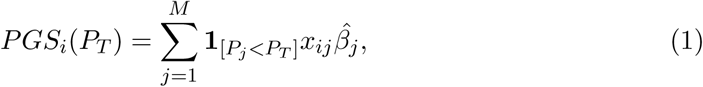

where *P_T_* represents various p-value threshold; *M* is the number of SNPs after clumping; *x_ij_* is the allele count, and 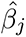 is the estimated effect size of *SNP_j_*.

## 3 Results

### 3.1 Imputation performance

We measure imputation performance using two metrics: (1) imputation accuracy, defined as the mean *r*^2^ of sites within the bin, and (2) imputation coverage, defined as the proportion of variants with *r*^2^ *≥* 0.8 over the total number of variants within the bin. These metrics are computed by chromosome basis for all autosome. The result visualizations are presented in Figure 2 and Figure 3.

**Figure 2:**
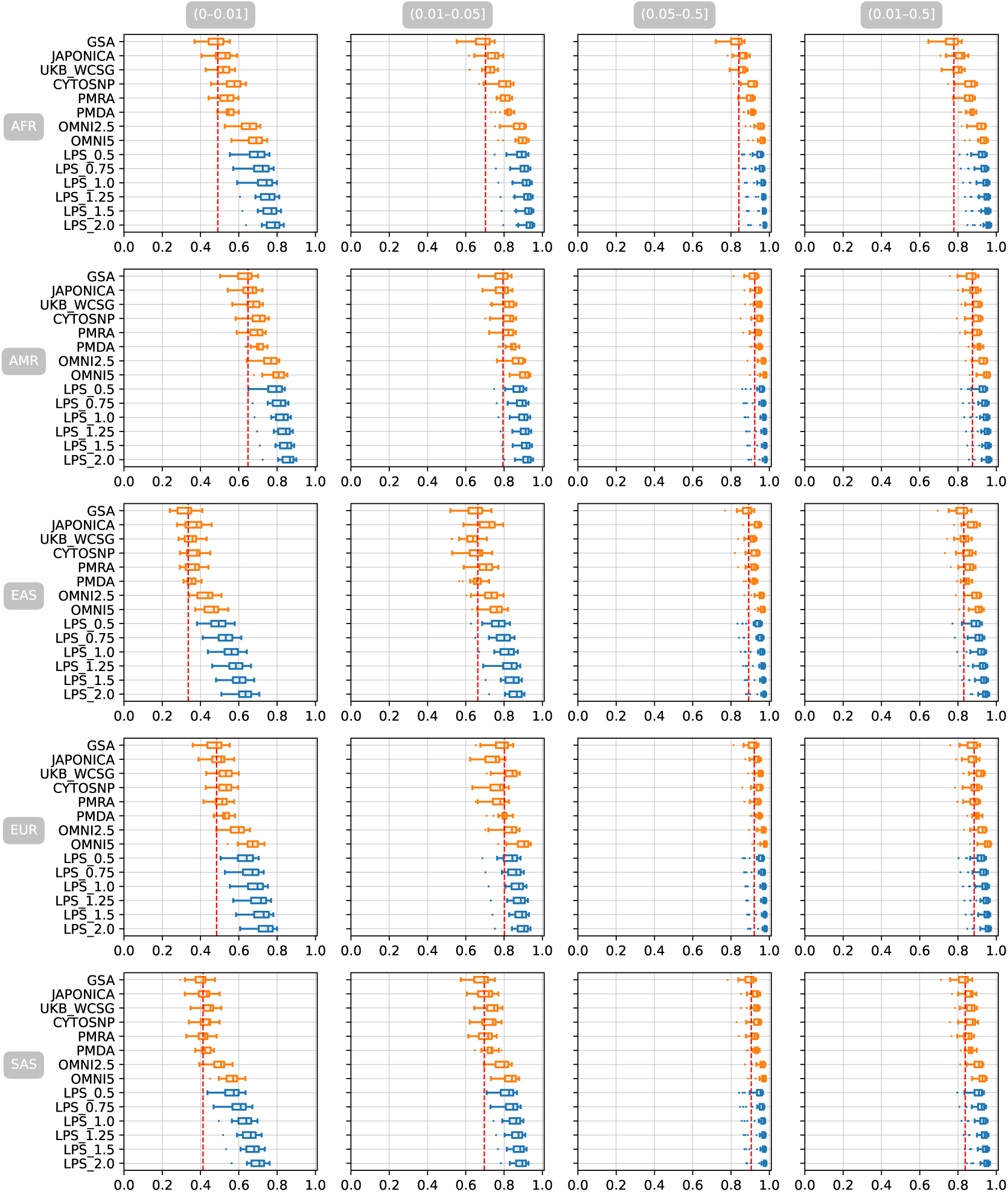
Imputation accuracy (mean *r*^2^) across 22 autosomes for eight genotyping arrays and six LPS coverages, evaluated across five populations for various MAF bins: (0 *−* 0.01], (0.01 *−* 0.05], (0.05 *−* 0.5], and (0.01 *−* 0.5]. The red dashed line represents the performance of the GSA array.

**Figure 3:**
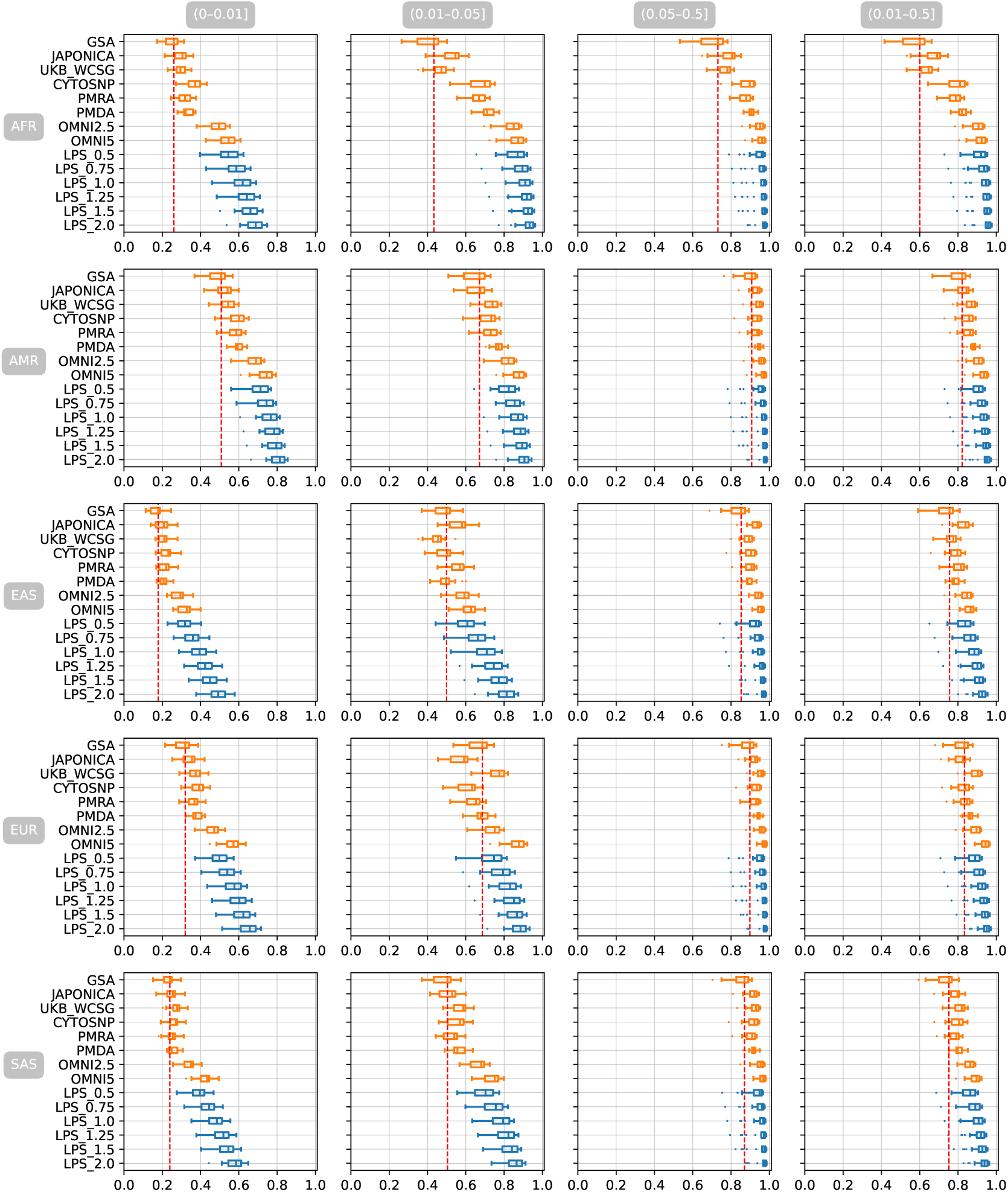
Imputation coverage across 22 autosomes for eight genotyping arrays and six LPS coverages, evaluated across five populations for various MAF bins: (0*−*0.01], (0.01*−*0.05], (0.05*−*0.5], and (0.01*−*0.5]. The red dashed line represents the performance of the GSA array.

In the most widely used MAF bin, (0.01 *−* 0.5], imputation performance strongly depends on array density and sequencing coverage, with additional variation attributable to population-specific optimizations for SNP arrays. As shown in Table 1 and Table 2, arrays such as GSA and sequencing at low coverage (e.g., LPS 0.5X) exhibit lower imputation accuracy across populations, while high-density arrays like OMNI5 and higher sequencing coverages (e.g., LPS 2.0X) consistently perform well across diverse populations. The performance of certain SNP arrays is also tailored to specific genetic backgrounds. For example, the UKB WCSG array achieves favorable imputation performance in the EUR population (0.912 for accuracy and 0.892 for coverage), but performs less effectively in other populations such as EAS (0.831 for accuracy and 0.764 for coverage) and SAS (0.865 for accuracy and 0.808 for coverage). Similarly, the JAPONICA array demonstrates superior performance in the EAS population (0.873 for accuracy and 0.826 for coverage), outperforming other arrays of similar density, such as GSA (0.816 for accuracy and 0.735 for coverage) and UKB WCSG (0.831 for accuracy and 0.764 for coverage). These results are expected due to the population-specific design of these arrays, with the UKB WCSG array optimized for European genetic variation through its development using cohorts such as the UK Biobank, while the JAPONICA array is tailored to capture Japanese genetic diversity that closely related to the EAS population. This observation is consistent with a previous evaluation of human genotyping arrays^[20]^. In contrast, the performance of LPS exhibits much less variation across populations. For example, at 0.5X coverage, imputation accuracy is 0.913, 0.918, 0.883, 0.909, and 0.897; and imputation coverage is 0.898, 0.890, 0.822, 0.872, and 0.848 for AFR, AMR, EAS, EUR, and SAS, respectively. Higher sequencing coverage improves imputation accuracy across all populations. For instance, LPS 2.0X achieves an average imputation accuracy of 0.946 in AFR populations compared to 0.913 for LPS 0.5X, highlighting the benefit of deeper sequencing. Similarly, in EUR populations, imputation accuracy increases from 0.909 at LPS 0.5X to 0.944 at LPS 2.0X. This trend is consistent across other populations, reinforcing the advantage of higher sequencing coverage in enhancing imputation accuracy.

**Table 1:**
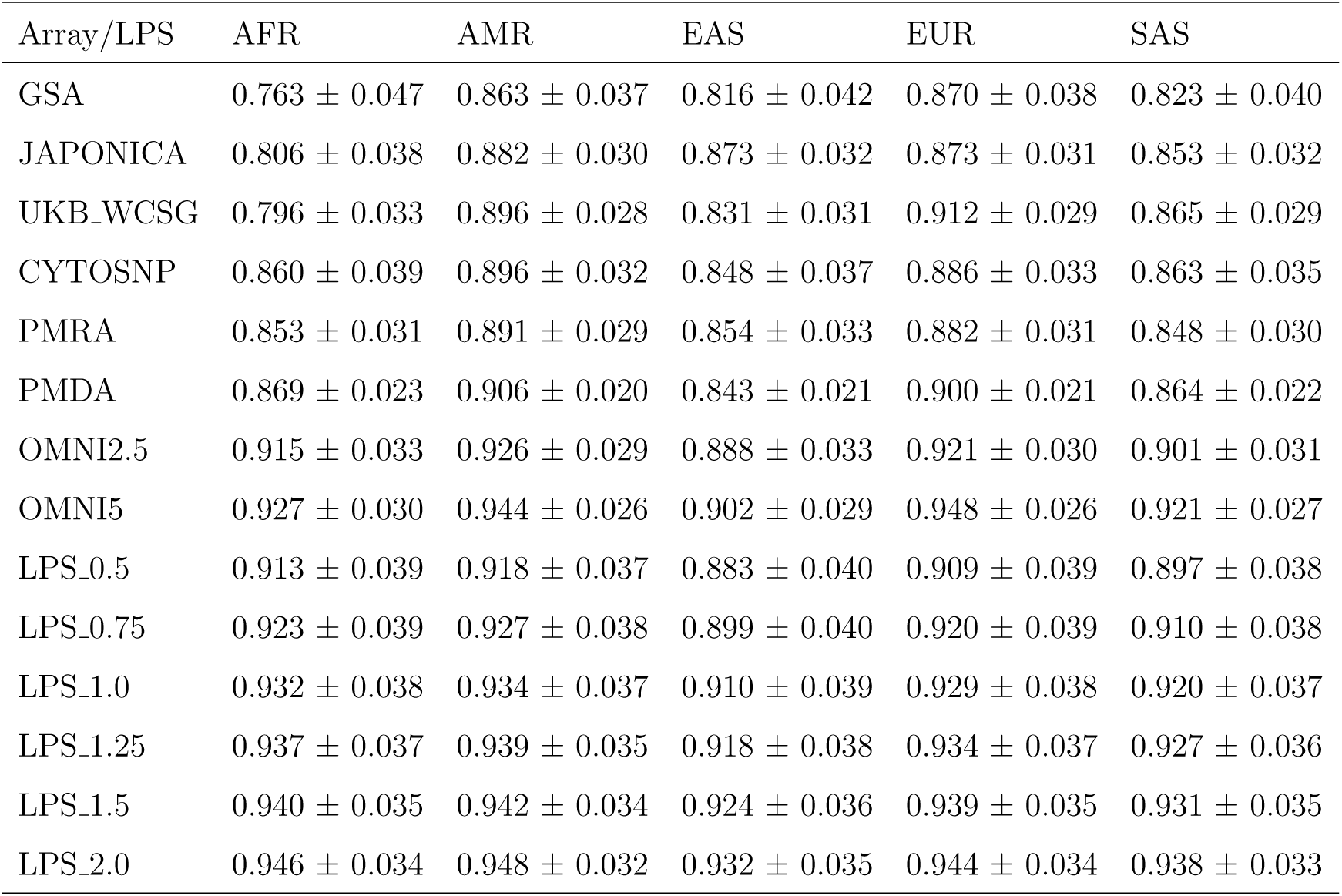
Imputation accuracy (mean and standard deviation across 22 autosomes) for eight genotyping arrays and six LPS coverages, evaluated across five populations for variant with allel frequency (0.01 *−* 0.5].

**Table 2:** Imputation coverage (mean and standard deviation across 22 autosomes) for eight genotyping arrays and six LPS coverages, evaluated across five populations for variant with allel frequency (0.01 *−* 0.5].

| Array/LPS | AFR | AMR | EAS | EUR | SAS |
| --- | --- | --- | --- | --- | --- |
| GSA | $0.567 \pm 0.076$ | $0.799 \pm 0.054$ | $0.735 \pm 0.053$ | $0.814 \pm 0.052$ | $0.731 \pm 0.054$ |
| JAPONICA | $0.668 \pm 0.062$ | $0.827 \pm 0.042$ | $0.826 \pm 0.039$ | $0.809 \pm 0.040$ | $0.780 \pm 0.041$ |
| UKB_WCSG | $0.633 \pm 0.046$ | $0.861 \pm 0.034$ | $0.764 \pm 0.035$ | $0.892 \pm 0.034$ | $0.808 \pm 0.034$ |
| CYTOSNP | $0.790 \pm 0.057$ | $0.848 \pm 0.040$ | $0.783 \pm 0.042$ | $0.828 \pm 0.040$ | $0.794 \pm 0.042$ |
| PMRA | $0.780 \pm 0.040$ | $0.850 \pm 0.035$ | $0.803 \pm 0.037$ | $0.834 \pm 0.037$ | $0.780 \pm 0.036$ |
| PMDA | $0.822 \pm 0.029$ | $0.879 \pm 0.022$ | $0.782 \pm 0.025$ | $0.862 \pm 0.024$ | $0.803 \pm 0.028$ |
| OMNI2.5 | $0.898 \pm 0.040$ | $0.900 \pm 0.034$ | $0.838 \pm 0.036$ | $0.887 \pm 0.035$ | $0.854 \pm 0.035$ |
| OMNI5 | $0.914 \pm 0.036$ | $0.929 \pm 0.030$ | $0.856 \pm 0.032$ | $0.936 \pm 0.031$ | $0.887 \pm 0.032$ |
| LPS_0.5 | $0.898 \pm 0.058$ | $0.890 \pm 0.055$ | $0.822 \pm 0.056$ | $0.872 \pm 0.056$ | $0.848 \pm 0.056$ |
| LPS_0.75 | $0.914 \pm 0.056$ | $0.905 \pm 0.054$ | $0.848 \pm 0.055$ | $0.892 \pm 0.055$ | $0.872 \pm 0.055$ |
| LPS_1.0 | $0.923 \pm 0.054$ | $0.915 \pm 0.053$ | $0.868 \pm 0.055$ | $0.907 \pm 0.054$ | $0.889 \pm 0.054$ |
| LPS_1.25 | $0.929 \pm 0.053$ | $0.922 \pm 0.051$ | $0.883 \pm 0.053$ | $0.916 \pm 0.052$ | $0.901 \pm 0.052$ |
| LPS_1.5 | $0.933 \pm 0.050$ | $0.928 \pm 0.048$ | $0.895 \pm 0.048$ | $0.924 \pm 0.048$ | $0.910 \pm 0.049$ |
| LPS_2.0 | $0.941 \pm 0.041$ | $0.937 \pm 0.041$ | $0.913 \pm 0.042$ | $0.935 \pm 0.040$ | $0.925 \pm 0.040$ |

In terms of performance stratified by MAF, LPS-based genotyping achieves significantly higher performance in both mean *r*^2^ and coverage for rare variants (MAF *<* 0.01) (Supplemental Tables S1,S4) and low MAF variants (MAF range (0.01 *−* 0.05]) (Supplemental Tables S2,S5) compared to SNP arrays. However, for common variants, both approaches exhibit high accuracy (Supplemental Tables S3,S6). The performance of LPS-based genotyping is also more stable across populations, with differences becoming negligible when an optimized array for each population is used.

For the rare variant bin, the GSA array consistently demonstrates low performance across five populations, with imputation accuracy ranging from 0.321 to 0.629 and imputation coverage ranging from 0.168 to 0.489 for EAS and AMR, respectively. In contrast, even at the lowest sequencing depth tested (0.5X), LPS-based genotyping achieves imputation accuracy between 0.492 and 0.785 and imputation coverage between 0.314 and 0.705 across the corresponding populations. This performance surpasses most tested arrays, except for OMNI5, which achieves slightly better performance, with imputation accuracy and coverage ranging from 0.461 to 0.800 and 0.319 to 0.734, respectively. LPS with higher coverage generally outperforms all tested arrays for both metrics, as summarized in Table S1 and Table S4.

A similar trend is observed in the low MAF bin ((0.01 *−* 0.05]), as shown in Table S2 and Table S5. LPS continues to outperform SNP arrays in both mean *r*^2^ and imputation coverage across all five populations, though higher sequencing depth is required to surpass all SNP arrays completely. For example, at 1X sequencing depth, LPS achieves mean *r*^2^ values ranging from 0.813 to 0.904 and imputation coverage ranging from 0.699 to 0.892. Notably, the OMNI5 array performs competitively, with mean *r*^2^ values between 0.754 and 0.900 and coverage values between 0.618 and 0.870, but LPS generally surpasses it at higher depths.

For common variants (MAF *>* 0.05), both genotyping approaches achieve very high performance, as detailed in Table S3 and Table S6. However, LPS performance is more stable across populations, consistently exceeding 0.9, whereas SNP array performance varies substantially across populations. Additionally, LPS-based genotyping exhibits lower standard deviation across chromosomes, indicating higher robustness. These results highlight the superiority of LPS for low MAF variants, particularly as sequencing depth increases, with substantial improvements in both accuracy and coverage compared to SNP arrays across all populations.

### 3.2 PGS performance

We evaluate PGS performance of LPS and genotyping arrays by adopting two metrics: (i) Pearson’s correlation between PGS derived from imputed SNP array data and PGS from WGS, referred to as PGS correlation, and (ii) the absolute difference in percentile ranking (ADPR) between PGS from array-imputed data and the gold standard WGS. These assessments are conducted across multiple p-value thresholds to ensure an unbiased comparison^[20]^. The evaluation is performed in four phenotypes including BMI, height, type 2 diabetes, and metabolic syndrome. Overall, PGS performance closely reflects imputation accuracy, with arrays and LPS designs that achieve better imputation results showing higher PGS correlation and lower ADPR than those with lower imputation quality.

#### 3.2.1 PGS correlation

The summary results of PGS correlations for four different phenotypes are presented in Figure 4 and Tables S.7-10. Overall, both genotyping arrays and LPS approaches demonstrate high PGS correlations across all populations and phenotypes, with most correlations ranging from 0.95 to 0.99.

**Figure 4:**
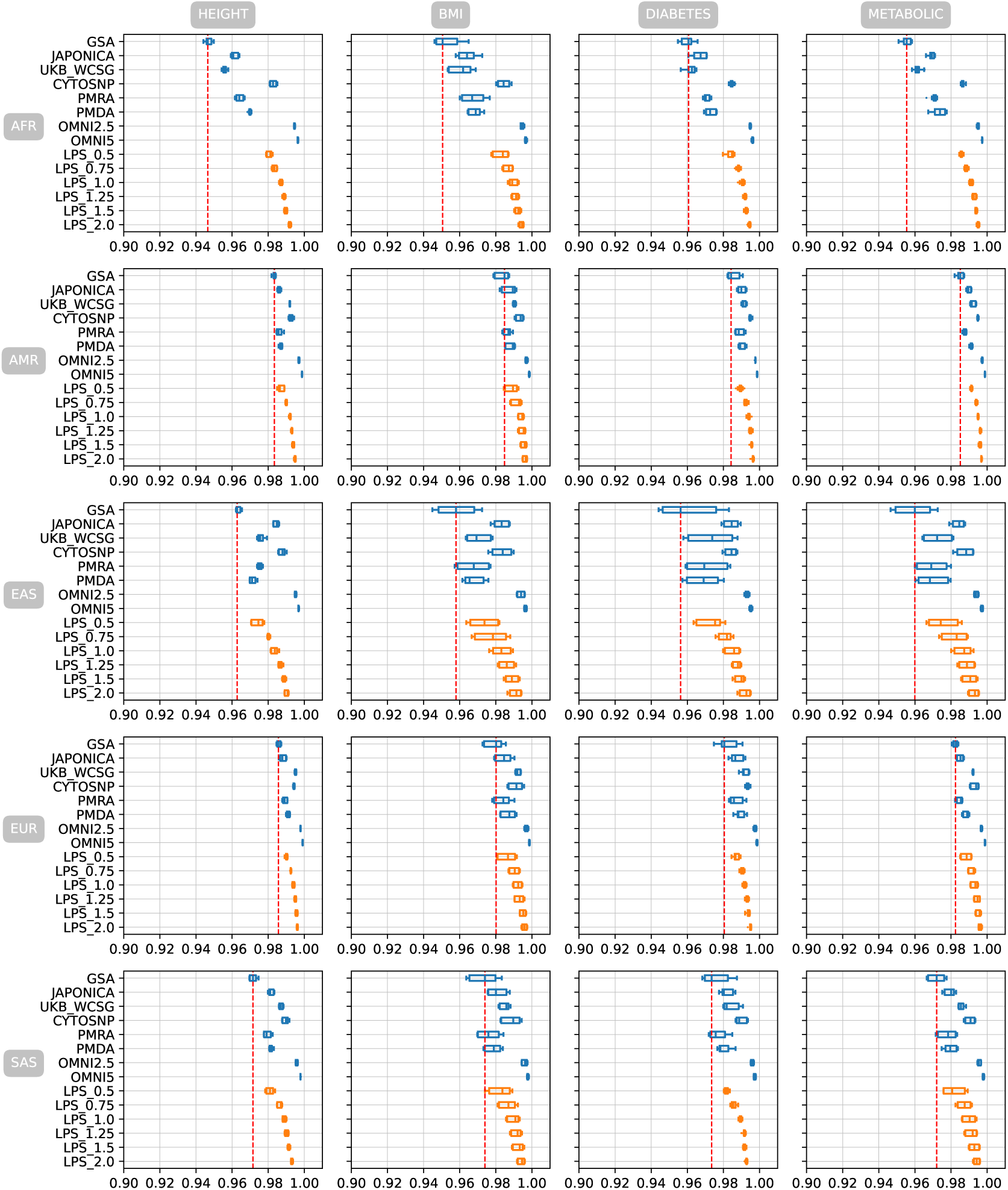
Correlations between PGS from imputed genotyping data (arrays and LPS) and PGS from WGS across five populations for four phenotypes (height, BMI, type 2 diabetes, and metabolic syndrome) at multiple PRSice p-value thresholds (5e-08 to 1). The red dashed line indicates the performance of the GSA array.

Among the genotyping arrays, dense arrays such as OMNI2.5 and OMNI5 consistently outperforms other arrays across all populations and traits. For instance, OMNI5 achieves correlations of 0.996 for height in AFR, 0.998 for BMI in AMR, 0.995 for diabetes in EAS, and 0.999 for metabolic traits in EUR. These results demonstrate the robustness and high performance of the high density arrays across diverse populations and complex traits. In contrast, GSA generally shows lower correlations due to its low density, particularly in the AFR population. For height, GSA achieves a correlation of 0.947 in AFR, which is the lowest among all arrays and populations. However, its performance improves in other populations, with correlations of 0.983 in AMR, 0.986 in EUR, and 0.972 in SAS populations for height.

Population-specific arrays demonstrate superior performance in their target populations. UKB WCSG excels in the EUR population, achieving correlations of 0.995 for height, 0.992 for BMI, 0.992 for diabetes, and 0.992 for metabolic traits. Similarly, JAPONICA performs well in the EAS population, with correlations of 0.984 for height, 0.983 for BMI, 0.984 for diabetes, and 0.984 for metabolic traits. In addition, the CYTOSNP array demonstrates remarkable consistency across all populations and traits regardless its moderate density. For example, in the height phenotype, CYTOSNP achieves correlations of 0.983 in AFR, 0.993 in AMR, 0.988 in EAS, 0.994 in EUR, and 0.990 in SAS populations. This consistency is maintained across other traits as well, suggesting CYTOSNP a reliable choice for PGS analysis in diverse population studies.

Regarding the LPS approaches, a clear trend of increasing PGS correlation with higher sequencing depth is observed across all populations and traits. LPS 2.0 consistently outperforms lower coverage options and serveral genotyping arrays. For instance, in the height phenotype, LPS 2.0 achieves correlations of 0.992 in AFR, 0.995 in AMR, 0.990 in EAS, 0.996 in EUR, and 0.993 in SAS populations.

Comparing LPS approaches to genotyping arrays, we observe that LPS 1.0 and higher obtain competitive well to most of the arrays, except for OMNI5 and OMNI2.5. For example, in the BMI trait for the AFR population, LPS 1.0 achieves a correlation of 0.990, which is higher than GSA (0.953), JAPONICA (0.964), and UKB WCSG (0.961),

but slightly lower than CYTOSNP (0.984) and OMNI5 (0.997). Interestingly, the relative performance of LPS approaches compared to arrays varies across traits and populations. In the EAS population, LPS options show particularly strong performance relative to arrays for the metabolic trait. LPS 1.0 achieves a correlation of 0.986, outperforming GSA (0.959), PMRA (0.969), and PMDA (0.970), and matching the performance of population-specific arrays like JAPONICA (0.984). In the AFR population, which often shows lower imputation accuracy due to higher genetic diversity, LPS approaches demonstrate notable improvements over some arrays. For the height trait, LPS 1.0 achieves a correlation of 0.987, outperforming GSA (0.947), JAPONICA (0.961), and UKB WCSG (0.956). This highlights the potential of LPS to capture population-specific variants that may be missed by some arrays, particularly in the context of diverse population or under-presentative populations.

#### 3.2.2 Absolute difference in percentile ranking

Regarding the ADPR metric, the performance of arrays and LPS aligns with the trends observed in PGS correlations, reflecting variations by array sizes, optimization populations, and LPS coverage. ADPR measurements across different PRSice-2 p-value settings are presented in Figure 5 and Figures S.1-11, with detailed results reported in Tables S.11-22. In general, ADPR values vary by traits and are slightly affected by cutoffs; however, most arrays and LPS coverages yield mean ADPRs around or below 5 across all four traits. Notably, ADPR values differ significantly between populations, with under-represented populations such as AFR and EAS exhibiting higher ADPRs compared to others.

**Figure 5:**
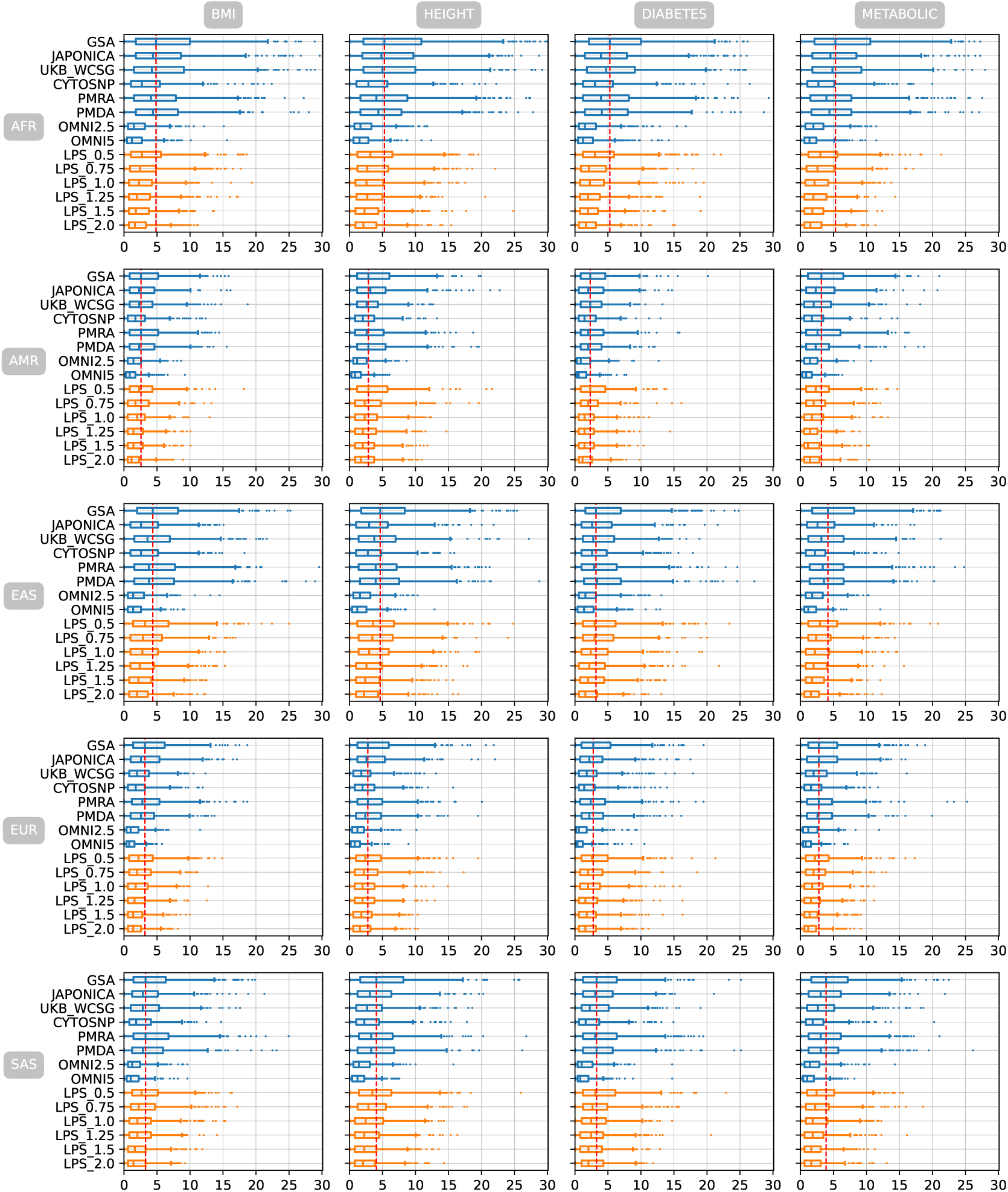
Absolute difference in percentile ranking between PGS from imputed genotyping data (arrays and LPS), and WGS across five populations for four phenotypes (height, BMI, type 2 diabetes, and metabolic syndrome) using a PRSice p-value threshold of 1e-5.

For instance, at a p-value cutoff of 1e-5 for the height phenotype (Figure 5), OMNI5 achieves ADPR means of 1.936, 1.268, 1.768, 1.043, and 1.580 in AFR, AMR, EAS, EUR, and SAS populations, respectively. A consistent trend is observed across other traits, where OMNI5 consistently demonstrates the lowest ADPR values among all arrays. For BMI, OMNI5 achieves ADPR means of 1.791 in AFR and 1.087 in EUR—the highest and lowest-performing populations for this trait, respectively. Similarly, for type 2 diabetes, OMNI5 achieves ADPR means of 1.928 in AFR and 0.929 in EUR. For metabolic traits, OMNI5 shows strong performance with ADPR means of 1.790 in AFR and 1.098 in EUR. Population-specific arrays also illustrate their advantages when comparing the ADPR metric. The UKB WCSG array performs particularly well for EUR populations with ADPR means of 2.365 for height, 2.564 for BMI, and 2.421 for type 2 diabetes. Similarly, the JAPONICA array demonstrates strong performance for EAS populations with ADPR means of 3.933 for height, 3.538 for BMI, and 3.647 for type 2 diabetes.

LPS approaches also show competitive or superior performance compared to arrays across multiple traits and populations as LPS coverage increases. For height at a p-value cutoff of 1e-5, LPS 0.5 achieves ADPR means of 4.398 in AFR and 3.348 in EUR; however, at higher coverage (LPS 2.0), these values decrease significantly to 2.858 in AFR and 2.161 in EUR—highlighting the improvement associated with increased sequencing depth. Similar trends are observed for BMI and type 2 diabetes traits with LPS approaches outperforming several arrays at higher coverage levels while maintaining competitive performance at lower coverage levels such as LPS 0.5 or LPS 0.75. For instance, for BMI in AFR populations, LPS 2.0 achieves an ADPR mean of 2.328 compared to GSA’s mean of 6.707 and JAPONICA’s mean of 5.857.

These results highlight the potential of LPS as a cost-effective alternative to genotyping arrays, particularly at coverages of 1.0x and above. The flexibility of LPS in capturing population-specific variants may contribute to its strong performance across diverse populations and traits, making it an attractive option for large-scale genetic studies and biobank initiatives, especially in populations with limited representation in existing genotyping arrays.

## 4 Discussion

This study provides a comprehensive evaluation of imputation accuracy and PGS performance across eight genotyping arrays and six LPS coverage levels in diverse populations. Our findings demonstrate that LPS, particularly at coverage levels of 1x or higher, offers significant advantages over traditional SNP arrays in capturing rare variants and maintaining consistent performance across diverse populations. One key observation is the superior imputation accuracy of LPS compared to most genotyping arrays, particularly for rare (MAF *<* 0.01) and low-frequency variants (0.01 *<* MAF *<* 0.05). Rare variants are often omitted from SNP array designs due to their low minor allele frequency, limiting their utility in genetic studies focused on complex traits or diseases with rare variant contributions. LPS addresses this limitation by providing broader genomic coverage, enabling more accurate imputation of rare variants across all studied populations.

The consistency of LPS performance across diverse populations is another notable advantage. While population-specific arrays such as UKB WCSG and JAPONICA excel within their target populations (EUR and EAS, respectively), their performance diminishes in other populations due to differences in genetic architecture^[20]^. In contrast, LPS exhibits less variation across populations, making it an attractive option for multi-ethnic studies where inclusivity is paramount. This consistency is particularly evident in underrepresented populations such as AFR, where higher genetic diversity often reduces the effectiveness of SNP arrays.

The evaluation of PGS performance further supports the competitiveness of LPS compared to genotyping arrays that align with previous studies^[26,28,29]^. Using both Pearson correlation with WGS-derived PGS and absolute difference in percentile ranking metrics, we observed that LPS achieves comparable or superior PGS performance at 1x coverage or higher across diverse traits such as BMI, height, type 2 diabetes, and metabolic syndrome. Notably, the advantage of LPS becomes more pronounced as sequencing depth increases, with 2x coverage consistently outperforming most tested arrays.

Despite these advantages, high-density arrays such as OMNI5 remain competitive for certain applications due to their high marker density and optimized designs for common variants. For researchers prioritizing on well-represented populations with established array designs, these arrays may still be suitable alternatives.

A limitation of this study is the use of simulated LPS data derived from high-coverage sequencing rather than real-world LPS datasets. While this approach enable the possible of large scale evaluation, it may not account for possible differences between sequencing protocol and platforms^[28]^. Additionally, expanding the analysis to include additional traits beyond those evaluated here would provide deeper insights into the relative merits of LPS versus SNP arrays.

In summary, our results highlight the flexibility and scalability of LPS as a genotyping approach for large-scale genetic studies. By achieving superior imputation accuracy for rare variants and maintaining consistent performance across diverse populations, LPS addresses key limitations associated with array-based genotyping approaches. Furthermore, its competitive PGS performance across multiple traits underscores its potential utility in both research and clinical applications. As sequencing costs continue to decline, LPS is poised to become an indispensable tool for genetic studies aiming to improve population inclusivity and uncover novel genetic associations.

## Supplemental files

• Supplemental figures

• Supplemental tables

## Availability of data and materials

Source codes, relevant data and detailed instructions to reproduce the study are available at https://github.com/KTest-VN/lps_paper and https://ktest-vn.github.io/lps_paper/.

## Authors’ contributions

DTN conceived the study. DTN, TVN, and TDHH designed the study. PTN and DTN implemented computational pipelines and analyzed the data. PTN, DTN wrote the manuscript. DTN, TVN and TDHH supervise the study and contributed to discussion and manuscript revision. All authors read and approved the final manuscript.

## Use of AI Software

Large language models were used to improve the wording and grammar of some texts, but not to generate new content.

## Acknowledgments

This work was supported by internal funding from KTest Vietnam.

