## Supplemental figures for "Imputation and polygenic score performance of low coverage whole-genome sequencing and genotyping arrays in diverse human populations"

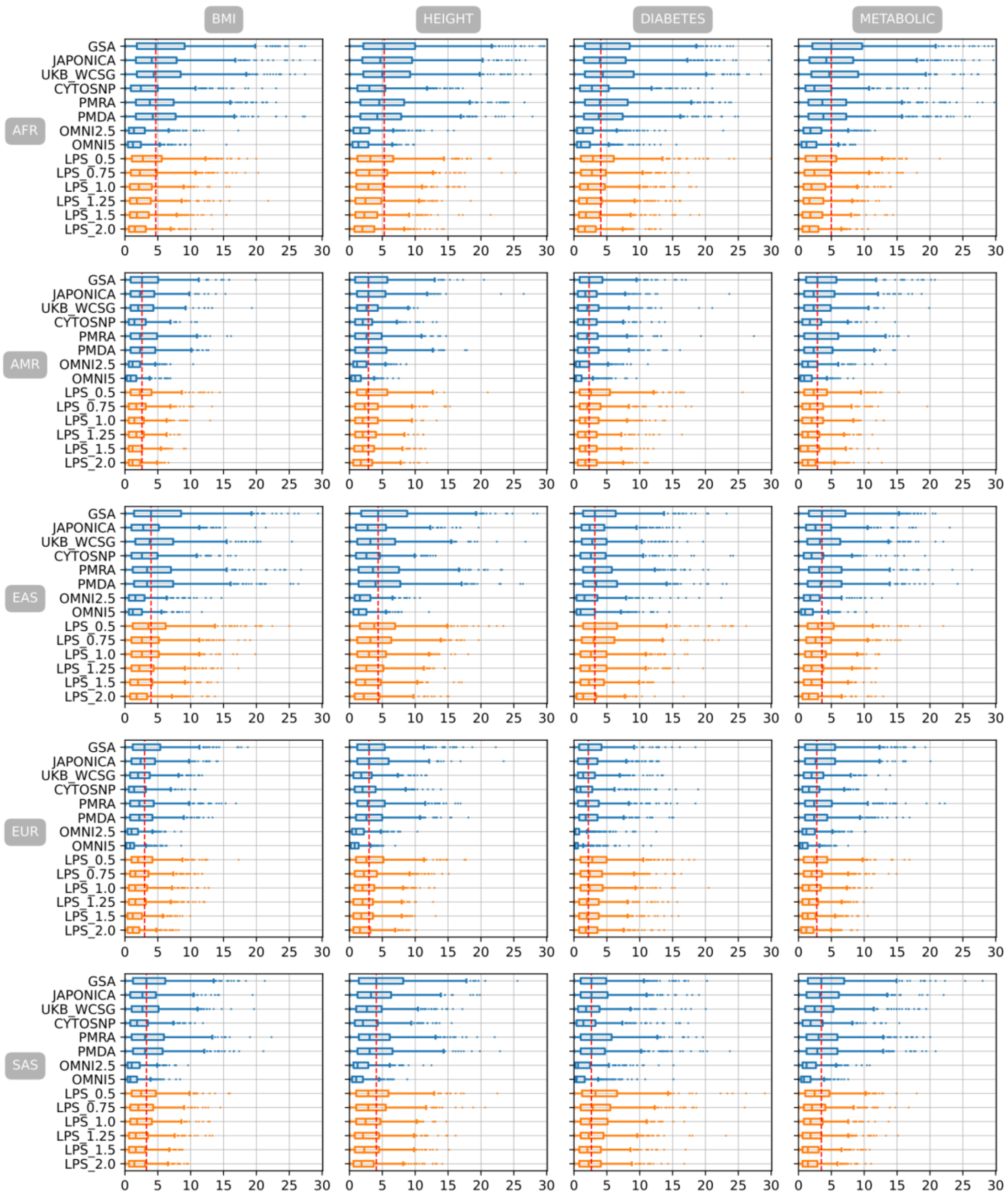


Figure S. 1 Mean absolute difference of percentile ranking between PGSs estimated from imputed genotyping data of eight genotyping arrays and six LPS coverages and PGS estimated from WGS in 5 different populations with PRsice p-value setting of 5e-08

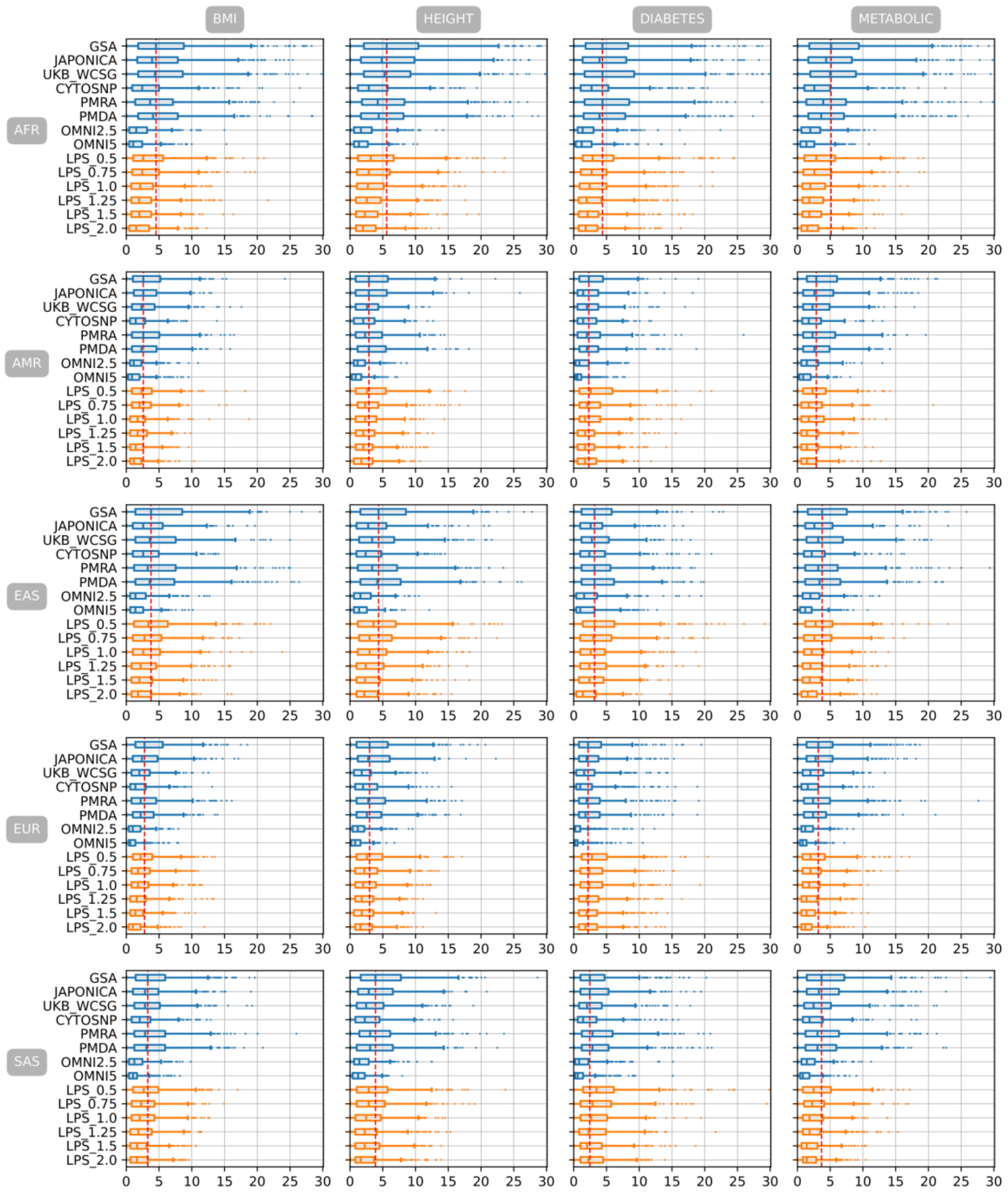


Figure S. 2 Mean absolute difference of percentile ranking between PGSs estimated from imputed genotyping data of eight genotyping arrays and six LPS coverages and PGS estimated from WGS in 5 different populations with PRsice p-value setting of 1e-07

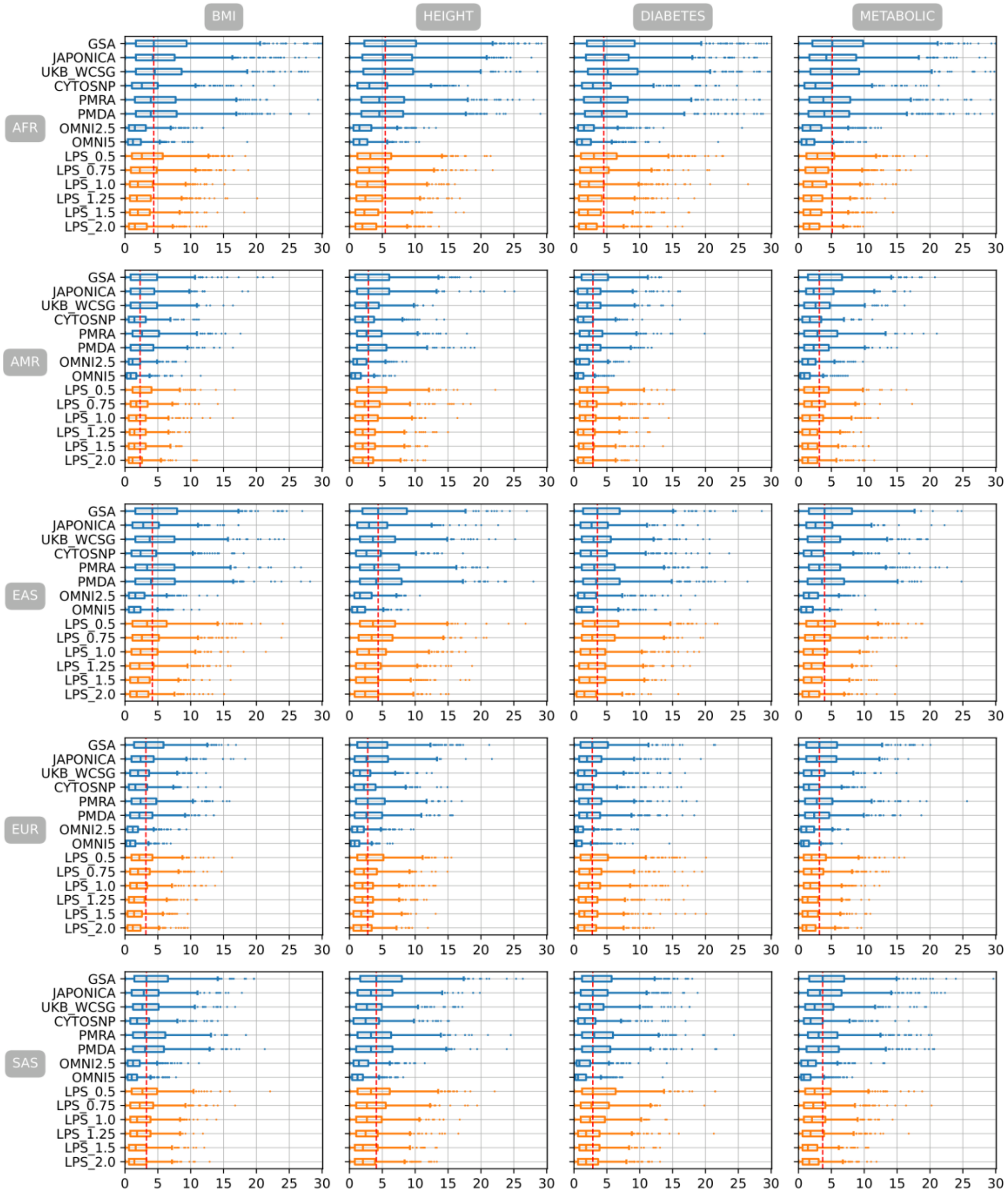


Figure S. 3 Mean absolute difference of percentile ranking between PGSs estimated from imputed genotyping data of eight genotyping arrays and six LPS coverages and PGS estimated from WGS in 5 different populations with PRsice p-value setting of 1e-06

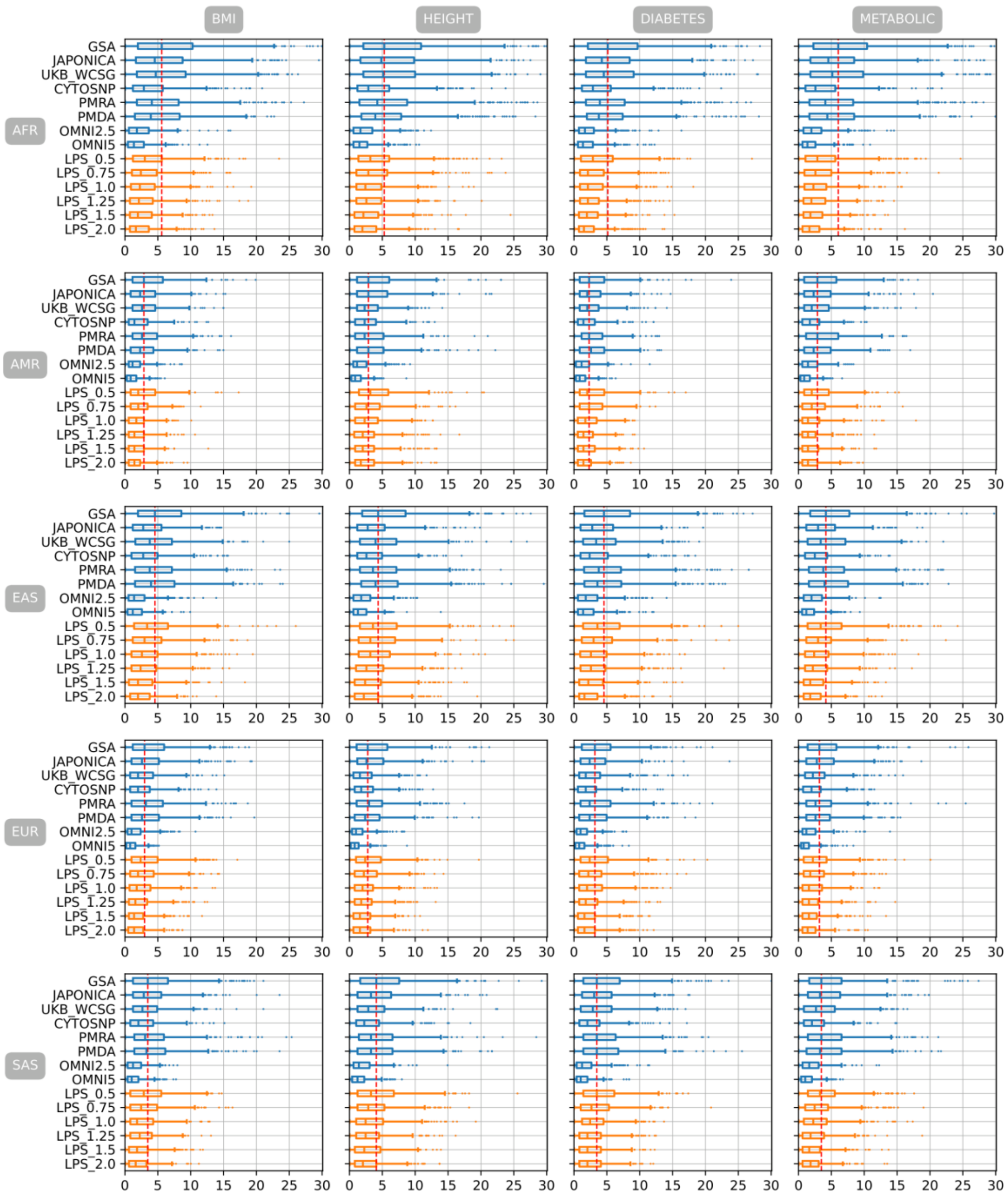


Figure S. 4 Mean absolute difference of percentile ranking between PGSs estimated from imputed genotyping data of eight genotyping arrays and six LPS coverages and PGS estimated from WGS in 5 different populations with PRsice p-value setting of 0.0001

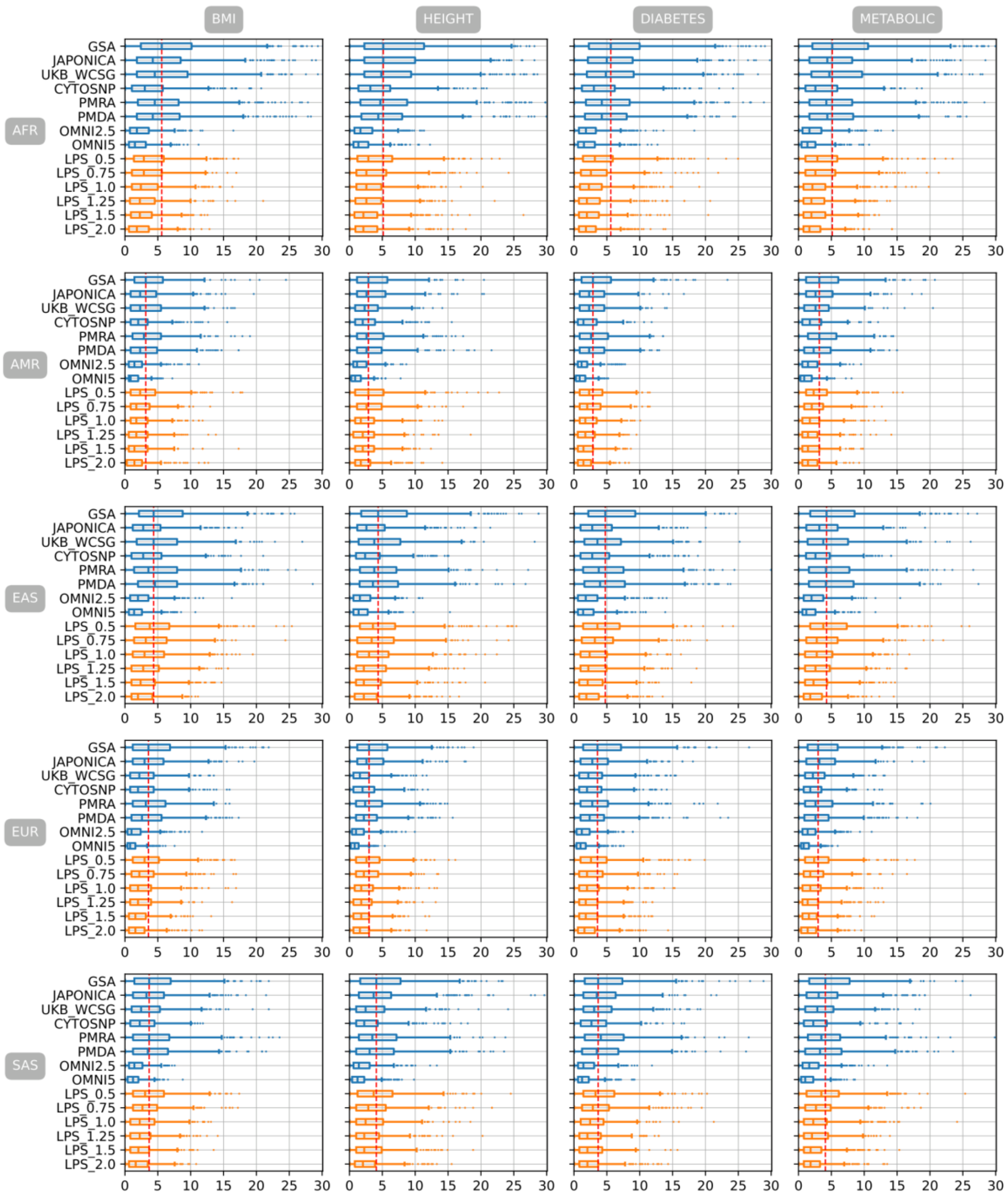


Figure S. 5 Mean absolute difference of percentile ranking between PGSs estimated from imputed genotyping data of eight genotyping arrays and six LPS coverages and PGS estimated from WGS in 5 different populations with PRsice p-value setting of 0.001

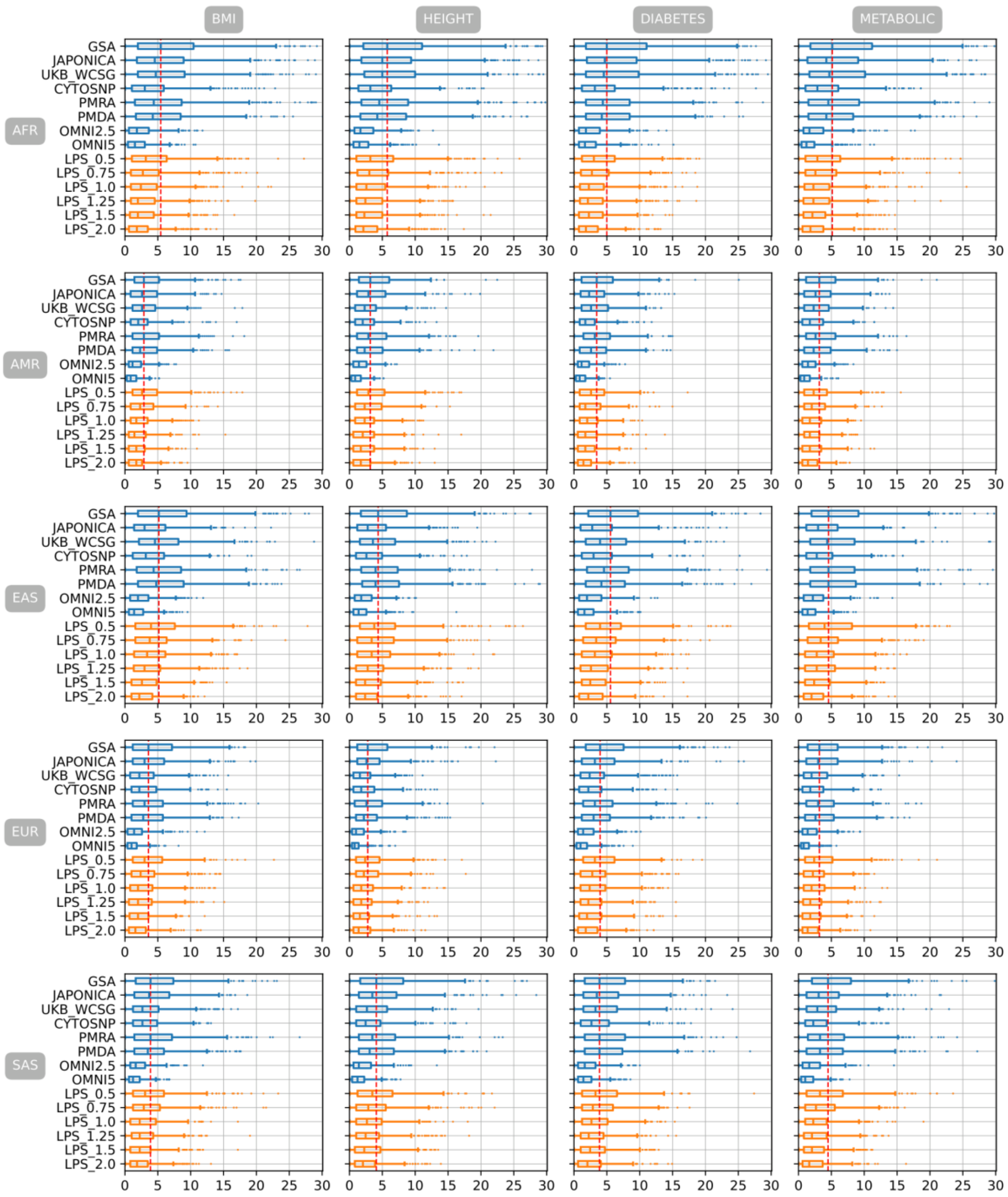


Figure S. 6 Mean absolute difference of percentile ranking between PGSs estimated from imputed genotyping data of eight genotyping arrays and six LPS coverages and PGS estimated from WGS in 5 different populations with PRsice p-value setting of 0.01

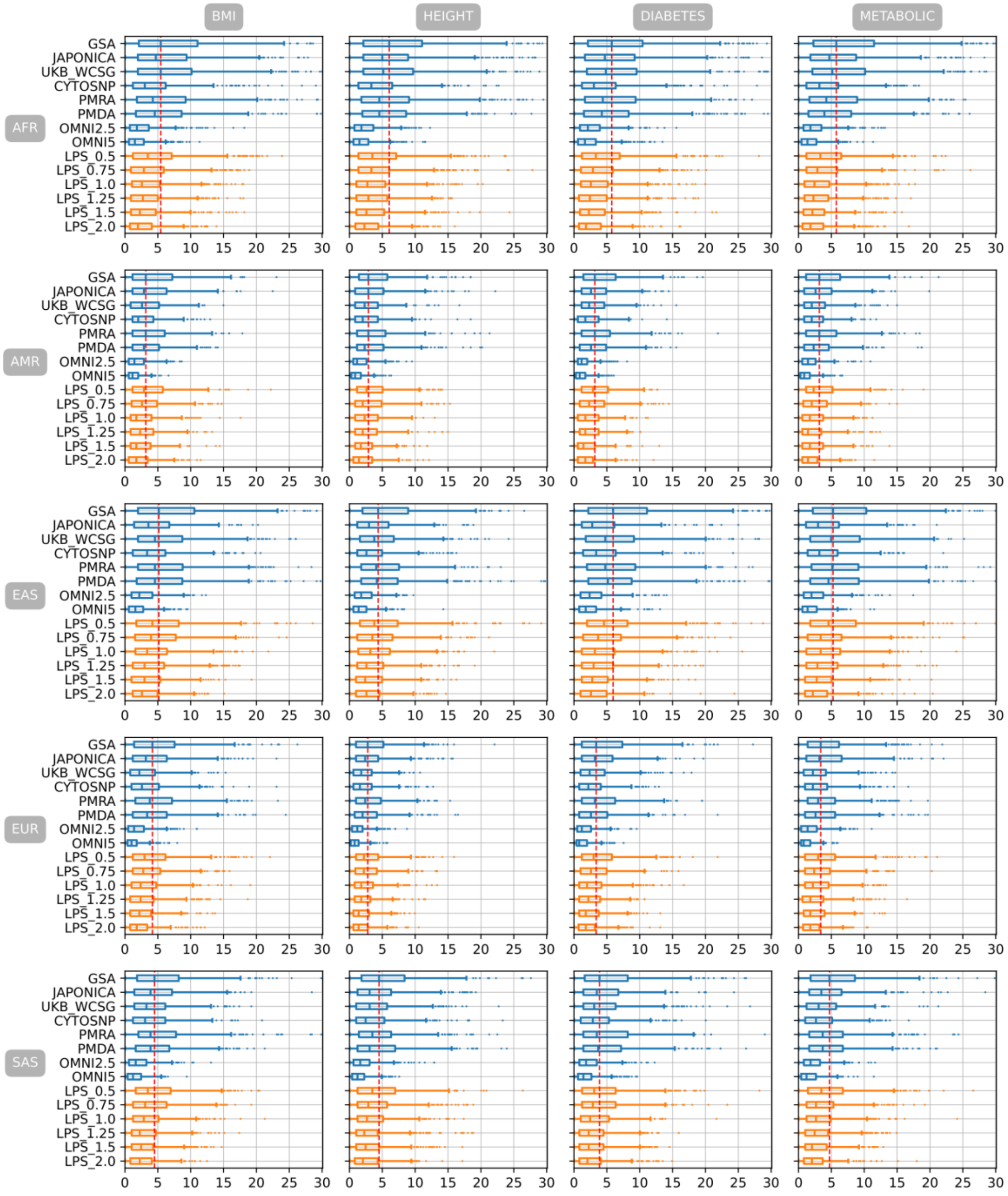


Figure S. 7 Mean absolute difference of percentile ranking between PGSs estimated from imputed genotyping data of eight genotyping arrays and six LPS coverages and PGS estimated from WGS in 5 different populations with PRsice p-value setting of 0.1

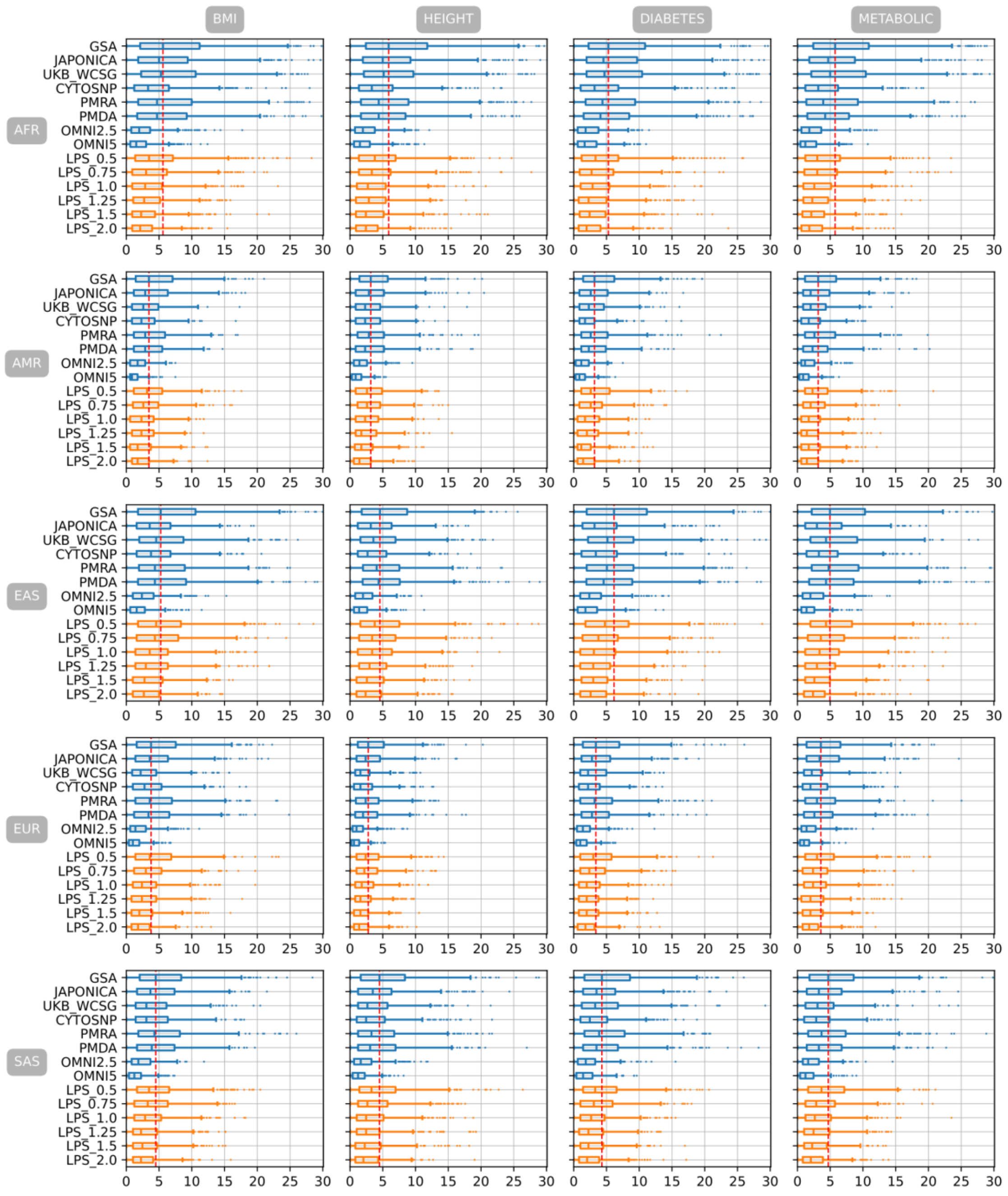


Figure S. 8 Mean absolute difference of percentile ranking between PGSs estimated from imputed genotyping data of eight genotyping arrays and six LPS coverages and PGS estimated from WGS in 5 different populations with PRsice p-value setting of 0.2

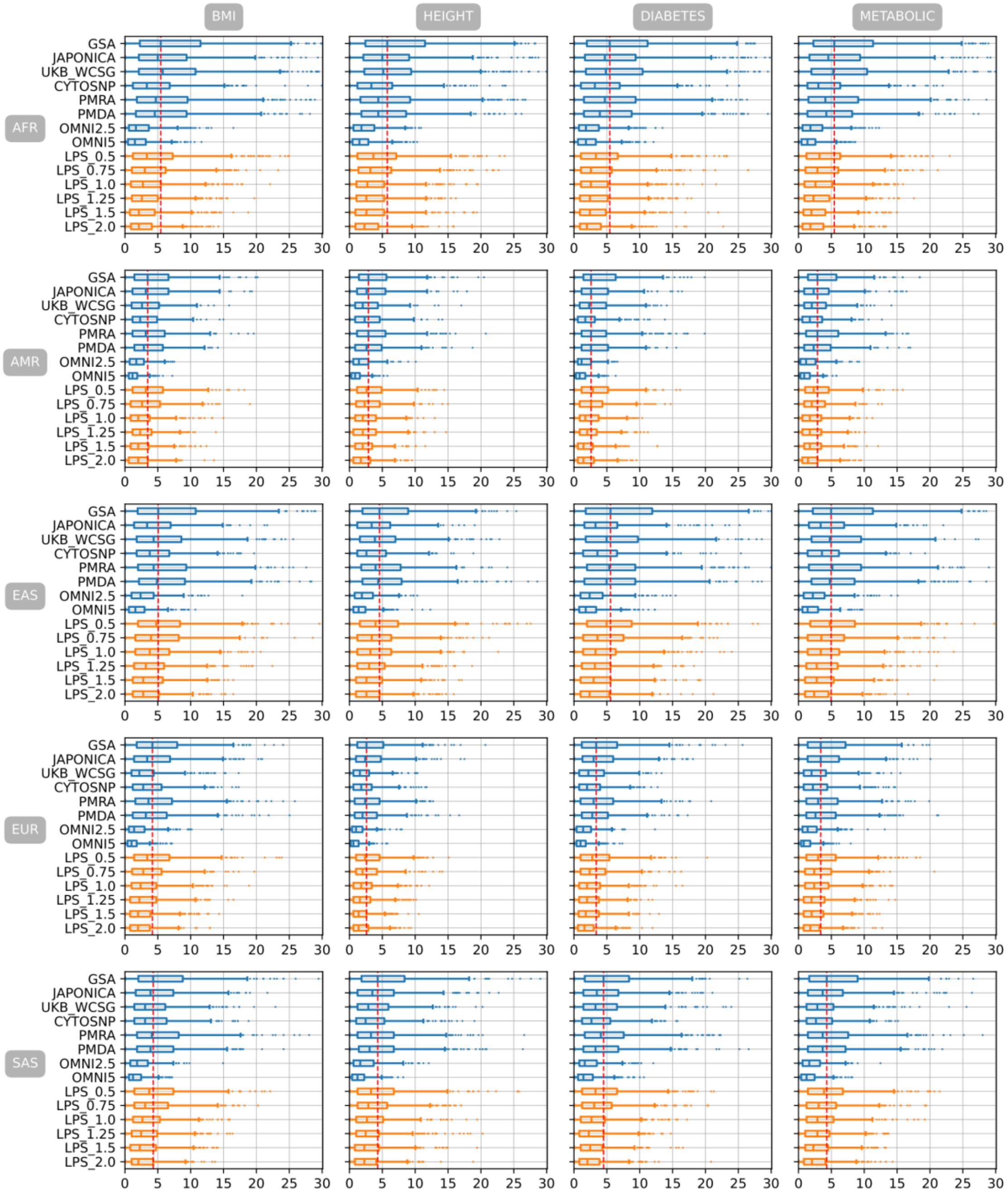


Figure S. 9 Mean absolute difference of percentile ranking between PGSs estimated from imputed genotyping data of eight genotyping arrays and six LPS coverages and PGS estimated from WGS in 5 different populations with PRsice p-value setting of 0.3

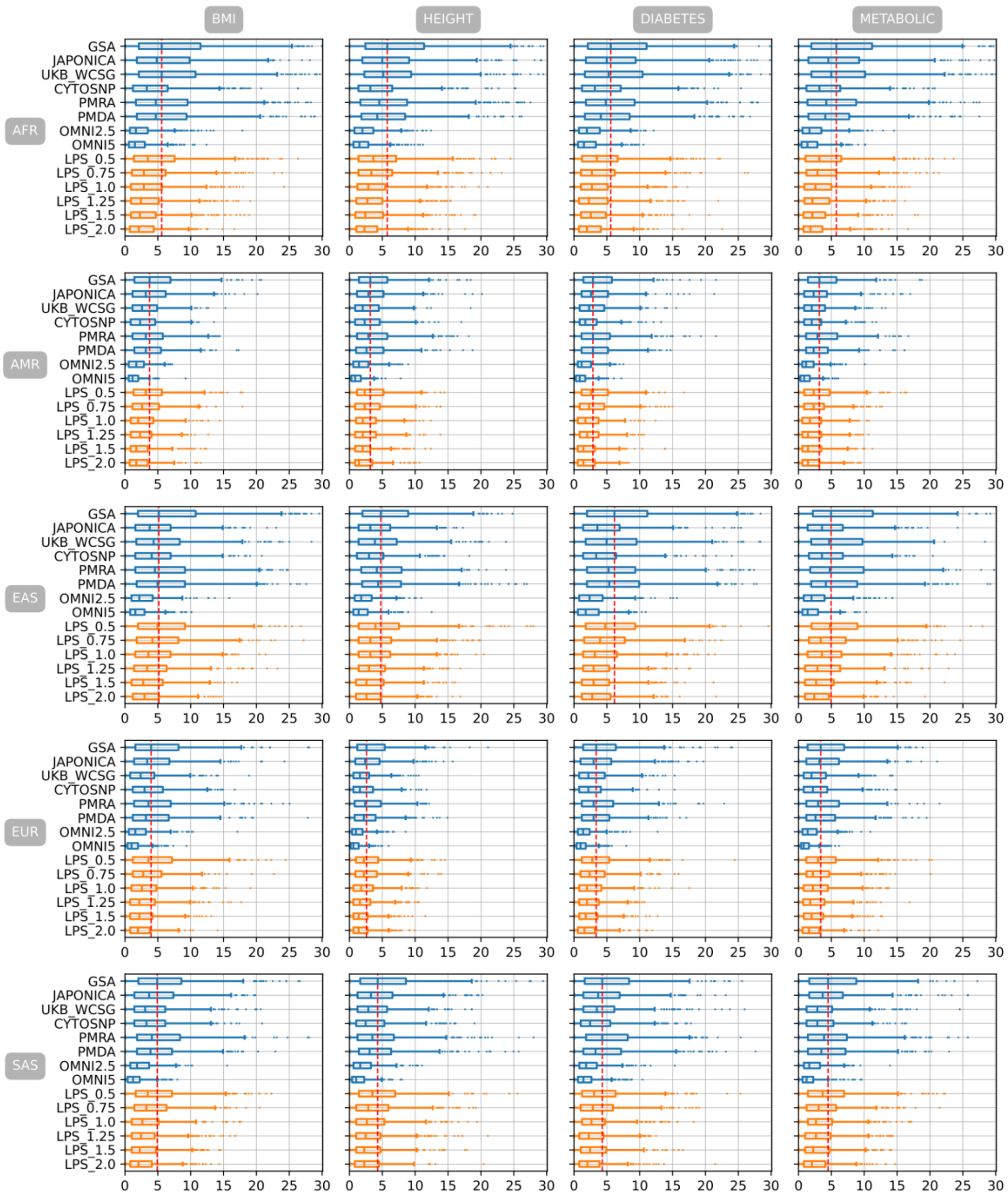


Figure S. 10 Mean absolute difference of percentile ranking between PGSs estimated from imputed genotyping data of eight genotyping arrays and six LPS coverages and PGS estimated from WGS in 5 different populations with PRsice p-value setting of 0.5

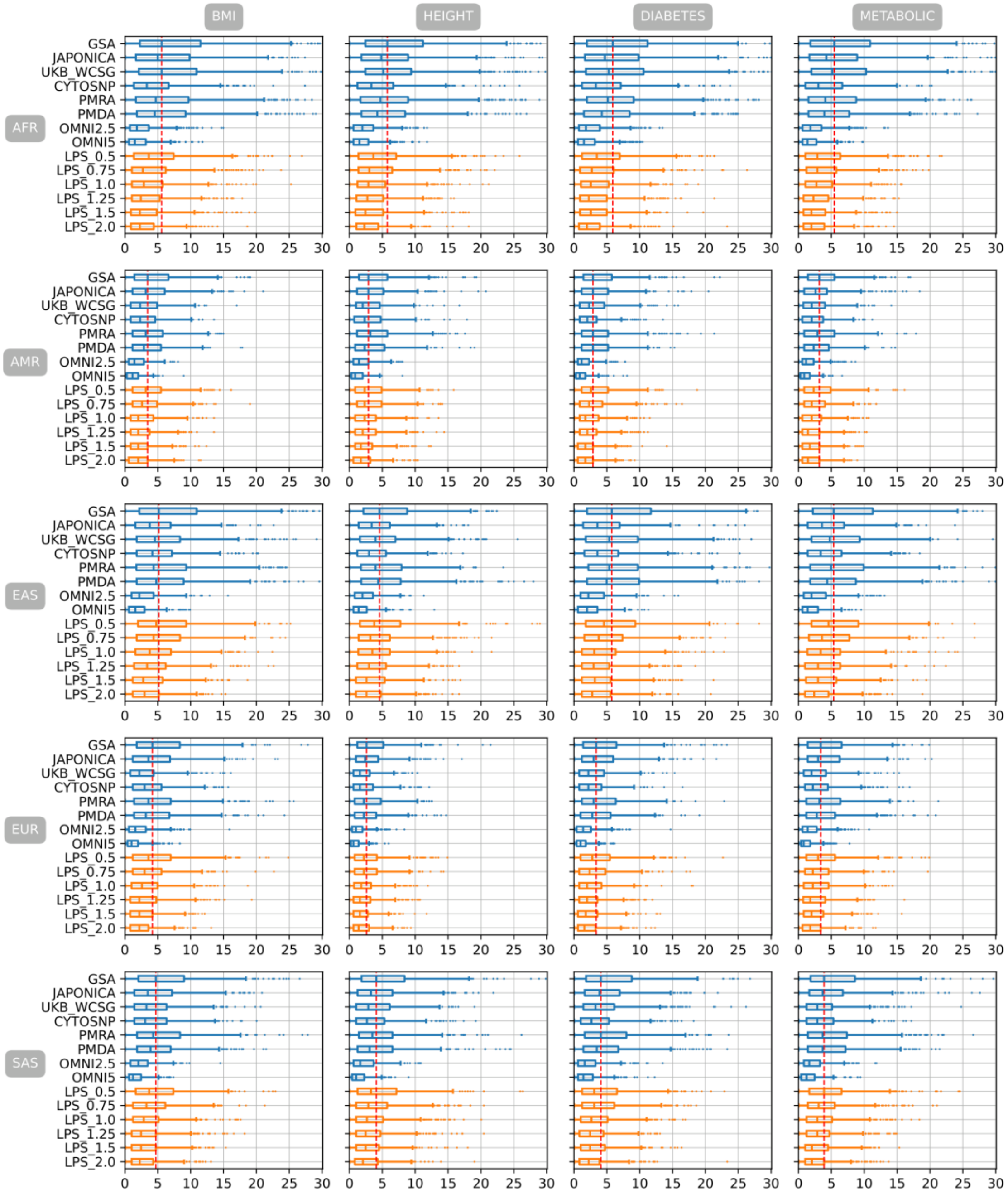


Figure S. 11 Mean absolute difference of percentile ranking between PGSs estimated from imputed genotyping data of eight genotyping arrays and six LPS coverages and PGS estimated from WGS in 5 different populations with PRsice p-value setting of 1
