## Supplemental tables for "Imputation and polygenic score performance of low coverage whole-genome sequencing and genotyping arrays in diverse human populations"

Table S. 1 Imputation accuracy (mean and standard deviation across 22 autosomes) for eight genotyping arrays and six LPS coverages, evaluated across five populations for variant with allel frequency (0–0.01]

| Array/LPS | AFR | AMR | EAS | EUR | SAS |
|---|---|---|---|---|---|
| GSA | 0.478 ± 0.051 | 0.629 ± 0.053 | 0.321 ± 0.046 | 0.471 ± 0.052 | 0.400 ± 0.045 |
| JAPONICA | 0.518 ± 0.048 | 0.658 ± 0.047 | 0.368 ± 0.048 | 0.497 ± 0.048 | 0.423 ± 0.044 |
| UKB_WCSG | 0.517 ± 0.040 | 0.669 ± 0.041 | 0.353 ± 0.038 | 0.528 ± 0.044 | 0.443 ± 0.039 |
| CYTOSNP | 0.567 ± 0.048 | 0.698 ± 0.045 | 0.366 ± 0.043 | 0.526 ± 0.044 | 0.428 ± 0.039 |
| PMRA | 0.536 ± 0.042 | 0.689 ± 0.041 | 0.364 ± 0.041 | 0.509 ± 0.042 | 0.417 ± 0.040 |
| PMDA | 0.551 ± 0.031 | 0.705 ± 0.030 | 0.351 ± 0.027 | 0.528 ± 0.031 | 0.425 ± 0.029 |
| OMNI2.5 | 0.648 ± 0.048 | 0.760 ± 0.044 | 0.429 ± 0.045 | 0.592 ± 0.046 | 0.499 ± 0.043 |
| OMNI5 | 0.682 ± 0.046 | 0.800 ± 0.044 | 0.461 ± 0.045 | 0.664 ± 0.047 | 0.564 ± 0.044 |
| LPS_0.5 | 0.691 ± 0.051 | 0.785 ± 0.047 | 0.492 ± 0.049 | 0.633 ± 0.050 | 0.562 ± 0.048 |
| LPS_0.75 | 0.715 ± 0.051 | 0.806 ± 0.047 | 0.528 ± 0.050 | 0.661 ± 0.051 | 0.598 ± 0.049 |
| LPS_1.0 | 0.734 ± 0.050 | 0.821 ± 0.046 | 0.558 ± 0.050 | 0.686 ± 0.050 | 0.627 ± 0.048 |
| LPS_1.25 | 0.748 ± 0.049 | 0.832 ± 0.045 | 0.581 ± 0.049 | 0.703 ± 0.049 | 0.650 ± 0.047 |
| LPS_1.5 | 0.759 ± 0.048 | 0.841 ± 0.044 | 0.599 ± 0.048 | 0.717 ± 0.048 | 0.668 ± 0.047 |
| LPS_2.0 | 0.776 ± 0.046 | 0.854 ± 0.042 | 0.629 ± 0.047 | 0.739 ± 0.047 | 0.696 ± 0.046 |

Table S. 2 Imputation accuracy (mean and standard deviation across 22 autosomes) for eight genotyping arrays and six LPS coverages, evaluated across five populations for variant with allele frequency (0.01–0.05]

| Array/LPS | AFR | AMR | EAS | EUR | SAS |
|---|---|---|---|---|---|
| GSA | 0.683 ± 0.056 | 0.781 ± 0.048 | 0.646 ± 0.057 | 0.782 ± 0.052 | 0.677 ± 0.052 |
| JAPONICA | 0.736 ± 0.048 | 0.788 ± 0.043 | 0.711 ± 0.054 | 0.738 ± 0.050 | 0.700 ± 0.048 |
| UKB_WCSG | 0.720 ± 0.040 | 0.820 ± 0.038 | 0.630 ± 0.047 | 0.830 ± 0.047 | 0.734 ± 0.040 |
| CYTOSNP | 0.797 ± 0.048 | 0.816 ± 0.043 | 0.653 ± 0.052 | 0.759 ± 0.051 | 0.720 ± 0.046 |
| PMRA | 0.797 ± 0.039 | 0.817 ± 0.038 | 0.699 ± 0.050 | 0.766 ± 0.049 | 0.703 ± 0.042 |
| PMDA | 0.818 ± 0.030 | 0.842 ± 0.028 | 0.656 ± 0.037 | 0.798 ± 0.033 | 0.729 ± 0.032 |
| OMNI2.5 | 0.872 ± 0.042 | 0.868 ± 0.039 | 0.726 ± 0.050 | 0.826 ± 0.049 | 0.787 ± 0.043 |
| OMNI5 | 0.887 ± 0.040 | 0.900 ± 0.036 | 0.754 ± 0.047 | 0.894 ± 0.043 | 0.828 ± 0.040 |
| LPS_0.5 | 0.881 ± 0.045 | 0.869 ± 0.044 | 0.763 ± 0.051 | 0.829 ± 0.049 | 0.812 ± 0.044 |
| LPS_0.75 | 0.894 ± 0.045 | 0.883 ± 0.043 | 0.791 ± 0.050 | 0.849 ± 0.049 | 0.834 ± 0.043 |
| LPS_1.0 | 0.904 ± 0.044 | 0.893 ± 0.042 | 0.813 ± 0.050 | 0.864 ± 0.047 | 0.851 ± 0.042 |
| LPS_1.25 | 0.910 ± 0.042 | 0.900 ± 0.040 | 0.829 ± 0.048 | 0.874 ± 0.046 | 0.863 ± 0.041 |
| LPS_1.5 | 0.915 ± 0.041 | 0.906 ± 0.039 | 0.840 ± 0.047 | 0.881 ± 0.045 | 0.871 ± 0.040 |
| LPS_2.0 | 0.922 ± 0.040 | 0.913 ± 0.037 | 0.857 ± 0.045 | 0.892 ± 0.044 | 0.884 ± 0.038 |

Table S. 3 Imputation accuracy (mean and standard deviation across 22 autosomes) for eight genotyping arrays and six LPS coverages, evaluated across five populations for variant with allele frequency (0.05–0.5]

| Array/LPS | AFR | AMR | EAS | EUR | SAS |
|---|---|---|---|---|---|
| GSA | 0.826 ± 0.040 | 0.914 ± 0.031 | 0.882 ± 0.035 | 0.910 ± 0.031 | 0.893 ± 0.035 |
| JAPONICA | 0.861 ± 0.031 | 0.938 ± 0.022 | 0.935 ± 0.023 | 0.934 ± 0.021 | 0.927 ± 0.024 |
| UKB_WCSG | 0.856 ± 0.027 | 0.941 ± 0.022 | 0.909 ± 0.024 | 0.949 ± 0.021 | 0.927 ± 0.025 |
| CYTOSNP | 0.908 ± 0.031 | 0.944 ± 0.027 | 0.923 ± 0.031 | 0.943 ± 0.025 | 0.932 ± 0.031 |
| PMRA | 0.897 ± 0.024 | 0.935 ± 0.023 | 0.914 ± 0.025 | 0.933 ± 0.022 | 0.918 ± 0.025 |
| PMDA | 0.909 ± 0.017 | 0.945 ± 0.016 | 0.916 ± 0.018 | 0.945 ± 0.016 | 0.929 ± 0.018 |
| OMNI2.5 | 0.950 ± 0.025 | 0.962 ± 0.023 | 0.950 ± 0.025 | 0.963 ± 0.022 | 0.956 ± 0.026 |
| OMNI5 | 0.959 ± 0.022 | 0.970 ± 0.020 | 0.960 ± 0.022 | 0.972 ± 0.019 | 0.966 ± 0.022 |
| LPS_0.5 | 0.938 ± 0.035 | 0.947 ± 0.035 | 0.929 ± 0.037 | 0.945 ± 0.035 | 0.938 ± 0.037 |
| LPS_0.75 | 0.947 ± 0.036 | 0.954 ± 0.036 | 0.940 ± 0.037 | 0.953 ± 0.036 | 0.947 ± 0.037 |
| LPS_1.0 | 0.953 ± 0.034 | 0.959 ± 0.035 | 0.947 ± 0.037 | 0.958 ± 0.035 | 0.953 ± 0.036 |
| LPS_1.25 | 0.957 ± 0.033 | 0.963 ± 0.033 | 0.953 ± 0.035 | 0.961 ± 0.034 | 0.957 ± 0.035 |
| LPS_1.5 | 0.960 ± 0.032 | 0.965 ± 0.032 | 0.956 ± 0.034 | 0.964 ± 0.033 | 0.960 ± 0.034 |
| LPS_2.0 | 0.965 ± 0.030 | 0.968 ± 0.030 | 0.961 ± 0.032 | 0.968 ± 0.031 | 0.965 ± 0.032 |

Table S. 4 Imputation coverage (mean and standard deviation across 22 autosomes) for eight genotyping arrays and six LPS coverages, evaluated across five populations for variant with allele frequency (0–0.01]

| Array/LPS | AFR | AMR | EAS | EUR | SAS |
|---|---|---|---|---|---|
| GSA | 0.248 ± 0.041 | 0.489 ± 0.057 | 0.168 ± 0.033 | 0.307 ± 0.047 | 0.230 ± 0.035 |
| JAPONICA | 0.294 ± 0.041 | 0.527 ± 0.050 | 0.200 ± 0.036 | 0.343 ± 0.043 | 0.247 ± 0.036 |
| UKB_WCSG | 0.295 ± 0.031 | 0.538 ± 0.042 | 0.206 ± 0.028 | 0.369 ± 0.040 | 0.273 ± 0.030 |
| CYTOSNP | 0.364 ± 0.045 | 0.589 ± 0.048 | 0.222 ± 0.034 | 0.384 ± 0.039 | 0.259 ± 0.031 |
| PMRA | 0.318 ± 0.036 | 0.580 ± 0.042 | 0.214 ± 0.030 | 0.364 ± 0.036 | 0.251 ± 0.031 |
| PMDA | 0.331 ± 0.028 | 0.597 ± 0.030 | 0.208 ± 0.021 | 0.378 ± 0.027 | 0.256 ± 0.024 |
| OMNI2.5 | 0.487 ± 0.049 | 0.678 ± 0.047 | 0.284 ± 0.037 | 0.464 ± 0.042 | 0.339 ± 0.037 |
| OMNI5 | 0.538 ± 0.047 | 0.734 ± 0.046 | 0.319 ± 0.037 | 0.564 ± 0.046 | 0.425 ± 0.040 |
| LPS_0.5 | 0.543 ± 0.058 | 0.705 ± 0.053 | 0.314 ± 0.044 | 0.496 ± 0.052 | 0.388 ± 0.047 |
| LPS_0.75 | 0.581 ± 0.058 | 0.734 ± 0.052 | 0.355 ± 0.047 | 0.535 ± 0.053 | 0.436 ± 0.049 |
| LPS_1.0 | 0.613 ± 0.057 | 0.755 ± 0.051 | 0.392 ± 0.048 | 0.570 ± 0.052 | 0.476 ± 0.049 |
| LPS_1.25 | 0.634 ± 0.055 | 0.771 ± 0.050 | 0.422 ± 0.049 | 0.595 ± 0.052 | 0.508 ± 0.050 |
| LPS_1.5 | 0.651 ± 0.054 | 0.783 ± 0.048 | 0.447 ± 0.049 | 0.615 ± 0.051 | 0.534 ± 0.050 |
| LPS_2.0 | 0.679 ± 0.051 | 0.801 ± 0.047 | 0.491 ± 0.049 | 0.648 ± 0.050 | 0.575 ± 0.048 |

Table S. 5 Imputation coverage (mean and standard deviation across 22 autosomes) for eight genotyping arrays and six LPS coverages, evaluated across five populations for variant with allele frequency (0.01–0.05]

| Array/LPS | AFR | AMR | EAS | EUR | SAS |
|---|---|---|---|---|---|
| GSA | 0.400 ± 0.074 | 0.644 ± 0.070 | 0.480 ± 0.056 | 0.663 ± 0.062 | 0.478 ± 0.059 |
| JAPONICA | 0.522 ± 0.067 | 0.656 ± 0.061 | 0.563 ± 0.059 | 0.568 ± 0.062 | 0.508 ± 0.055 |
| UKB_WCSG | 0.466 ± 0.047 | 0.727 ± 0.048 | 0.448 ± 0.045 | 0.758 ± 0.056 | 0.570 ± 0.043 |
| CYTOSNP | 0.674 ± 0.069 | 0.714 ± 0.054 | 0.488 ± 0.050 | 0.608 ± 0.058 | 0.549 ± 0.052 |
| PMRA | 0.662 ± 0.047 | 0.722 ± 0.046 | 0.557 ± 0.049 | 0.632 ± 0.056 | 0.525 ± 0.043 |
| PMDA | 0.714 ± 0.037 | 0.771 ± 0.031 | 0.498 ± 0.043 | 0.682 ± 0.040 | 0.564 ± 0.039 |
| OMNI2.5 | 0.836 ± 0.054 | 0.811 ± 0.047 | 0.581 ± 0.050 | 0.733 ± 0.057 | 0.659 ± 0.048 |
| OMNI5 | 0.861 ± 0.049 | 0.870 ± 0.041 | 0.618 ± 0.048 | 0.866 ± 0.050 | 0.734 ± 0.047 |
| LPS_0.5 | 0.852 ± 0.066 | 0.811 ± 0.060 | 0.598 ± 0.064 | 0.732 ± 0.069 | 0.690 ± 0.062 |
| LPS_0.75 | 0.877 ± 0.063 | 0.839 ± 0.059 | 0.653 ± 0.066 | 0.777 ± 0.068 | 0.740 ± 0.061 |
| LPS_1.0 | 0.892 ± 0.061 | 0.857 ± 0.058 | 0.699 ± 0.066 | 0.812 ± 0.066 | 0.778 ± 0.059 |
| LPS_1.25 | 0.900 ± 0.059 | 0.871 ± 0.056 | 0.735 ± 0.064 | 0.834 ± 0.063 | 0.805 ± 0.057 |
| LPS_1.5 | 0.907 ± 0.056 | 0.880 ± 0.054 | 0.763 ± 0.063 | 0.850 ± 0.059 | 0.825 ± 0.055 |
| LPS_2.0 | 0.917 ± 0.049 | 0.893 ± 0.049 | 0.804 ± 0.058 | 0.872 ± 0.054 | 0.852 ± 0.050 |

Table S. 6 Imputation coverage (mean and standard deviation across 22 autosomes) for eight genotyping arrays and six LPS coverages, evaluated across five populations for variant with allele frequency (0.05–0.5]

| Array/LPS | AFR | AMR | EAS | EUR | SAS |
|---|---|---|---|---|---|
| GSA | 0.697 ± 0.078 | 0.893 ± 0.044 | 0.834 ± 0.051 | 0.882 ± 0.047 | 0.853 ± 0.052 |
| JAPONICA | 0.782 ± 0.058 | 0.930 ± 0.030 | 0.928 ± 0.030 | 0.917 ± 0.029 | 0.912 ± 0.034 |
| UKB_WCSG | 0.764 ± 0.045 | 0.943 ± 0.026 | 0.886 ± 0.030 | 0.952 ± 0.024 | 0.922 ± 0.030 |
| CYTOSNP | 0.881 ± 0.048 | 0.929 ± 0.033 | 0.897 ± 0.039 | 0.926 ± 0.031 | 0.911 ± 0.039 |
| PMRA | 0.872 ± 0.034 | 0.929 ± 0.029 | 0.898 ± 0.032 | 0.924 ± 0.028 | 0.902 ± 0.032 |
| PMDA | 0.907 ± 0.023 | 0.945 ± 0.018 | 0.892 ± 0.023 | 0.943 ± 0.018 | 0.918 ± 0.023 |
| OMNI2.5 | 0.946 ± 0.030 | 0.954 ± 0.027 | 0.937 ± 0.029 | 0.956 ± 0.025 | 0.948 ± 0.030 |
| OMNI5 | 0.956 ± 0.026 | 0.965 ± 0.024 | 0.949 ± 0.026 | 0.968 ± 0.022 | 0.960 ± 0.026 |
| LPS_0.5 | 0.935 ± 0.052 | 0.938 ± 0.053 | 0.908 ± 0.055 | 0.935 ± 0.052 | 0.924 ± 0.056 |
| LPS_0.75 | 0.943 ± 0.051 | 0.946 ± 0.053 | 0.924 ± 0.054 | 0.944 ± 0.052 | 0.936 ± 0.055 |
| LPS_1.0 | 0.948 ± 0.050 | 0.950 ± 0.052 | 0.934 ± 0.053 | 0.950 ± 0.051 | 0.943 ± 0.054 |
| LPS_1.25 | 0.951 ± 0.049 | 0.954 ± 0.050 | 0.940 ± 0.051 | 0.953 ± 0.049 | 0.947 ± 0.052 |
| LPS_1.5 | 0.953 ± 0.047 | 0.957 ± 0.047 | 0.947 ± 0.044 | 0.957 ± 0.045 | 0.951 ± 0.048 |
| LPS_2.0 | 0.960 ± 0.036 | 0.964 ± 0.037 | 0.955 ± 0.038 | 0.964 ± 0.036 | 0.960 ± 0.038 |

Table S. 7 Mean and the standard deviation of PGS correlation of eight genotyping arrays and six LPS coverages of the phenotype the phenotype body mass index (BMI)

| Array/LPS | AFR | AMR | EAS | EUR | SAS |
|---|---|---|---|---|---|
| GSA | 0.953 ± 0.007 | 0.983 ± 0.004 | 0.958 ± 0.011 | 0.979 ± 0.005 | 0.973 ± 0.008 |
| JAPONICA | 0.964 ± 0.005 | 0.987 ± 0.004 | 0.983 ± 0.004 | 0.984 ± 0.004 | 0.981 ± 0.005 |
| UKB_WCSG | 0.961 ± 0.006 | 0.990 ± 0.001 | 0.971 ± 0.007 | 0.992 ± 0.001 | 0.985 ± 0.003 |
| CYTOSNP | 0.984 ± 0.003 | 0.993 ± 0.002 | 0.983 ± 0.006 | 0.991 ± 0.004 | 0.988 ± 0.005 |
| PMRA | 0.967 ± 0.007 | 0.986 ± 0.002 | 0.967 ± 0.009 | 0.984 ± 0.005 | 0.976 ± 0.006 |
| PMDA | 0.969 ± 0.004 | 0.988 ± 0.003 | 0.968 ± 0.006 | 0.987 ± 0.004 | 0.978 ± 0.004 |
| OMNI2.5 | 0.995 ± 0.001 | 0.997 ± 0.001 | 0.994 ± 0.002 | 0.997 ± 0.001 | 0.996 ± 0.001 |
| OMNI5 | 0.997 ± 0.000 | 0.998 ± 0.000 | 0.996 ± 0.001 | 0.999 ± 0.000 | 0.998 ± 0.000 |
| LPS_0.5 | 0.983 ± 0.004 | 0.989 ± 0.003 | 0.973 ± 0.008 | 0.986 ± 0.005 | 0.982 ± 0.006 |
| LPS_0.75 | 0.987 ± 0.003 | 0.991 ± 0.003 | 0.977 ± 0.009 | 0.990 ± 0.003 | 0.986 ± 0.005 |
| LPS_1.0 | 0.990 ± 0.002 | 0.994 ± 0.001 | 0.983 ± 0.005 | 0.992 ± 0.002 | 0.990 ± 0.003 |
| LPS_1.25 | 0.991 ± 0.002 | 0.995 ± 0.002 | 0.986 ± 0.004 | 0.993 ± 0.002 | 0.991 ± 0.003 |
| LPS_1.5 | 0.992 ± 0.001 | 0.995 ± 0.001 | 0.989 ± 0.004 | 0.995 ± 0.002 | 0.992 ± 0.003 |
| LPS_2.0 | 0.994 ± 0.001 | 0.996 ± 0.001 | 0.991 ± 0.003 | 0.996 ± 0.001 | 0.994 ± 0.002 |

Table S. 8 Mean and the standard deviation of PGS correlation of eight genotyping arrays and six LPS coverages of the phenotype height

| Array/LPS | AFR | AMR | EAS | EUR | SAS |
|---|---|---|---|---|---|
| GSA | 0.947 ± 0.002 | 0.983 ± 0.001 | 0.963 ± 0.001 | 0.986 ± 0.001 | 0.972 ± 0.002 |
| JAPONICA | 0.961 ± 0.002 | 0.986 ± 0.001 | 0.984 ± 0.001 | 0.988 ± 0.002 | 0.982 ± 0.001 |
| UKB_WCSG | 0.956 ± 0.001 | 0.992 ± 0.000 | 0.976 ± 0.002 | 0.995 ± 0.000 | 0.987 ± 0.001 |
| CYTOSNP | 0.983 ± 0.002 | 0.993 ± 0.001 | 0.988 ± 0.002 | 0.994 ± 0.000 | 0.990 ± 0.002 |
| PMRA | 0.964 ± 0.002 | 0.986 ± 0.002 | 0.975 ± 0.001 | 0.989 ± 0.001 | 0.980 ± 0.002 |
| PMDA | 0.970 ± 0.001 | 0.987 ± 0.001 | 0.971 ± 0.002 | 0.991 ± 0.001 | 0.982 ± 0.001 |
| OMNI2.5 | 0.995 ± 0.000 | 0.997 ± 0.000 | 0.995 ± 0.000 | 0.998 ± 0.000 | 0.996 ± 0.001 |
| OMNI5 | 0.996 ± 0.000 | 0.999 ± 0.000 | 0.997 ± 0.000 | 0.999 ± 0.000 | 0.998 ± 0.000 |
| LPS_0.5 | 0.981 ± 0.001 | 0.987 ± 0.002 | 0.974 ± 0.003 | 0.990 ± 0.001 | 0.981 ± 0.002 |
| LPS_0.75 | 0.984 ± 0.001 | 0.990 ± 0.000 | 0.980 ± 0.001 | 0.993 ± 0.000 | 0.986 ± 0.001 |
| LPS_1.0 | 0.987 ± 0.001 | 0.992 ± 0.000 | 0.984 ± 0.002 | 0.994 ± 0.001 | 0.989 ± 0.001 |
| LPS_1.25 | 0.989 ± 0.001 | 0.993 ± 0.000 | 0.987 ± 0.001 | 0.995 ± 0.000 | 0.990 ± 0.001 |
| LPS_1.5 | 0.990 ± 0.001 | 0.994 ± 0.000 | 0.989 ± 0.001 | 0.996 ± 0.001 | 0.991 ± 0.001 |
| LPS_2.0 | 0.992 ± 0.001 | 0.995 ± 0.000 | 0.990 ± 0.001 | 0.996 ± 0.000 | 0.993 ± 0.000 |

Table S. 9 Mean and the standard deviation of PGS correlation of eight genotyping arrays and six LPS coverages of the phenotype diabetes

| Array/LPS | AFR | AMR | EAS | EUR | SAS |
|---|---|---|---|---|---|
| GSA | 0.960 ± 0.003 | 0.986 ± 0.003 | 0.960 ± 0.016 | 0.983 ± 0.005 | 0.976 ± 0.008 |
| JAPONICA | 0.967 ± 0.004 | 0.990 ± 0.002 | 0.984 ± 0.004 | 0.988 ± 0.003 | 0.982 ± 0.003 |
| UKB_WCSG | 0.962 ± 0.003 | 0.991 ± 0.001 | 0.973 ± 0.012 | 0.992 ± 0.002 | 0.984 ± 0.004 |
| CYTOSNP | 0.985 ± 0.001 | 0.995 ± 0.001 | 0.984 ± 0.003 | 0.993 ± 0.001 | 0.990 ± 0.003 |
| PMRA | 0.971 ± 0.002 | 0.989 ± 0.002 | 0.970 ± 0.011 | 0.987 ± 0.004 | 0.977 ± 0.005 |
| PMDA | 0.973 ± 0.003 | 0.990 ± 0.002 | 0.968 ± 0.009 | 0.989 ± 0.002 | 0.980 ± 0.004 |
| OMNI2.5 | 0.995 ± 0.000 | 0.998 ± 0.000 | 0.993 ± 0.001 | 0.998 ± 0.001 | 0.996 ± 0.001 |
| OMNI5 | 0.996 ± 0.000 | 0.999 ± 0.000 | 0.995 ± 0.001 | 0.999 ± 0.000 | 0.997 ± 0.001 |
| LPS_0.5 | 0.984 ± 0.002 | 0.989 ± 0.001 | 0.972 ± 0.007 | 0.987 ± 0.002 | 0.982 ± 0.001 |
| LPS_0.75 | 0.988 ± 0.001 | 0.992 ± 0.001 | 0.981 ± 0.004 | 0.990 ± 0.001 | 0.986 ± 0.002 |
| LPS_1.0 | 0.991 ± 0.001 | 0.994 ± 0.001 | 0.985 ± 0.004 | 0.992 ± 0.001 | 0.989 ± 0.001 |
| LPS_1.25 | 0.992 ± 0.001 | 0.995 ± 0.001 | 0.987 ± 0.002 | 0.993 ± 0.001 | 0.992 ± 0.001 |
| LPS_1.5 | 0.993 ± 0.001 | 0.996 ± 0.001 | 0.989 ± 0.003 | 0.994 ± 0.001 | 0.992 ± 0.001 |
| LPS_2.0 | 0.994 ± 0.001 | 0.996 ± 0.001 | 0.992 ± 0.003 | 0.995 ± 0.001 | 0.993 ± 0.001 |

Table S. 10 Mean and the standard deviation of PGS correlation of eight genotyping arrays and six LPS coverages of the phenotype metabolic

| Array/LPS | AFR | AMR | EAS | EUR | SAS |
|---|---|---|---|---|---|
| GSA | 0.955 ± 0.002 | 0.985 ± 0.002 | 0.959 ± 0.010 | 0.982 ± 0.001 | 0.972 ± 0.005 |
| JAPONICA | 0.969 ± 0.001 | 0.990 ± 0.001 | 0.984 ± 0.003 | 0.985 ± 0.002 | 0.979 ± 0.003 |
| UKB_WCSG | 0.961 ± 0.002 | 0.992 ± 0.001 | 0.973 ± 0.007 | 0.992 ± 0.000 | 0.986 ± 0.001 |
| CYTOSNP | 0.987 ± 0.001 | 0.995 ± 0.000 | 0.988 ± 0.005 | 0.993 ± 0.002 | 0.990 ± 0.003 |
| PMRA | 0.971 ± 0.002 | 0.988 ± 0.001 | 0.969 ± 0.008 | 0.985 ± 0.002 | 0.978 ± 0.005 |
| PMDA | 0.974 ± 0.003 | 0.991 ± 0.001 | 0.970 ± 0.008 | 0.988 ± 0.002 | 0.980 ± 0.003 |
| OMNI2.5 | 0.995 ± 0.000 | 0.997 ± 0.000 | 0.994 ± 0.001 | 0.997 ± 0.000 | 0.996 ± 0.001 |
| OMNI5 | 0.997 ± 0.000 | 0.999 ± 0.000 | 0.997 ± 0.001 | 0.999 ± 0.000 | 0.998 ± 0.000 |
| LPS_0.5 | 0.986 ± 0.001 | 0.991 ± 0.000 | 0.975 ± 0.008 | 0.988 ± 0.003 | 0.982 ± 0.006 |
| LPS_0.75 | 0.988 ± 0.001 | 0.994 ± 0.000 | 0.982 ± 0.007 | 0.991 ± 0.002 | 0.987 ± 0.004 |
| LPS_1.0 | 0.991 ± 0.001 | 0.995 ± 0.000 | 0.986 ± 0.005 | 0.993 ± 0.002 | 0.990 ± 0.003 |
| LPS_1.25 | 0.993 ± 0.001 | 0.996 ± 0.000 | 0.989 ± 0.004 | 0.994 ± 0.001 | 0.991 ± 0.003 |
| LPS_1.5 | 0.994 ± 0.000 | 0.996 ± 0.001 | 0.990 ± 0.004 | 0.995 ± 0.001 | 0.993 ± 0.003 |
| LPS_2.0 | 0.995 ± 0.000 | 0.997 ± 0.000 | 0.993 ± 0.003 | 0.996 ± 0.001 | 0.994 ± 0.002 |

Table S. 11 Mean absolute difference of percentile ranking between PGSs estimated from imputed genotyping data of eight genotyping arrays and six LPS coverages and PGS estimated from WGS in 5 different populations with PRsice p-value setting of 5e-08

| Trait | Array/LPS | AFR | AMR | EAS | EUR | SAS |
|---|---|---|---|---|---|---|
| BMI | GSA | 6.238 ± 5.600 | 3.508 ± 3.273 | 5.556 ± 5.350 | 3.860 ± 3.346 | 4.124 ± 3.773 |
| BMI | JAPONICA | 5.442 ± 4.932 | 3.030 ± 2.685 | 3.723 ± 3.493 | 3.126 ± 2.801 | 3.323 ± 3.006 |
| BMI | UKB_WCSG | 5.943 ± 5.295 | 2.918 ± 2.786 | 4.965 ± 4.503 | 2.639 ± 2.332 | 3.434 ± 3.077 |
| BMI | CYTOSNP | 3.490 ± 3.478 | 2.041 ± 1.945 | 3.430 ± 3.191 | 2.057 ± 1.951 | 2.452 ± 2.212 |
| BMI | PMRA | 5.108 ± 4.590 | 3.211 ± 2.978 | 5.002 ± 4.960 | 3.103 ± 3.016 | 4.006 ± 3.640 |
| BMI | PMDA | 5.334 ± 4.683 | 3.077 ± 2.769 | 5.050 ± 5.027 | 2.890 ± 2.575 | 3.972 ± 3.570 |
| BMI | OMNI2.5 | 2.133 ± 2.136 | 1.525 ± 1.515 | 2.105 ± 2.112 | 1.424 ± 1.360 | 1.590 ± 1.553 |
| BMI | OMNI5 | 1.709 ± 1.745 | 1.227 ± 1.302 | 1.843 ± 1.800 | 1.046 ± 1.148 | 1.264 ± 1.294 |
| BMI | LPS_0.5 | 3.799 ± 3.634 | 2.810 ± 2.552 | 4.573 ± 4.329 | 2.967 ± 2.714 | 3.220 ± 2.850 |
| BMI | LPS_0.75 | 3.369 ± 3.306 | 2.314 ± 2.150 | 3.606 ± 3.406 | 2.542 ± 2.408 | 2.883 ± 2.576 |
| BMI | LPS_1.0 | 2.802 ± 2.581 | 2.025 ± 1.938 | 3.474 ± 3.107 | 2.400 ± 2.311 | 2.647 ± 2.398 |
| BMI | LPS_1.25 | 2.736 ± 2.806 | 1.946 ± 1.657 | 3.059 ± 2.910 | 2.229 ± 2.145 | 2.429 ± 2.302 |
| BMI | LPS_1.5 | 2.554 ± 2.462 | 1.808 ± 1.664 | 2.800 ± 2.615 | 1.786 ± 1.706 | 2.143 ± 1.888 |
| BMI | LPS_2.0 | 2.205 ± 2.131 | 1.614 ± 1.472 | 2.482 ± 2.351 | 1.556 ± 1.462 | 2.082 ± 1.872 |
| DIABETES | GSA | 5.785 ± 5.322 | 3.219 ± 3.201 | 4.330 ± 3.956 | 2.932 ± 2.903 | 3.551 ± 3.538 |
| DIABETES | JAPONICA | 5.557 ± 5.429 | 2.634 ± 2.818 | 3.338 ± 3.078 | 2.649 ± 2.642 | 3.637 ± 3.432 |
| DIABETES | UKB_WCSG | 6.265 ± 6.166 | 2.733 ± 2.958 | 3.564 ± 3.280 | 2.182 ± 2.329 | 2.861 ± 3.009 |
| DIABETES | CYTOSNP | 3.736 ± 3.600 | 2.334 ± 2.384 | 3.475 ± 3.394 | 2.101 ± 2.671 | 2.352 ± 2.825 |
| DIABETES | PMRA | 5.535 ± 5.223 | 2.775 ± 3.142 | 4.174 ± 3.853 | 2.630 ± 2.782 | 3.890 ± 3.674 |
| DIABETES | PMDA | 5.221 ± 4.853 | 2.717 ± 2.808 | 4.549 ± 4.205 | 2.569 ± 2.542 | 3.556 ± 3.639 |
| DIABETES | OMNI2.5 | 2.113 ± 2.371 | 1.571 ± 1.927 | 2.479 ± 2.799 | 0.913 ± 1.551 | 1.538 ± 2.129 |
| DIABETES | OMNI5 | 1.761 ± 1.998 | 0.985 ± 1.488 | 2.072 ± 2.458 | 0.669 ± 1.307 | 1.247 ± 1.841 |

| | | | | | | |
|---|---|---|---|---|---|---|
| DIABETES | LPS_0.5 | 4.217 ± 4.228 | 3.831 ± 3.873 | 4.647 ± 4.637 | 3.604 ± 3.355 | 4.511 ± 4.373 |
| DIABETES | LPS_0.75 | 3.450 ± 3.163 | 2.947 ± 2.901 | 4.204 ± 3.945 | 3.084 ± 2.853 | 3.865 ± 3.672 |
| DIABETES | LPS_1.0 | 3.194 ± 3.178 | 2.713 ± 2.727 | 3.550 ± 3.336 | 3.066 ± 2.815 | 3.345 ± 3.068 |
| DIABETES | LPS_1.25 | 2.890 ± 2.917 | 2.452 ± 2.461 | 3.376 ± 3.154 | 2.697 ± 2.543 | 3.155 ± 3.100 |
| DIABETES | LPS_1.5 | 2.662 ± 2.652 | 2.457 ± 2.309 | 3.141 ± 2.794 | 2.686 ± 2.535 | 2.841 ± 2.665 |
| DIABETES | LPS_2.0 | 2.372 ± 2.268 | 2.375 ± 2.268 | 2.116 ± 2.184 | 2.496 ± 2.297 | 2.903 ± 2.608 |
| HEIGHT | GSA | 7.154 ± 6.563 | 3.888 ± 3.659 | 5.941 ± 5.495 | 3.987 ± 3.747 | 5.344 ± 4.797 |
| HEIGHT | JAPONICA | 6.566 ± 5.962 | 3.834 ± 3.639 | 3.758 ± 3.534 | 3.958 ± 3.474 | 4.274 ± 3.844 |
| HEIGHT | UKB_WCSG | 6.525 ± 5.817 | 3.043 ± 2.394 | 4.546 ± 4.281 | 2.391 ± 2.242 | 3.370 ± 3.157 |
| HEIGHT | CYTOSNP | 4.001 ± 3.694 | 2.549 ± 2.293 | 3.232 ± 2.839 | 2.727 ± 2.534 | 2.998 ± 2.772 |
| HEIGHT | PMRA | 5.795 ± 5.373 | 3.413 ± 3.083 | 5.008 ± 4.724 | 3.698 ± 3.385 | 4.314 ± 3.914 |
| HEIGHT | PMDA | 5.632 ± 5.255 | 3.614 ± 3.414 | 5.323 ± 4.955 | 3.399 ± 3.103 | 4.343 ± 4.003 |
| HEIGHT | OMNI2.5 | 2.306 ± 2.322 | 1.712 ± 1.661 | 2.185 ± 1.922 | 1.517 ± 1.472 | 1.933 ± 1.861 |
| HEIGHT | OMNI5 | 1.899 ± 1.758 | 1.264 ± 1.238 | 1.718 ± 1.596 | 1.063 ± 1.056 | 1.454 ± 1.373 |
| HEIGHT | LPS_0.5 | 4.416 ± 4.060 | 3.653 ± 3.164 | 4.774 ± 4.179 | 3.503 ± 3.254 | 3.966 ± 3.707 |
| HEIGHT | LPS_0.75 | 4.007 ± 3.718 | 3.021 ± 2.867 | 4.367 ± 3.979 | 2.921 ± 2.741 | 3.708 ± 3.376 |
| HEIGHT | LPS_1.0 | 3.612 ± 3.210 | 2.878 ± 2.554 | 3.872 ± 3.445 | 2.753 ± 2.531 | 3.280 ± 2.880 |
| HEIGHT | LPS_1.25 | 3.263 ± 2.934 | 2.654 ± 2.316 | 3.480 ± 3.095 | 2.477 ± 2.234 | 3.012 ± 2.854 |
| HEIGHT | LPS_1.5 | 3.015 ± 2.835 | 2.610 ± 2.371 | 3.205 ± 2.951 | 2.397 ± 2.194 | 2.991 ± 2.735 |
| HEIGHT | LPS_2.0 | 2.683 ± 2.512 | 2.389 ± 2.169 | 2.932 ± 2.692 | 2.236 ± 2.009 | 2.596 ± 2.585 |
| METABOLIC | GSA | 6.711 ± 6.067 | 4.181 ± 4.005 | 4.999 ± 4.543 | 4.013 ± 3.878 | 4.715 ± 4.405 |
| METABOLIC | JAPONICA | 5.834 ± 5.441 | 3.535 ± 3.556 | 3.568 ± 3.371 | 3.718 ± 3.434 | 4.319 ± 3.866 |
| METABOLIC | UKB_WCSG | 6.197 ± 5.556 | 3.279 ± 3.053 | 4.516 ± 4.129 | 2.675 ± 2.399 | 3.648 ± 3.672 |
| METABOLIC | CYTOSNP | 3.455 ± 3.375 | 2.437 ± 2.254 | 2.757 ± 2.636 | 2.260 ± 2.109 | 2.606 ± 2.510 |
| METABOLIC | PMRA | 5.260 ± | 3.862 ± | 4.436 ± | 3.532 ± | 4.195 ± |

| | | | | | | |
|---|---|---|---|---|---|---|
| | | 5.028 | 3.528 | 4.065 | 3.571 | 3.871 |
| METABOLIC | PMDA | 5.140 ± 4.902 | 3.237 ± 2.980 | 4.430 ± 4.003 | 3.209 ± 3.005 | 4.117 ± 3.890 |
| METABOLIC | OMNI2.5 | 2.438 ± 2.194 | 2.018 ± 2.073 | 2.294 ± 2.041 | 1.718 ± 1.611 | 1.834 ± 1.781 |
| METABOLIC | OMNI5 | 1.783 ± 1.702 | 1.289 ± 1.365 | 1.547 ± 1.511 | 1.011 ± 1.059 | 1.251 ± 1.227 |
| METABOLIC | LPS_0.5 | 3.766 ± 3.501 | 3.136 ± 2.960 | 3.812 ± 3.360 | 3.021 ± 2.772 | 3.318 ± 2.965 |
| METABOLIC | LPS_0.75 | 3.228 ± 3.027 | 2.520 ± 2.570 | 3.385 ± 3.044 | 2.624 ± 2.368 | 2.910 ± 2.795 |
| METABOLIC | LPS_1.0 | 2.779 ± 2.613 | 2.566 ± 2.413 | 2.772 ± 2.458 | 2.236 ± 2.047 | 2.463 ± 2.201 |
| METABOLIC | LPS_1.25 | 2.478 ± 2.328 | 2.080 ± 1.990 | 2.630 ± 2.420 | 2.047 ± 1.892 | 2.477 ± 2.428 |
| METABOLIC | LPS_1.5 | 2.464 ± 2.262 | 2.016 ± 2.027 | 2.425 ± 2.125 | 1.858 ± 1.735 | 2.072 ± 1.983 |
| METABOLIC | LPS_2.0 | 2.154 ± 1.949 | 1.830 ± 1.835 | 2.124 ± 2.012 | 1.649 ± 1.583 | 2.024 ± 1.939 |

Table S. 12 Mean absolute difference of percentile ranking between PGSs estimated from imputed genotyping data of eight genotyping arrays and six LPS coverages and PGS estimated from WGS in 5 different populations with PRsice p-value setting of 1e-07

| Trait | Array/LPS | AFR | AMR | EAS | EUR | SAS |
|---|---|---|---|---|---|---|
| BMI | GSA | 6.199 ± 5.721 | 3.538 ± 3.356 | 5.625 ± 5.278 | 3.861 ± 3.337 | 4.270 ± 3.843 |
| BMI | JAPONICA | 5.451 ± 5.033 | 3.207 ± 2.774 | 3.699 ± 3.448 | 3.203 ± 2.956 | 3.529 ± 3.143 |
| BMI | UKB_WCSG | 5.974 ± 5.374 | 3.098 ± 2.970 | 4.954 ± 4.552 | 2.500 ± 2.175 | 3.571 ± 3.134 |
| BMI | CYTOSNP | 3.460 ± 3.460 | 2.162 ± 2.038 | 3.438 ± 3.102 | 2.112 ± 2.053 | 2.653 ± 2.424 |
| BMI | PMRA | 4.989 ± 4.618 | 3.455 ± 3.122 | 5.079 ± 4.992 | 3.084 ± 2.980 | 4.100 ± 3.807 |
| BMI | PMDA | 5.359 ± 4.679 | 3.182 ± 2.895 | 5.200 ± 5.017 | 2.974 ± 2.578 | 4.092 ± 3.737 |
| BMI | OMNI2.5 | 2.104 ± 2.102 | 1.667 ± 1.657 | 2.102 ± 2.133 | 1.468 ± 1.369 | 1.677 ± 1.623 |
| BMI | OMNI5 | 1.672 ± 1.748 | 1.382 ± 1.567 | 1.838 ± 1.704 | 1.073 ± 1.153 | 1.292 ± 1.310 |
| BMI | LPS_0.5 | 3.791 ± 3.624 | 2.869 ± 2.613 | 4.563 ± 4.315 | 2.921 ± 2.618 | 3.472 ± 3.043 |
| BMI | LPS_0.75 | 3.391 ± 3.238 | 2.546 ± 2.307 | 3.680 ± 3.400 | 2.559 ± 2.329 | 3.013 ± 2.692 |
| BMI | LPS_1.0 | 2.863 ± 2.607 | 2.139 ± 2.174 | 3.524 ± 3.259 | 2.406 ± 2.287 | 2.803 ± 2.567 |
| BMI | LPS_1.25 | 2.754 ± 2.815 | 2.060 ± 1.823 | 3.062 ± 2.994 | 2.168 ± 2.055 | 2.546 ± 2.350 |
| BMI | LPS_1.5 | 2.628 ± 2.452 | 1.820 ± 1.756 | 2.750 ± 2.601 | 1.790 ± 1.693 | 2.236 ± 2.083 |
| BMI | LPS_2.0 | 2.240 ± 2.208 | 1.716 ± 1.583 | 2.555 ± 2.415 | 1.555 ± 1.510 | 2.164 ± 1.968 |
| DIABETES | GSA | 6.020 ± 5.633 | 3.109 ± 2.974 | 4.139 ± 3.852 | 3.005 ± 2.961 | 3.536 ± 3.559 |
| DIABETES | JAPONICA | 5.646 ± 5.618 | 2.564 ± 2.664 | 3.244 ± 2.935 | 2.777 ± 2.769 | 3.627 ± 3.396 |
| DIABETES | UKB_WCSG | 6.378 ± 6.097 | 2.707 ± 2.669 | 3.668 ± 3.208 | 2.246 ± 2.394 | 3.023 ± 2.981 |
| DIABETES | CYTOSNP | 3.741 ± 3.589 | 2.296 ± 2.204 | 3.503 ± 3.394 | 2.120 ± 2.688 | 2.415 ± 2.729 |
| DIABETES | PMRA | 5.747 ± 5.254 | 2.920 ± 3.191 | 4.130 ± 3.813 | 2.781 ± 2.834 | 4.031 ± 3.811 |
| DIABETES | PMDA | 5.386 ± 5.059 | 2.895 ± 2.967 | 4.335 ± 3.986 | 2.754 ± 2.778 | 3.823 ± 3.600 |
| DIABETES | OMNI2.5 | 2.175 ± 2.377 | 1.614 ± 1.830 | 2.444 ± 2.723 | 0.975 ± 1.602 | 1.639 ± 2.089 |
| DIABETES | OMNI5 | 1.789 ± 1.992 | 1.021 ± 1.374 | 1.982 ± 2.306 | 0.721 ± 1.288 | 1.259 ± 1.779 |

| | | | | | | |
|---|---|---|---|---|---|---|
| DIABETES | LPS_0.5 | 4.428 ± 4.425 | 3.734 ± 3.522 | 4.554 ± 4.461 | 3.665 ± 3.318 | 4.596 ± 4.356 |
| DIABETES | LPS_0.75 | 3.613 ± 3.328 | 2.885 ± 2.804 | 4.019 ± 3.749 | 3.089 ± 2.779 | 3.881 ± 3.705 |
| DIABETES | LPS_1.0 | 3.427 ± 3.322 | 2.699 ± 2.738 | 3.383 ± 3.153 | 3.042 ± 2.814 | 3.477 ± 3.194 |
| DIABETES | LPS_1.25 | 3.004 ± 2.971 | 2.405 ± 2.385 | 3.247 ± 3.006 | 2.724 ± 2.592 | 3.259 ± 3.107 |
| DIABETES | LPS_1.5 | 2.831 ± 2.794 | 2.340 ± 2.315 | 2.992 ± 2.667 | 2.662 ± 2.544 | 2.978 ± 2.847 |
| DIABETES | LPS_2.0 | 2.509 ± 2.431 | 2.296 ± 2.045 | 2.131 ± 2.119 | 2.502 ± 2.309 | 3.038 ± 2.716 |
| HEIGHT | GSA | 7.328 ± 6.591 | 3.987 ± 3.705 | 5.894 ± 5.487 | 3.975 ± 3.714 | 5.271 ± 4.657 |
| HEIGHT | JAPONICA | 6.550 ± 5.988 | 3.832 ± 3.688 | 3.756 ± 3.597 | 3.873 ± 3.512 | 4.253 ± 3.867 |
| HEIGHT | UKB_WCSG | 6.652 ± 5.886 | 2.983 ± 2.410 | 4.608 ± 4.227 | 2.342 ± 2.225 | 3.457 ± 3.215 |
| HEIGHT | CYTOSNP | 4.003 ± 3.655 | 2.501 ± 2.314 | 3.258 ± 2.977 | 2.752 ± 2.541 | 3.017 ± 2.722 |
| HEIGHT | PMRA | 5.869 ± 5.445 | 3.337 ± 3.068 | 4.915 ± 4.667 | 3.650 ± 3.332 | 4.299 ± 3.942 |
| HEIGHT | PMDA | 5.664 ± 5.179 | 3.700 ± 3.335 | 5.380 ± 4.923 | 3.354 ± 3.058 | 4.279 ± 3.980 |
| HEIGHT | OMNI2.5 | 2.303 ± 2.297 | 1.670 ± 1.605 | 2.163 ± 1.949 | 1.540 ± 1.481 | 2.012 ± 1.935 |
| HEIGHT | OMNI5 | 1.886 ± 1.694 | 1.197 ± 1.191 | 1.785 ± 1.595 | 1.049 ± 1.004 | 1.468 ± 1.342 |
| HEIGHT | LPS_0.5 | 4.434 ± 4.146 | 3.582 ± 3.236 | 4.698 ± 4.273 | 3.435 ± 3.171 | 4.002 ± 3.779 |
| HEIGHT | LPS_0.75 | 4.009 ± 3.735 | 3.073 ± 2.806 | 4.399 ± 4.042 | 2.919 ± 2.670 | 3.703 ± 3.354 |
| HEIGHT | LPS_1.0 | 3.595 ± 3.237 | 2.830 ± 2.458 | 3.930 ± 3.577 | 2.724 ± 2.531 | 3.283 ± 2.944 |
| HEIGHT | LPS_1.25 | 3.281 ± 2.966 | 2.626 ± 2.344 | 3.396 ± 3.143 | 2.418 ± 2.145 | 3.005 ± 2.815 |
| HEIGHT | LPS_1.5 | 3.066 ± 2.851 | 2.594 ± 2.378 | 3.175 ± 2.966 | 2.356 ± 2.145 | 2.996 ± 2.756 |
| HEIGHT | LPS_2.0 | 2.733 ± 2.509 | 2.344 ± 2.203 | 2.955 ± 2.701 | 2.253 ± 1.961 | 2.556 ± 2.585 |
| METABOLIC | GSA | 6.668 ± 6.045 | 4.375 ± 4.209 | 5.263 ± 4.846 | 3.967 ± 3.691 | 4.972 ± 4.611 |
| METABOLIC | JAPONICA | 5.704 ± 5.238 | 3.684 ± 3.655 | 3.645 ± 3.394 | 3.752 ± 3.305 | 4.392 ± 4.037 |
| METABOLIC | UKB_WCSG | 6.295 ± 5.573 | 3.392 ± 3.221 | 4.686 ± 4.347 | 2.707 ± 2.414 | 3.650 ± 3.705 |
| METABOLIC | CYTOSNP | 3.510 ± 3.363 | 2.432 ± 2.244 | 2.930 ± 2.684 | 2.225 ± 2.052 | 2.656 ± 2.596 |
| METABOLIC | PMRA | 5.280 ± | 3.812 ± | 4.412 ± | 3.498 ± | 4.340 ± |

| | | | | | | |
|---|---|---|---|---|---|---|
| | | 4.999 | 3.690 | 4.293 | 3.522 | 4.150 |
| METABOLIC | PMDA | 5.183 ± 4.837 | 3.216 ± 2.863 | 4.504 ± 4.161 | 3.207 ± 3.076 | 4.126 ± 3.856 |
| METABOLIC | OMNI2.5 | 2.450 ± 2.219 | 2.010 ± 1.933 | 2.408 ± 2.125 | 1.673 ± 1.502 | 1.883 ± 1.803 |
| METABOLIC | OMNI5 | 1.796 ± 1.708 | 1.319 ± 1.343 | 1.608 ± 1.557 | 1.037 ± 1.035 | 1.296 ± 1.282 |
| METABOLIC | LPS_0.5 | 3.847 ± 3.559 | 3.028 ± 2.869 | 3.841 ± 3.504 | 2.887 ± 2.764 | 3.418 ± 3.160 |
| METABOLIC | LPS_0.75 | 3.345 ± 3.032 | 2.654 ± 2.696 | 3.463 ± 3.030 | 2.629 ± 2.403 | 2.999 ± 2.857 |
| METABOLIC | LPS_1.0 | 2.796 ± 2.679 | 2.535 ± 2.472 | 2.831 ± 2.466 | 2.266 ± 2.007 | 2.549 ± 2.362 |
| METABOLIC | LPS_1.25 | 2.590 ± 2.460 | 2.154 ± 2.006 | 2.694 ± 2.414 | 2.013 ± 1.826 | 2.485 ± 2.418 |
| METABOLIC | LPS_1.5 | 2.508 ± 2.338 | 2.114 ± 1.948 | 2.505 ± 2.202 | 1.836 ± 1.660 | 2.108 ± 1.962 |
| METABOLIC | LPS_2.0 | 2.190 ± 2.058 | 1.940 ± 1.911 | 2.190 ± 2.067 | 1.602 ± 1.468 | 2.058 ± 1.829 |

Table S. 13 Mean absolute difference of percentile ranking between PGSs estimated from imputed genotyping data of eight genotyping arrays and six LPS coverages and PGS estimated from WGS in 5 different populations with PRsice p-value setting of 1e-06

| Trait | Array/LPS | AFR | AMR | EAS | EUR | SAS |
|---|---|---|---|---|---|---|
| BMI | GSA | 6.585 ± 6.475 | 3.590 ± 3.628 | 5.691 ± 5.234 | 4.071 ± 3.395 | 4.478 ± 3.978 |
| BMI | JAPONICA | 5.708 ± 5.332 | 3.205 ± 3.000 | 3.574 ± 3.160 | 3.227 ± 2.951 | 3.561 ± 3.031 |
| BMI | UKB_WCSG | 5.966 ± 5.457 | 3.099 ± 2.871 | 4.970 ± 4.511 | 2.519 ± 2.152 | 3.499 ± 3.050 |
| BMI | CYTOSNP | 3.612 ± 3.526 | 2.119 ± 2.038 | 3.456 ± 3.258 | 2.305 ± 2.149 | 2.598 ± 2.472 |
| BMI | PMRA | 5.226 ± 4.658 | 3.689 ± 3.393 | 5.041 ± 4.662 | 3.351 ± 3.105 | 4.054 ± 3.549 |
| BMI | PMDA | 5.428 ± 5.002 | 3.142 ± 2.920 | 5.254 ± 4.788 | 2.916 ± 2.589 | 4.038 ± 3.614 |
| BMI | OMNI2.5 | 2.234 ± 2.201 | 1.732 ± 1.673 | 2.056 ± 2.007 | 1.490 ± 1.426 | 1.712 ± 1.694 |
| BMI | OMNI5 | 1.790 ± 1.836 | 1.351 ± 1.523 | 1.781 ± 1.681 | 1.079 ± 1.107 | 1.393 ± 1.322 |
| BMI | LPS_0.5 | 3.743 ± 3.477 | 2.941 ± 2.614 | 4.454 ± 4.214 | 2.866 ± 2.523 | 3.370 ± 2.968 |
| BMI | LPS_0.75 | 3.407 ± 3.376 | 2.496 ± 2.309 | 3.697 ± 3.505 | 2.623 ± 2.445 | 3.060 ± 2.789 |
| BMI | LPS_1.0 | 2.855 ± 2.649 | 2.270 ± 2.252 | 3.412 ± 3.130 | 2.301 ± 2.168 | 2.633 ± 2.430 |
| BMI | LPS_1.25 | 2.708 ± 2.765 | 2.086 ± 1.825 | 3.057 ± 2.991 | 1.981 ± 1.902 | 2.481 ± 2.155 |
| BMI | LPS_1.5 | 2.661 ± 2.532 | 1.960 ± 1.824 | 2.668 ± 2.564 | 1.750 ± 1.680 | 2.246 ± 2.115 |
| BMI | LPS_2.0 | 2.224 ± 2.220 | 1.810 ± 1.743 | 2.525 ± 2.374 | 1.658 ± 1.596 | 2.141 ± 2.032 |
| DIABETES | GSA | 6.534 ± 6.318 | 3.570 ± 3.000 | 4.849 ± 4.549 | 3.569 ± 3.282 | 3.988 ± 3.656 |
| DIABETES | JAPONICA | 6.052 ± 5.813 | 2.830 ± 2.921 | 3.540 ± 3.221 | 2.962 ± 2.958 | 3.753 ± 3.383 |
| DIABETES | UKB_WCSG | 6.887 ± 6.303 | 2.782 ± 2.732 | 3.936 ± 3.570 | 2.400 ± 2.524 | 3.277 ± 3.097 |
| DIABETES | CYTOSNP | 4.022 ± 3.926 | 2.102 ± 2.104 | 3.607 ± 3.490 | 2.126 ± 2.459 | 2.407 ± 2.628 |
| DIABETES | PMRA | 5.588 ± 5.222 | 3.107 ± 2.991 | 4.304 ± 4.076 | 2.990 ± 2.975 | 4.094 ± 3.827 |
| DIABETES | PMDA | 5.662 ± 5.436 | 2.864 ± 2.644 | 4.910 ± 4.643 | 2.860 ± 2.957 | 3.917 ± 3.701 |
| DIABETES | OMNI2.5 | 2.236 ± 2.416 | 1.504 ± 1.664 | 2.466 ± 2.685 | 1.164 ± 1.584 | 1.648 ± 1.859 |
| DIABETES | OMNI5 | 1.908 ± 2.167 | 0.992 ± 1.256 | 2.075 ± 2.333 | 0.924 ± 1.379 | 1.320 ± 1.596 |

| | | | | | | |
|---|---|---|---|---|---|---|
| DIABETES | LPS_0.5 | 4.403 ± 4.289 | 3.298 ± 3.128 | 4.601 ± 4.427 | 3.510 ± 3.393 | 4.203 ± 3.906 |
| DIABETES | LPS_0.75 | 3.561 ± 3.373 | 2.605 ± 2.418 | 4.001 ± 3.771 | 2.957 ± 2.787 | 3.466 ± 3.150 |
| DIABETES | LPS_1.0 | 3.349 ± 3.342 | 2.443 ± 2.355 | 3.385 ± 3.212 | 2.948 ± 2.904 | 3.131 ± 2.729 |
| DIABETES | LPS_1.25 | 2.916 ± 2.794 | 2.150 ± 1.984 | 3.241 ± 3.164 | 2.564 ± 2.595 | 2.758 ± 2.825 |
| DIABETES | LPS_1.5 | 2.823 ± 2.671 | 2.130 ± 2.058 | 3.086 ± 2.757 | 2.517 ± 2.632 | 2.579 ± 2.290 |
| DIABETES | LPS_2.0 | 2.477 ± 2.354 | 1.966 ± 1.839 | 2.262 ± 2.228 | 2.337 ± 2.274 | 2.620 ± 2.410 |
| HEIGHT | GSA | 7.290 ± 6.511 | 4.188 ± 4.028 | 5.950 ± 5.306 | 3.993 ± 3.836 | 5.371 ± 4.748 |
| HEIGHT | JAPONICA | 6.572 ± 5.859 | 4.042 ± 3.982 | 3.842 ± 3.562 | 3.761 ± 3.481 | 4.373 ± 3.916 |
| HEIGHT | UKB_WCSG | 6.681 ± 5.888 | 3.120 ± 2.584 | 4.779 ± 4.291 | 2.281 ± 2.196 | 3.411 ± 3.080 |
| HEIGHT | CYTOSNP | 4.045 ± 3.695 | 2.632 ± 2.462 | 3.366 ± 2.914 | 2.724 ± 2.522 | 3.033 ± 2.850 |
| HEIGHT | PMRA | 5.938 ± 5.451 | 3.500 ± 3.185 | 4.952 ± 4.344 | 3.617 ± 3.116 | 4.441 ± 4.078 |
| HEIGHT | PMDA | 5.731 ± 5.233 | 3.943 ± 3.647 | 5.373 ± 4.887 | 3.331 ± 3.052 | 4.414 ± 4.034 |
| HEIGHT | OMNI2.5 | 2.333 ± 2.268 | 1.717 ± 1.664 | 2.232 ± 1.985 | 1.579 ± 1.521 | 2.000 ± 1.924 |
| HEIGHT | OMNI5 | 1.916 ± 1.701 | 1.267 ± 1.201 | 1.702 ± 1.595 | 1.039 ± 1.006 | 1.503 ± 1.449 |
| HEIGHT | LPS_0.5 | 4.385 ± 4.025 | 3.764 ± 3.431 | 4.706 ± 4.227 | 3.422 ± 2.965 | 4.270 ± 3.840 |
| HEIGHT | LPS_0.75 | 4.015 ± 3.608 | 3.154 ± 3.085 | 4.375 ± 3.836 | 2.899 ± 2.568 | 3.732 ± 3.416 |
| HEIGHT | LPS_1.0 | 3.621 ± 3.177 | 3.020 ± 2.729 | 3.929 ± 3.527 | 2.644 ± 2.498 | 3.322 ± 3.039 |
| HEIGHT | LPS_1.25 | 3.323 ± 2.932 | 2.747 ± 2.596 | 3.349 ± 3.130 | 2.558 ± 2.226 | 3.056 ± 2.736 |
| HEIGHT | LPS_1.5 | 3.079 ± 2.781 | 2.695 ± 2.436 | 3.200 ± 2.968 | 2.384 ± 2.185 | 2.912 ± 2.673 |
| HEIGHT | LPS_2.0 | 2.791 ± 2.564 | 2.444 ± 2.207 | 2.934 ± 2.617 | 2.185 ± 1.937 | 2.721 ± 2.516 |
| METABOLIC | GSA | 6.898 ± 6.078 | 4.466 ± 4.075 | 5.305 ± 4.567 | 4.098 ± 3.802 | 4.978 ± 4.527 |
| METABOLIC | JAPONICA | 5.805 ± 5.261 | 3.598 ± 3.398 | 3.521 ± 3.172 | 3.770 ± 3.218 | 4.576 ± 4.209 |
| METABOLIC | UKB_WCSG | 6.310 ± 5.651 | 3.261 ± 3.041 | 4.440 ± 3.920 | 2.655 ± 2.397 | 3.694 ± 3.539 |
| METABOLIC | CYTOSNP | 3.603 ± 3.477 | 2.372 ± 2.134 | 2.789 ± 2.629 | 2.258 ± 2.054 | 2.647 ± 2.621 |
| METABOLIC | PMRA | 5.381 ± | 3.910 ± | 4.539 ± | 3.555 ± | 4.327 ± |

| | | | | | | |
|---|---|---|---|---|---|---|
| | | 5.005 | 3.757 | 4.303 | 3.440 | 3.932 |
| METABOLIC | PMDA | 5.463 ± 5.257 | 3.168 ± 2.959 | 4.613 ± 3.998 | 3.255 ± 3.072 | 4.207 ± 3.929 |
| METABOLIC | OMNI2.5 | 2.422 ± 2.294 | 1.887 ± 1.839 | 2.180 ± 1.944 | 1.616 ± 1.434 | 1.882 ± 1.934 |
| METABOLIC | OMNI5 | 1.713 ± 1.681 | 1.309 ± 1.350 | 1.499 ± 1.360 | 1.052 ± 1.028 | 1.362 ± 1.373 |
| METABOLIC | LPS_0.5 | 3.754 ± 3.344 | 3.001 ± 2.683 | 3.942 ± 3.550 | 2.869 ± 2.742 | 3.469 ± 3.290 |
| METABOLIC | LPS_0.75 | 3.305 ± 3.035 | 2.681 ± 2.511 | 3.320 ± 3.136 | 2.735 ± 2.512 | 2.995 ± 2.862 |
| METABOLIC | LPS_1.0 | 2.895 ± 2.711 | 2.518 ± 2.479 | 2.900 ± 2.537 | 2.224 ± 2.024 | 2.752 ± 2.561 |
| METABOLIC | LPS_1.25 | 2.497 ± 2.196 | 1.993 ± 1.780 | 2.675 ± 2.366 | 2.027 ± 1.868 | 2.565 ± 2.498 |
| METABOLIC | LPS_1.5 | 2.417 ± 2.251 | 2.041 ± 2.026 | 2.454 ± 2.227 | 2.006 ± 1.815 | 2.034 ± 1.949 |
| METABOLIC | LPS_2.0 | 2.195 ± 1.970 | 2.031 ± 1.966 | 2.187 ± 2.064 | 1.753 ± 1.622 | 2.176 ± 2.022 |

Table S. 14 Mean absolute difference of percentile ranking between PGSs estimated from imputed genotyping data of eight genotyping arrays and six LPS coverages and PGS estimated from WGS in 5 different populations with PRsice p-value setting of 1e-05

| Trait | Array/LPS | AFR | AMR | EAS | EUR | SAS |
|---|---|---|---|---|---|---|
| BMI | GSA | 6.707 ± 6.253 | 3.520 ± 3.288 | 5.610 ± 4.866 | 4.157 ± 3.535 | 4.446 ± 4.070 |
| BMI | JAPONICA | 5.857 ± 5.176 | 3.122 ± 2.880 | 3.538 ± 3.161 | 3.533 ± 3.233 | 3.719 ± 3.265 |
| BMI | UKB_WCSG | 6.068 ± 5.862 | 3.149 ± 3.121 | 4.818 ± 4.301 | 2.564 ± 2.229 | 3.595 ± 3.246 |
| BMI | CYTOSNP | 3.735 ± 3.665 | 2.298 ± 2.259 | 3.482 ± 3.231 | 2.295 ± 2.154 | 2.681 ± 2.487 |
| BMI | PMRA | 5.403 ± 4.869 | 3.560 ± 3.279 | 5.130 ± 4.588 | 3.710 ± 3.384 | 4.512 ± 4.167 |
| BMI | PMDA | 5.629 ± 4.904 | 3.201 ± 3.083 | 5.234 ± 4.822 | 3.167 ± 2.759 | 3.984 ± 3.810 |
| BMI | OMNI2.5 | 2.189 ± 2.063 | 1.720 ± 1.526 | 2.089 ± 2.063 | 1.537 ± 1.479 | 1.728 ± 1.602 |
| BMI | OMNI5 | 1.791 ± 1.778 | 1.333 ± 1.368 | 1.761 ± 1.599 | 1.087 ± 1.005 | 1.449 ± 1.397 |
| BMI | LPS_0.5 | 3.760 ± 3.528 | 3.065 ± 2.717 | 4.656 ± 4.299 | 3.003 ± 2.815 | 3.542 ± 3.051 |
| BMI | LPS_0.75 | 3.329 ± 3.234 | 2.419 ± 2.330 | 3.847 ± 3.489 | 2.715 ± 2.381 | 3.220 ± 3.050 |
| BMI | LPS_1.0 | 2.996 ± 2.778 | 2.384 ± 2.174 | 3.521 ± 3.023 | 2.390 ± 2.181 | 2.885 ± 2.765 |
| BMI | LPS_1.25 | 2.823 ± 2.798 | 2.020 ± 1.922 | 3.212 ± 2.993 | 2.217 ± 2.031 | 2.739 ± 2.382 |
| BMI | LPS_1.5 | 2.645 ± 2.466 | 1.967 ± 1.859 | 2.838 ± 2.546 | 1.878 ± 1.742 | 2.313 ± 2.143 |
| BMI | LPS_2.0 | 2.328 ± 2.186 | 1.766 ± 1.676 | 2.551 ± 2.221 | 1.702 ± 1.530 | 2.152 ± 2.003 |
| DIABETES | GSA | 6.684 ± 5.810 | 3.254 ± 2.949 | 5.023 ± 4.792 | 3.673 ± 3.357 | 4.277 ± 3.895 |
| DIABETES | JAPONICA | 5.493 ± 5.297 | 2.814 ± 2.671 | 3.647 ± 3.457 | 2.958 ± 2.941 | 3.929 ± 3.393 |
| DIABETES | UKB_WCSG | 6.198 ± 5.619 | 2.789 ± 2.353 | 4.000 ± 3.659 | 2.421 ± 2.470 | 3.401 ± 3.240 |
| DIABETES | CYTOSNP | 3.970 ± 3.683 | 2.086 ± 2.023 | 3.467 ± 3.310 | 2.156 ± 2.285 | 2.501 ± 2.518 |
| DIABETES | PMRA | 5.392 ± 5.098 | 2.899 ± 2.648 | 4.309 ± 4.293 | 3.192 ± 3.060 | 4.310 ± 3.970 |
| DIABETES | PMDA | 5.333 ± 4.784 | 2.777 ± 2.509 | 4.871 ± 4.561 | 3.057 ± 2.934 | 4.172 ± 3.925 |
| DIABETES | OMNI2.5 | 2.224 ± 2.249 | 1.543 ± 1.582 | 2.376 ± 2.439 | 1.199 ± 1.381 | 1.752 ± 1.827 |
| DIABETES | OMNI5 | 1.928 ± 2.051 | 1.110 ± 1.242 | 2.012 ± 2.039 | 0.929 ± 1.220 | 1.384 ± 1.540 |

| | | | | | | |
|---|---|---|---|---|---|---|
| DIABETES | LPS_0.5 | 4.186 ± 3.933 | 3.044 ± 2.831 | 4.355 ± 4.168 | 3.446 ± 3.070 | 4.135 ± 3.753 |
| DIABETES | LPS_0.75 | 3.238 ± 3.092 | 2.529 ± 2.361 | 3.846 ± 3.484 | 2.819 ± 2.607 | 3.463 ± 3.077 |
| DIABETES | LPS_1.0 | 3.119 ± 3.083 | 2.130 ± 2.042 | 3.361 ± 3.204 | 2.778 ± 2.619 | 3.194 ± 2.777 |
| DIABETES | LPS_1.25 | 2.653 ± 2.571 | 2.097 ± 1.992 | 3.206 ± 3.285 | 2.389 ± 2.276 | 2.949 ± 2.688 |
| DIABETES | LPS_1.5 | 2.553 ± 2.392 | 1.964 ± 1.914 | 2.950 ± 2.832 | 2.373 ± 2.288 | 2.776 ± 2.400 |
| DIABETES | LPS_2.0 | 2.286 ± 2.219 | 1.797 ± 1.648 | 2.292 ± 2.212 | 2.187 ± 1.997 | 2.822 ± 2.431 |
| HEIGHT | GSA | 7.251 ± 6.672 | 4.209 ± 3.970 | 5.930 ± 5.366 | 4.039 ± 3.750 | 5.478 ± 4.765 |
| HEIGHT | JAPONICA | 6.585 ± 5.871 | 3.969 ± 3.818 | 3.933 ± 3.668 | 3.697 ± 3.441 | 4.274 ± 3.947 |
| HEIGHT | UKB_WCSG | 6.680 ± 6.148 | 3.078 ± 2.580 | 4.804 ± 4.336 | 2.365 ± 2.222 | 3.442 ± 3.181 |
| HEIGHT | CYTOSNP | 4.016 ± 3.828 | 2.730 ± 2.448 | 3.377 ± 2.993 | 2.624 ± 2.472 | 3.163 ± 2.919 |
| HEIGHT | PMRA | 5.977 ± 5.669 | 3.531 ± 3.388 | 4.870 ± 4.287 | 3.648 ± 3.241 | 4.436 ± 3.996 |
| HEIGHT | PMDA | 5.627 ± 5.093 | 3.790 ± 3.540 | 5.284 ± 4.780 | 3.239 ± 2.992 | 4.475 ± 4.124 |
| HEIGHT | OMNI2.5 | 2.371 ± 2.229 | 1.715 ± 1.696 | 2.176 ± 1.972 | 1.538 ± 1.475 | 2.055 ± 1.976 |
| HEIGHT | OMNI5 | 1.936 ± 1.738 | 1.268 ± 1.235 | 1.768 ± 1.692 | 1.043 ± 1.040 | 1.580 ± 1.427 |
| HEIGHT | LPS_0.5 | 4.398 ± 4.029 | 3.849 ± 3.509 | 4.732 ± 4.319 | 3.348 ± 3.132 | 4.389 ± 3.926 |
| HEIGHT | LPS_0.75 | 4.041 ± 3.818 | 3.266 ± 3.145 | 4.433 ± 3.823 | 2.971 ± 2.751 | 3.697 ± 3.260 |
| HEIGHT | LPS_1.0 | 3.506 ± 3.238 | 2.991 ± 2.703 | 4.083 ± 3.626 | 2.666 ± 2.364 | 3.344 ± 3.063 |
| HEIGHT | LPS_1.25 | 3.418 ± 2.995 | 2.731 ± 2.554 | 3.491 ± 3.227 | 2.547 ± 2.279 | 3.048 ± 2.771 |
| HEIGHT | LPS_1.5 | 3.106 ± 2.986 | 2.662 ± 2.400 | 3.315 ± 3.086 | 2.283 ± 2.087 | 2.886 ± 2.645 |
| HEIGHT | LPS_2.0 | 2.858 ± 2.675 | 2.526 ± 2.303 | 3.016 ± 2.767 | 2.161 ± 1.965 | 2.697 ± 2.600 |
| METABOLIC | GSA | 7.064 ± 6.211 | 4.309 ± 3.949 | 5.501 ± 4.742 | 3.907 ± 3.671 | 4.997 ± 4.461 |
| METABOLIC | JAPONICA | 5.846 ± 5.148 | 3.501 ± 3.419 | 3.474 ± 3.040 | 3.730 ± 3.343 | 4.182 ± 3.687 |
| METABOLIC | UKB_WCSG | 6.444 ± 5.697 | 3.174 ± 3.214 | 4.360 ± 3.903 | 2.703 ± 2.446 | 3.510 ± 3.292 |
| METABOLIC | CYTOSNP | 3.674 ± 3.387 | 2.422 ± 2.345 | 2.712 ± 2.417 | 2.210 ± 2.015 | 2.606 ± 2.482 |
| METABOLIC | PMRA | 5.440 ± | 3.940 ± | 4.552 ± | 3.574 ± | 4.223 ± |

| | | | | | | |
|---|---|---|---|---|---|---|
| | | 5.202 | 3.743 | 4.152 | 3.400 | 3.766 |
| METABOLIC | PMDA | 5.542 ± 4.919 | 3.199 ± 3.175 | 4.626 ± 4.014 | 3.290 ± 2.994 | 4.210 ± 4.009 |
| METABOLIC | OMNI2.5 | 2.484 ± 2.170 | 1.827 ± 1.690 | 2.316 ± 1.932 | 1.716 ± 1.598 | 1.913 ± 1.841 |
| METABOLIC | OMNI5 | 1.790 ± 1.689 | 1.214 ± 1.229 | 1.593 ± 1.439 | 1.098 ± 1.071 | 1.434 ± 1.425 |
| METABOLIC | LPS_0.5 | 3.830 ± 3.424 | 3.030 ± 2.859 | 4.014 ± 3.644 | 2.972 ± 2.770 | 3.525 ± 3.250 |
| METABOLIC | LPS_0.75 | 3.379 ± 2.987 | 2.611 ± 2.493 | 3.212 ± 2.785 | 2.669 ± 2.364 | 3.019 ± 2.902 |
| METABOLIC | LPS_1.0 | 2.857 ± 2.621 | 2.345 ± 2.264 | 3.027 ± 2.702 | 2.315 ± 2.130 | 2.698 ± 2.531 |
| METABOLIC | LPS_1.25 | 2.564 ± 2.322 | 1.835 ± 1.629 | 2.668 ± 2.400 | 2.066 ± 1.890 | 2.516 ± 2.418 |
| METABOLIC | LPS_1.5 | 2.386 ± 2.189 | 1.988 ± 2.021 | 2.426 ± 2.137 | 1.885 ± 1.779 | 2.123 ± 1.974 |
| METABOLIC | LPS_2.0 | 2.170 ± 2.006 | 2.005 ± 1.866 | 2.090 ± 2.008 | 1.746 ± 1.625 | 2.141 ± 2.057 |

Table S. 15 Mean absolute difference of percentile ranking between PGSs estimated from imputed genotyping data of eight genotyping arrays and six LPS coverages and PGS estimated from WGS in 5 different populations with PRsice p-value setting of 0.0001

| Trait | Array/LPS | AFR | AMR | EAS | EUR | SAS |
|---|---|---|---|---|---|---|
| BMI | GSA | 7.107 ± 6.484 | 3.898 ± 3.463 | 5.900 ± 5.003 | 4.246 ± 3.840 | 4.560 ± 4.194 |
| BMI | JAPONICA | 6.036 ± 5.396 | 3.163 ± 2.837 | 3.676 ± 3.172 | 3.645 ± 3.482 | 3.802 ± 3.367 |
| BMI | UKB_WCSG | 6.271 ± 5.694 | 3.167 ± 2.740 | 4.802 ± 4.052 | 2.881 ± 2.695 | 3.498 ± 3.087 |
| BMI | CYTOSNP | 3.924 ± 3.653 | 2.289 ± 2.253 | 3.490 ± 3.105 | 2.641 ± 2.424 | 2.823 ± 2.560 |
| BMI | PMRA | 5.651 ± 5.050 | 3.547 ± 3.136 | 4.942 ± 4.372 | 3.849 ± 3.421 | 4.352 ± 4.111 |
| BMI | PMDA | 5.470 ± 4.957 | 3.110 ± 2.766 | 5.132 ± 4.545 | 3.550 ± 3.271 | 4.288 ± 3.889 |
| BMI | OMNI2.5 | 2.397 ± 2.217 | 1.666 ± 1.526 | 2.061 ± 2.058 | 1.604 ± 1.635 | 1.634 ± 1.459 |
| BMI | OMNI5 | 1.876 ± 1.777 | 1.221 ± 1.172 | 1.701 ± 1.595 | 1.102 ± 1.044 | 1.477 ± 1.379 |
| BMI | LPS_0.5 | 3.922 ± 3.428 | 3.021 ± 2.740 | 4.475 ± 3.982 | 3.381 ± 3.140 | 3.668 ± 3.220 |
| BMI | LPS_0.75 | 3.355 ± 3.050 | 2.472 ± 2.070 | 3.880 ± 3.437 | 2.858 ± 2.627 | 3.283 ± 2.964 |
| BMI | LPS_1.0 | 3.144 ± 2.868 | 2.107 ± 1.878 | 3.566 ± 3.330 | 2.615 ± 2.501 | 2.816 ± 2.568 |
| BMI | LPS_1.25 | 2.895 ± 2.727 | 2.003 ± 1.817 | 3.245 ± 2.952 | 2.369 ± 2.223 | 2.730 ± 2.289 |
| BMI | LPS_1.5 | 2.723 ± 2.469 | 1.883 ± 1.780 | 2.836 ± 2.769 | 1.956 ± 1.872 | 2.393 ± 2.175 |
| BMI | LPS_2.0 | 2.430 ± 2.279 | 1.752 ± 1.591 | 2.527 ± 2.188 | 1.898 ± 1.759 | 2.187 ± 1.937 |
| DIABETES | GSA | 6.755 ± 5.928 | 3.315 ± 3.284 | 6.001 ± 5.758 | 3.918 ± 3.569 | 4.918 ± 4.598 |
| DIABETES | JAPONICA | 5.815 ± 5.265 | 2.806 ± 2.588 | 3.830 ± 3.660 | 3.279 ± 3.050 | 3.973 ± 3.585 |
| DIABETES | UKB_WCSG | 6.041 ± 5.379 | 2.889 ± 2.652 | 4.434 ± 3.969 | 2.812 ± 2.935 | 3.811 ± 3.447 |
| DIABETES | CYTOSNP | 4.000 ± 3.652 | 2.163 ± 2.149 | 3.536 ± 3.437 | 2.358 ± 2.324 | 2.863 ± 2.791 |
| DIABETES | PMRA | 5.548 ± 5.077 | 3.043 ± 2.593 | 5.219 ± 5.015 | 3.650 ± 3.511 | 4.559 ± 4.047 |
| DIABETES | PMDA | 5.315 ± 4.804 | 3.135 ± 2.645 | 5.053 ± 4.684 | 3.328 ± 3.131 | 4.624 ± 4.157 |
| DIABETES | OMNI2.5 | 2.245 ± 2.055 | 1.616 ± 1.574 | 2.492 ± 2.438 | 1.390 ± 1.416 | 1.846 ± 1.894 |
| DIABETES | OMNI5 | 1.960 ± 1.901 | 1.228 ± 1.138 | 2.066 ± 1.995 | 1.152 ± 1.240 | 1.483 ± 1.386 |

| | | | | | | |
|---|---|---|---|---|---|---|
| DIABETES | LPS_0.5 | 3.943 ± 3.688 | 3.041 ± 2.781 | 4.854 ± 4.568 | 3.499 ± 3.282 | 4.193 ± 3.484 |
| DIABETES | LPS_0.75 | 3.099 ± 2.927 | 2.771 ± 2.248 | 4.134 ± 3.827 | 3.080 ± 2.865 | 3.471 ± 3.142 |
| DIABETES | LPS_1.0 | 3.022 ± 2.798 | 2.325 ± 2.148 | 3.459 ± 3.258 | 2.901 ± 2.874 | 3.123 ± 2.565 |
| DIABETES | LPS_1.25 | 2.740 ± 2.444 | 1.913 ± 1.772 | 3.435 ± 3.177 | 2.522 ± 2.411 | 2.739 ± 2.462 |
| DIABETES | LPS_1.5 | 2.522 ± 2.267 | 2.104 ± 2.027 | 2.997 ± 2.835 | 2.322 ± 2.275 | 2.714 ± 2.344 |
| DIABETES | LPS_2.0 | 2.172 ± 2.107 | 1.867 ± 1.658 | 2.533 ± 2.478 | 2.270 ± 2.220 | 2.649 ± 2.255 |
| HEIGHT | GSA | 7.474 ± 7.015 | 4.132 ± 3.956 | 6.012 ± 5.397 | 3.991 ± 3.758 | 5.359 ± 4.719 |
| HEIGHT | JAPONICA | 6.554 ± 6.103 | 3.873 ± 3.736 | 3.758 ± 3.449 | 3.604 ± 3.295 | 4.386 ± 4.005 |
| HEIGHT | UKB_WCSG | 6.732 ± 6.048 | 3.103 ± 2.633 | 4.904 ± 4.279 | 2.298 ± 2.224 | 3.612 ± 3.290 |
| HEIGHT | CYTOSNP | 4.193 ± 4.017 | 2.707 ± 2.468 | 3.423 ± 2.992 | 2.485 ± 2.238 | 3.088 ± 2.860 |
| HEIGHT | PMRA | 6.056 ± 6.004 | 3.592 ± 3.446 | 4.949 ± 4.351 | 3.563 ± 3.174 | 4.463 ± 4.062 |
| HEIGHT | PMDA | 5.563 ± 5.215 | 3.755 ± 3.595 | 5.267 ± 4.668 | 3.129 ± 2.977 | 4.526 ± 4.199 |
| HEIGHT | OMNI2.5 | 2.345 ± 2.220 | 1.732 ± 1.769 | 2.192 ± 1.961 | 1.463 ± 1.427 | 2.081 ± 1.947 |
| HEIGHT | OMNI5 | 1.943 ± 1.815 | 1.212 ± 1.186 | 1.775 ± 1.684 | 1.013 ± 1.032 | 1.567 ± 1.455 |
| HEIGHT | LPS_0.5 | 4.411 ± 4.128 | 4.004 ± 3.540 | 4.915 ± 4.465 | 3.137 ± 2.853 | 4.299 ± 3.881 |
| HEIGHT | LPS_0.75 | 3.976 ± 3.874 | 3.328 ± 3.013 | 4.485 ± 3.943 | 2.893 ± 2.611 | 3.710 ± 3.189 |
| HEIGHT | LPS_1.0 | 3.446 ± 3.279 | 3.086 ± 2.853 | 4.108 ± 3.622 | 2.602 ± 2.362 | 3.418 ± 3.127 |
| HEIGHT | LPS_1.25 | 3.387 ± 3.063 | 2.711 ± 2.691 | 3.512 ± 3.276 | 2.413 ± 2.167 | 3.103 ± 2.667 |
| HEIGHT | LPS_1.5 | 3.087 ± 3.061 | 2.731 ± 2.614 | 3.331 ± 3.208 | 2.153 ± 1.991 | 3.127 ± 2.783 |
| HEIGHT | LPS_2.0 | 2.819 ± 2.694 | 2.561 ± 2.481 | 2.986 ± 2.875 | 2.088 ± 1.861 | 2.784 ± 2.525 |
| METABOLIC | GSA | 7.323 ± 6.255 | 3.965 ± 3.740 | 5.593 ± 4.984 | 4.057 ± 3.737 | 4.849 ± 4.607 |
| METABOLIC | JAPONICA | 5.941 ± 5.547 | 3.370 ± 3.351 | 3.766 ± 3.331 | 3.680 ± 3.269 | 4.287 ± 3.904 |
| METABOLIC | UKB_WCSG | 6.811 ± 6.301 | 3.197 ± 3.236 | 4.730 ± 4.212 | 2.847 ± 2.680 | 3.644 ± 3.443 |
| METABOLIC | CYTOSNP | 3.763 ± 3.616 | 2.292 ± 2.039 | 3.026 ± 2.594 | 2.430 ± 2.248 | 2.624 ± 2.345 |
| METABOLIC | PMRA | 5.958 ± | 4.016 ± | 4.894 ± | 3.543 ± | 4.512 ± |

| | | | | | | |
|---|---|---|---|---|---|---|
| | | 5.855 | 3.723 | 4.372 | 3.408 | 3.996 |
| METABOLIC | PMDA | 5.874 ± 5.307 | 3.280 ± 3.147 | 5.128 ± 4.464 | 3.363 ± 2.921 | 4.504 ± 4.099 |
| METABOLIC | OMNI2.5 | 2.474 ± 2.284 | 1.832 ± 1.641 | 2.425 ± 2.069 | 1.807 ± 1.699 | 2.070 ± 1.776 |
| METABOLIC | OMNI5 | 1.811 ± 1.695 | 1.173 ± 1.171 | 1.649 ± 1.539 | 1.145 ± 1.151 | 1.394 ± 1.243 |
| METABOLIC | LPS_0.5 | 3.938 ± 3.785 | 3.048 ± 2.933 | 4.613 ± 4.257 | 3.073 ± 2.976 | 3.954 ± 3.439 |
| METABOLIC | LPS_0.75 | 3.566 ± 3.415 | 2.721 ± 2.627 | 3.642 ± 3.192 | 2.674 ± 2.477 | 3.213 ± 3.011 |
| METABOLIC | LPS_1.0 | 3.001 ± 2.768 | 2.287 ± 2.388 | 3.149 ± 2.968 | 2.422 ± 2.261 | 2.977 ± 2.802 |
| METABOLIC | LPS_1.25 | 2.692 ± 2.494 | 1.904 ± 1.845 | 2.902 ± 2.612 | 2.136 ± 2.058 | 2.627 ± 2.505 |
| METABOLIC | LPS_1.5 | 2.584 ± 2.436 | 2.045 ± 2.063 | 2.690 ± 2.462 | 2.050 ± 1.870 | 2.235 ± 2.125 |
| METABOLIC | LPS_2.0 | 2.251 ± 2.170 | 2.015 ± 1.946 | 2.320 ± 2.056 | 1.768 ± 1.636 | 2.253 ± 2.048 |

Table S. 16 Mean absolute difference of percentile ranking between PGSs estimated from imputed genotyping data of eight genotyping arrays and six LPS coverages and PGS estimated from WGS in 5 different populations with PRsice p-value setting of 0.001

| Trait | Array/LPS | AFR | AMR | EAS | EUR | SAS |
|---|---|---|---|---|---|---|
| BMI | GSA | 7.310 ± 6.490 | 4.021 ± 3.617 | 6.067 ± 5.287 | 4.665 ± 4.316 | 4.811 ± 4.265 |
| BMI | JAPONICA | 5.903 ± 5.408 | 3.347 ± 3.292 | 3.813 ± 3.383 | 4.007 ± 3.626 | 4.177 ± 3.789 |
| BMI | UKB_WCSG | 6.370 ± 5.767 | 3.526 ± 3.340 | 5.400 ± 4.544 | 2.901 ± 2.677 | 3.579 ± 3.405 |
| BMI | CYTOSNP | 4.024 ± 3.794 | 2.637 ± 2.512 | 3.854 ± 3.516 | 2.960 ± 2.817 | 3.029 ± 2.596 |
| BMI | PMRA | 5.806 ± 5.098 | 3.802 ± 3.464 | 5.267 ± 4.947 | 3.993 ± 3.416 | 4.718 ± 4.325 |
| BMI | PMDA | 5.723 ± 5.277 | 3.398 ± 3.456 | 5.694 ± 4.759 | 3.659 ± 3.351 | 4.473 ± 4.001 |
| BMI | OMNI2.5 | 2.512 ± 2.251 | 1.850 ± 1.876 | 2.475 ± 2.189 | 1.655 ± 1.742 | 1.808 ± 1.579 |
| BMI | OMNI5 | 2.105 ± 1.854 | 1.319 ± 1.263 | 1.844 ± 1.752 | 1.117 ± 1.113 | 1.462 ± 1.302 |
| BMI | LPS_0.5 | 4.061 ± 3.549 | 3.318 ± 3.329 | 4.827 ± 4.267 | 3.739 ± 3.275 | 3.993 ± 3.484 |
| BMI | LPS_0.75 | 3.664 ± 3.206 | 2.627 ± 2.341 | 4.289 ± 3.625 | 3.098 ± 2.897 | 3.461 ± 3.089 |
| BMI | LPS_1.0 | 3.352 ± 3.037 | 2.496 ± 2.287 | 4.027 ± 3.617 | 2.749 ± 2.637 | 3.021 ± 2.705 |
| BMI | LPS_1.25 | 3.066 ± 2.828 | 2.310 ± 2.223 | 3.562 ± 3.071 | 2.581 ± 2.333 | 2.764 ± 2.441 |
| BMI | LPS_1.5 | 2.916 ± 2.483 | 2.195 ± 2.259 | 3.192 ± 2.831 | 2.243 ± 2.060 | 2.523 ± 2.230 |
| BMI | LPS_2.0 | 2.471 ± 2.294 | 1.944 ± 2.115 | 2.752 ± 2.291 | 2.051 ± 1.923 | 2.308 ± 2.054 |
| DIABETES | GSA | 6.997 ± 6.069 | 3.889 ± 3.735 | 6.354 ± 5.472 | 4.794 ± 4.359 | 5.108 ± 4.707 |
| DIABETES | JAPONICA | 6.390 ± 5.696 | 2.943 ± 2.700 | 3.924 ± 3.718 | 3.583 ± 3.220 | 4.355 ± 3.703 |
| DIABETES | UKB_WCSG | 6.287 ± 5.542 | 3.115 ± 2.824 | 4.913 ± 4.353 | 3.033 ± 2.980 | 4.248 ± 3.795 |
| DIABETES | CYTOSNP | 4.211 ± 3.972 | 2.229 ± 2.089 | 3.847 ± 3.770 | 2.793 ± 2.594 | 3.273 ± 2.873 |
| DIABETES | PMRA | 5.878 ± 5.251 | 3.272 ± 2.912 | 5.161 ± 4.835 | 3.751 ± 3.578 | 5.065 ± 4.403 |
| DIABETES | PMDA | 5.514 ± 4.818 | 3.282 ± 2.891 | 5.382 ± 4.839 | 3.432 ± 3.456 | 4.651 ± 4.204 |
| DIABETES | OMNI2.5 | 2.414 ± 2.342 | 1.521 ± 1.525 | 2.502 ± 2.461 | 1.685 ± 1.584 | 2.096 ± 1.990 |
| DIABETES | OMNI5 | 2.139 ± 2.106 | 1.195 ± 1.096 | 2.037 ± 2.083 | 1.304 ± 1.279 | 1.682 ± 1.579 |

| | | | | | | |
|---|---|---|---|---|---|---|
| DIABETES | LPS_0.5 | 4.200 ± 3.845 | 2.891 ± 2.396 | 4.693 ± 4.149 | 3.527 ± 3.381 | 4.344 ± 3.763 |
| DIABETES | LPS_0.75 | 3.414 ± 3.164 | 2.687 ± 2.266 | 4.124 ± 3.717 | 2.965 ± 2.844 | 3.758 ± 3.432 |
| DIABETES | LPS_1.0 | 3.052 ± 2.877 | 2.382 ± 2.090 | 3.405 ± 3.068 | 2.838 ± 2.683 | 3.258 ± 2.967 |
| DIABETES | LPS_1.25 | 2.725 ± 2.720 | 2.187 ± 1.896 | 3.281 ± 3.101 | 2.531 ± 2.277 | 2.768 ± 2.488 |
| DIABETES | LPS_1.5 | 2.628 ± 2.490 | 2.055 ± 1.847 | 2.950 ± 2.795 | 2.511 ± 2.298 | 2.900 ± 2.674 |
| DIABETES | LPS_2.0 | 2.348 ± 2.213 | 1.761 ± 1.637 | 2.604 ± 2.379 | 2.322 ± 2.216 | 2.645 ± 2.446 |
| HEIGHT | GSA | 7.661 ± 7.209 | 3.944 ± 3.599 | 6.063 ± 5.562 | 3.880 ± 3.467 | 5.469 ± 4.911 |
| HEIGHT | JAPONICA | 6.723 ± 5.845 | 3.736 ± 3.479 | 3.792 ± 3.659 | 3.572 ± 3.233 | 4.741 ± 4.626 |
| HEIGHT | UKB_WCSG | 6.752 ± 6.097 | 2.969 ± 2.556 | 5.021 ± 4.507 | 2.186 ± 2.026 | 3.688 ± 3.571 |
| HEIGHT | CYTOSNP | 4.280 ± 3.950 | 2.799 ± 2.642 | 3.311 ± 3.142 | 2.499 ± 2.320 | 3.134 ± 3.033 |
| HEIGHT | PMRA | 6.201 ± 6.068 | 3.620 ± 3.345 | 5.017 ± 4.399 | 3.365 ± 3.063 | 4.763 ± 4.370 |
| HEIGHT | PMDA | 5.698 ± 5.345 | 3.677 ± 3.614 | 5.143 ± 4.839 | 3.065 ± 2.778 | 4.490 ± 4.384 |
| HEIGHT | OMNI2.5 | 2.376 ± 2.206 | 1.717 ± 1.623 | 2.235 ± 2.038 | 1.449 ± 1.414 | 2.111 ± 2.019 |
| HEIGHT | OMNI5 | 1.974 ± 1.868 | 1.195 ± 1.177 | 1.939 ± 1.819 | 0.962 ± 0.868 | 1.573 ± 1.547 |
| HEIGHT | LPS_0.5 | 4.309 ± 4.155 | 3.773 ± 3.713 | 5.049 ± 4.836 | 3.138 ± 2.670 | 4.541 ± 4.013 |
| HEIGHT | LPS_0.75 | 3.904 ± 3.845 | 3.337 ± 3.053 | 4.451 ± 3.938 | 2.882 ± 2.530 | 3.750 ± 3.493 |
| HEIGHT | LPS_1.0 | 3.479 ± 3.220 | 2.765 ± 2.685 | 4.110 ± 3.793 | 2.467 ± 2.252 | 3.463 ± 3.203 |
| HEIGHT | LPS_1.25 | 3.341 ± 3.131 | 2.659 ± 2.680 | 3.588 ± 3.455 | 2.325 ± 2.108 | 3.130 ± 2.921 |
| HEIGHT | LPS_1.5 | 3.051 ± 3.107 | 2.629 ± 2.402 | 3.269 ± 3.264 | 2.165 ± 1.927 | 3.196 ± 2.983 |
| HEIGHT | LPS_2.0 | 2.928 ± 2.746 | 2.296 ± 2.289 | 2.981 ± 2.860 | 2.010 ± 1.809 | 2.810 ± 2.642 |
| METABOLIC | GSA | 7.146 ± 6.573 | 4.249 ± 4.086 | 5.975 ± 5.555 | 4.191 ± 3.908 | 5.215 ± 4.638 |
| METABOLIC | JAPONICA | 5.779 ± 5.359 | 3.513 ± 3.234 | 4.009 ± 3.430 | 3.813 ± 3.234 | 4.310 ± 4.041 |
| METABOLIC | UKB_WCSG | 6.687 ± 6.409 | 3.249 ± 2.982 | 5.135 ± 4.658 | 2.805 ± 2.471 | 3.737 ± 3.487 |
| METABOLIC | CYTOSNP | 3.893 ± 3.561 | 2.398 ± 2.274 | 3.300 ± 2.834 | 2.449 ± 2.281 | 2.953 ± 2.788 |
| METABOLIC | PMRA | 5.859 ± | 3.998 ± | 5.416 ± | 3.601 ± | 4.543 ± |

| | | | | | | |
|---|---|---|---|---|---|---|
| | | 5.504 | 3.312 | 4.889 | 3.328 | 4.159 |
| METABOLIC | PMDA | 5.612 ± 4.945 | 3.310 ± 3.122 | 5.518 ± 4.883 | 3.302 ± 3.102 | 4.507 ± 4.277 |
| METABOLIC | OMNI2.5 | 2.407 ± 2.294 | 1.912 ± 1.727 | 2.552 ± 2.260 | 1.847 ± 1.747 | 2.245 ± 2.070 |
| METABOLIC | OMNI5 | 1.769 ± 1.695 | 1.287 ± 1.269 | 1.788 ± 1.727 | 1.151 ± 1.077 | 1.536 ± 1.497 |
| METABOLIC | LPS_0.5 | 4.097 ± 3.814 | 3.113 ± 2.918 | 5.100 ± 4.458 | 3.141 ± 2.938 | 4.331 ± 3.922 |
| METABOLIC | LPS_0.75 | 3.635 ± 3.370 | 2.614 ± 2.437 | 4.027 ± 3.706 | 2.605 ± 2.390 | 3.447 ± 3.190 |
| METABOLIC | LPS_1.0 | 2.967 ± 2.935 | 2.284 ± 2.213 | 3.549 ± 3.197 | 2.417 ± 2.230 | 3.172 ± 3.177 |
| METABOLIC | LPS_1.25 | 2.807 ± 2.651 | 2.033 ± 2.025 | 3.355 ± 2.943 | 2.102 ± 2.149 | 2.845 ± 2.566 |
| METABOLIC | LPS_1.5 | 2.630 ± 2.394 | 2.186 ± 2.030 | 3.124 ± 2.872 | 2.020 ± 1.835 | 2.601 ± 2.533 |
| METABOLIC | LPS_2.0 | 2.300 ± 2.113 | 1.917 ± 1.701 | 2.478 ± 2.270 | 1.820 ± 1.669 | 2.475 ± 2.288 |

Table S. 17 Mean absolute difference of percentile ranking between PGSs estimated from imputed genotyping data of eight genotyping arrays and six LPS coverages and PGS estimated from WGS in 5 different populations with PRsice p-value setting of 0.01

| Trait | Array/LPS | AFR | AMR | EAS | EUR | SAS |
|---|---|---|---|---|---|---|
| BMI | GSA | 7.166 ± 6.461 | 3.868 ± 3.437 | 6.544 ± 5.648 | 4.647 ± 4.168 | 5.129 ± 4.471 |
| BMI | JAPONICA | 6.157 ± 5.748 | 3.259 ± 2.937 | 4.111 ± 3.709 | 4.091 ± 3.764 | 4.649 ± 3.886 |
| BMI | UKB_WCSG | 6.398 ± 5.879 | 3.168 ± 2.807 | 5.790 ± 4.822 | 2.982 ± 2.795 | 3.581 ± 3.238 |
| BMI | CYTOSNP | 4.105 ± 3.947 | 2.535 ± 2.450 | 4.041 ± 3.588 | 3.116 ± 2.883 | 3.419 ± 2.797 |
| BMI | PMRA | 6.126 ± 5.827 | 3.553 ± 3.019 | 5.814 ± 5.109 | 4.186 ± 3.919 | 5.016 ± 4.509 |
| BMI | PMDA | 5.740 ± 5.256 | 3.306 ± 3.095 | 6.026 ± 5.074 | 3.775 ± 3.478 | 4.218 ± 3.696 |
| BMI | OMNI2.5 | 2.399 ± 2.202 | 1.680 ± 1.525 | 2.594 ± 2.258 | 1.793 ± 1.766 | 2.146 ± 1.866 |
| BMI | OMNI5 | 1.977 ± 1.846 | 1.149 ± 1.075 | 1.955 ± 1.774 | 1.260 ± 1.167 | 1.598 ± 1.375 |
| BMI | LPS_0.5 | 4.341 ± 4.119 | 3.491 ± 3.215 | 5.294 ± 4.784 | 3.845 ± 3.333 | 4.154 ± 3.887 |
| BMI | LPS_0.75 | 3.730 ± 3.530 | 2.888 ± 2.538 | 4.625 ± 3.911 | 3.218 ± 2.923 | 3.780 ± 3.458 |
| BMI | LPS_1.0 | 3.402 ± 3.327 | 2.532 ± 2.427 | 4.248 ± 3.631 | 2.860 ± 2.650 | 3.097 ± 2.795 |
| BMI | LPS_1.25 | 3.078 ± 3.004 | 2.170 ± 2.084 | 3.892 ± 3.384 | 2.800 ± 2.652 | 2.950 ± 2.552 |
| BMI | LPS_1.5 | 2.924 ± 2.786 | 2.116 ± 1.847 | 3.312 ± 2.929 | 2.425 ± 2.104 | 2.707 ± 2.469 |
| BMI | LPS_2.0 | 2.543 ± 2.533 | 1.971 ± 1.788 | 2.880 ± 2.436 | 2.165 ± 2.033 | 2.495 ± 2.345 |
| DIABETES | GSA | 7.176 ± 6.712 | 4.116 ± 3.573 | 6.837 ± 5.936 | 5.331 ± 4.699 | 5.265 ± 4.631 |
| DIABETES | JAPONICA | 6.690 ± 6.290 | 3.285 ± 3.034 | 4.035 ± 4.048 | 4.250 ± 3.986 | 4.572 ± 4.063 |
| DIABETES | UKB_WCSG | 6.633 ± 6.334 | 3.305 ± 2.771 | 5.496 ± 4.687 | 3.421 ± 3.456 | 4.522 ± 4.063 |
| DIABETES | CYTOSNP | 4.440 ± 4.319 | 2.344 ± 2.179 | 3.980 ± 3.600 | 2.886 ± 2.684 | 3.558 ± 3.250 |
| DIABETES | PMRA | 5.877 ± 5.343 | 3.692 ± 2.981 | 5.842 ± 5.068 | 4.201 ± 4.024 | 5.263 ± 4.759 |
| DIABETES | PMDA | 5.720 ± 5.027 | 3.244 ± 2.935 | 5.519 ± 5.030 | 3.894 ± 3.595 | 4.889 ± 4.234 |
| DIABETES | OMNI2.5 | 2.584 ± 2.393 | 1.575 ± 1.454 | 2.678 ± 2.387 | 1.969 ± 1.854 | 2.129 ± 1.910 |
| DIABETES | OMNI5 | 2.372 ± 2.211 | 1.198 ± 1.099 | 2.159 ± 2.007 | 1.477 ± 1.393 | 1.782 ± 1.616 |

| | | | | | | |
|---|---|---|---|---|---|---|
| DIABETES | LPS_0.5 | 4.206 ± 3.926 | 3.204 ± 2.819 | 5.106 ± 4.479 | 4.132 ± 3.590 | 4.332 ± 3.848 |
| DIABETES | LPS_0.75 | 3.642 ± 3.345 | 2.643 ± 2.476 | 4.401 ± 3.798 | 3.405 ± 3.004 | 4.031 ± 3.490 |
| DIABETES | LPS_1.0 | 3.200 ± 3.037 | 2.433 ± 2.031 | 3.990 ± 3.467 | 3.148 ± 2.861 | 3.486 ± 3.031 |
| DIABETES | LPS_1.25 | 3.019 ± 2.910 | 2.292 ± 2.103 | 3.519 ± 3.160 | 2.927 ± 2.608 | 2.986 ± 2.668 |
| DIABETES | LPS_1.5 | 2.857 ± 2.558 | 2.104 ± 1.902 | 3.320 ± 2.859 | 2.775 ± 2.518 | 3.152 ± 2.682 |
| DIABETES | LPS_2.0 | 2.505 ± 2.361 | 1.949 ± 1.807 | 2.961 ± 2.659 | 2.420 ± 2.234 | 2.724 ± 2.538 |
| HEIGHT | GSA | 7.764 ± 7.260 | 4.155 ± 3.585 | 5.850 ± 5.374 | 3.913 ± 3.683 | 5.527 ± 5.001 |
| HEIGHT | JAPONICA | 6.539 ± 6.004 | 3.804 ± 3.514 | 3.915 ± 3.530 | 3.342 ± 3.080 | 4.752 ± 4.537 |
| HEIGHT | UKB_WCSG | 6.809 ± 6.253 | 2.903 ± 2.643 | 4.837 ± 4.265 | 2.203 ± 2.037 | 3.879 ± 3.536 |
| HEIGHT | CYTOSNP | 4.235 ± 3.827 | 2.654 ± 2.299 | 3.455 ± 3.134 | 2.463 ± 2.343 | 3.419 ± 3.196 |
| HEIGHT | PMRA | 6.274 ± 6.094 | 3.808 ± 3.511 | 5.090 ± 4.365 | 3.331 ± 3.083 | 4.822 ± 4.459 |
| HEIGHT | PMDA | 5.719 ± 5.178 | 3.524 ± 3.491 | 5.427 ± 4.854 | 2.979 ± 2.832 | 4.534 ± 4.163 |
| HEIGHT | OMNI2.5 | 2.392 ± 2.194 | 1.689 ± 1.503 | 2.356 ± 1.961 | 1.498 ± 1.459 | 2.079 ± 1.979 |
| HEIGHT | OMNI5 | 2.021 ± 1.897 | 1.174 ± 1.049 | 1.873 ± 1.807 | 0.989 ± 0.992 | 1.566 ± 1.475 |
| HEIGHT | LPS_0.5 | 4.518 ± 4.340 | 3.702 ± 3.355 | 5.035 ± 4.712 | 3.104 ± 2.847 | 4.511 ± 4.027 |
| HEIGHT | LPS_0.75 | 4.082 ± 4.034 | 3.235 ± 2.955 | 4.427 ± 3.901 | 2.971 ± 2.582 | 3.883 ± 3.712 |
| HEIGHT | LPS_1.0 | 3.720 ± 3.434 | 2.663 ± 2.356 | 4.204 ± 3.697 | 2.456 ± 2.362 | 3.348 ± 2.951 |
| HEIGHT | LPS_1.25 | 3.611 ± 3.274 | 2.620 ± 2.429 | 3.633 ± 3.352 | 2.318 ± 2.111 | 3.239 ± 2.929 |
| HEIGHT | LPS_1.5 | 3.319 ± 3.169 | 2.531 ± 2.422 | 3.344 ± 3.073 | 2.088 ± 1.997 | 3.103 ± 2.843 |
| HEIGHT | LPS_2.0 | 3.046 ± 2.861 | 2.293 ± 2.109 | 3.055 ± 2.728 | 2.036 ± 1.902 | 2.734 ± 2.521 |
| METABOLIC | GSA | 7.587 ± 7.280 | 4.001 ± 3.504 | 6.458 ± 6.012 | 4.190 ± 3.712 | 5.581 ± 4.695 |
| METABOLIC | JAPONICA | 6.055 ± 5.785 | 3.377 ± 3.100 | 4.061 ± 3.628 | 4.009 ± 3.689 | 4.249 ± 3.792 |
| METABOLIC | UKB_WCSG | 6.828 ± 6.807 | 3.036 ± 2.740 | 5.720 ± 4.899 | 2.863 ± 2.619 | 3.925 ± 3.592 |
| METABOLIC | CYTOSNP | 4.050 ± 3.767 | 2.536 ± 2.299 | 3.612 ± 3.096 | 2.511 ± 2.382 | 3.107 ± 2.764 |
| METABOLIC | PMRA | 6.076 ± | 3.751 ± | 5.930 ± | 3.729 ± | 4.680 ± |

| | | | | | | |
|---|---|---|---|---|---|---|
| | | 5.624 | 3.360 | 5.385 | 3.296 | 4.350 |
| METABOLIC | PMDA | 5.711 ± 5.172 | 3.217 ± 2.993 | 5.875 ± 5.016 | 3.525 ± 3.171 | 4.504 ± 4.215 |
| METABOLIC | OMNI2.5 | 2.544 ± 2.516 | 1.810 ± 1.639 | 2.644 ± 2.298 | 1.889 ± 1.690 | 2.232 ± 2.037 |
| METABOLIC | OMNI5 | 1.743 ± 1.725 | 1.174 ± 1.064 | 1.780 ± 1.657 | 1.100 ± 1.057 | 1.546 ± 1.488 |
| METABOLIC | LPS_0.5 | 4.254 ± 4.322 | 3.050 ± 2.814 | 5.440 ± 4.935 | 3.499 ± 3.289 | 4.476 ± 4.072 |
| METABOLIC | LPS_0.75 | 3.816 ± 3.571 | 2.676 ± 2.461 | 4.262 ± 3.762 | 2.693 ± 2.334 | 3.699 ± 3.368 |
| METABOLIC | LPS_1.0 | 3.301 ± 3.276 | 2.266 ± 1.965 | 3.659 ± 3.300 | 2.673 ± 2.317 | 3.049 ± 2.877 |
| METABOLIC | LPS_1.25 | 3.150 ± 2.999 | 2.188 ± 1.937 | 3.591 ± 3.074 | 2.375 ± 2.221 | 2.910 ± 2.646 |
| METABOLIC | LPS_1.5 | 2.871 ± 2.678 | 2.319 ± 2.184 | 3.234 ± 2.906 | 2.234 ± 1.931 | 2.702 ± 2.434 |
| METABOLIC | LPS_2.0 | 2.604 ± 2.562 | 1.879 ± 1.671 | 2.671 ± 2.468 | 1.979 ± 1.766 | 2.451 ± 2.395 |

Table S. 18 Mean absolute difference of percentile ranking between PGSs estimated from imputed genotyping data of eight genotyping arrays and six LPS coverages and PGS estimated from WGS in 5 different populations with PRsice p-value setting of 0.1

| Trait | Array/LPS | AFR | AMR | EAS | EUR | SAS |
|---|---|---|---|---|---|---|
| BMI | GSA | 7.444 ± 6.721 | 4.658 ± 4.285 | 7.156 ± 6.495 | 5.107 ± 4.593 | 5.697 ± 4.929 |
| BMI | JAPONICA | 6.339 ± 5.937 | 4.229 ± 3.936 | 4.542 ± 3.866 | 4.374 ± 4.045 | 4.901 ± 4.236 |
| BMI | UKB_WCSG | 7.196 ± 6.703 | 3.361 ± 2.959 | 6.019 ± 5.391 | 3.077 ± 2.942 | 4.205 ± 3.639 |
| BMI | CYTOSNP | 4.328 ± 4.262 | 2.905 ± 2.571 | 4.331 ± 3.826 | 3.599 ± 3.449 | 4.072 ± 3.540 |
| BMI | PMRA | 6.383 ± 6.222 | 4.045 ± 3.615 | 6.291 ± 5.857 | 4.806 ± 4.194 | 5.516 ± 5.146 |
| BMI | PMDA | 6.082 ± 5.685 | 3.763 ± 3.218 | 6.067 ± 5.530 | 4.378 ± 4.045 | 4.887 ± 4.155 |
| BMI | OMNI2.5 | 2.544 ± 2.481 | 1.994 ± 1.745 | 2.870 ± 2.370 | 1.968 ± 1.890 | 2.274 ± 2.052 |
| BMI | OMNI5 | 2.041 ± 1.912 | 1.355 ± 1.179 | 2.049 ± 1.834 | 1.281 ± 1.280 | 1.618 ± 1.441 |
| BMI | LPS_0.5 | 4.901 ± 4.721 | 4.012 ± 3.706 | 5.615 ± 5.037 | 4.265 ± 4.054 | 4.756 ± 4.176 |
| BMI | LPS_0.75 | 4.035 ± 3.967 | 3.481 ± 3.164 | 5.270 ± 4.721 | 3.648 ± 3.195 | 4.144 ± 3.656 |
| BMI | LPS_1.0 | 3.653 ± 3.453 | 2.750 ± 2.672 | 4.657 ± 4.024 | 3.261 ± 3.016 | 3.658 ± 3.210 |
| BMI | LPS_1.25 | 3.500 ± 3.174 | 2.829 ± 2.497 | 4.110 ± 3.812 | 3.136 ± 2.978 | 3.203 ± 2.908 |
| BMI | LPS_1.5 | 3.178 ± 3.030 | 2.537 ± 2.330 | 3.637 ± 3.151 | 2.732 ± 2.536 | 3.180 ± 2.790 |
| BMI | LPS_2.0 | 2.756 ± 2.561 | 2.335 ± 2.271 | 3.382 ± 2.905 | 2.354 ± 2.105 | 2.804 ± 2.412 |
| DIABETES | GSA | 7.440 ± 6.979 | 4.300 ± 3.759 | 7.583 ± 6.828 | 4.795 ± 4.541 | 5.544 ± 5.203 |
| DIABETES | JAPONICA | 6.464 ± 6.270 | 3.311 ± 2.951 | 4.325 ± 4.347 | 3.985 ± 3.460 | 4.620 ± 4.155 |
| DIABETES | UKB_WCSG | 6.799 ± 6.650 | 3.128 ± 2.637 | 6.286 ± 5.637 | 3.189 ± 2.979 | 4.479 ± 4.254 |
| DIABETES | CYTOSNP | 4.532 ± 4.402 | 2.377 ± 2.078 | 4.364 ± 3.896 | 2.797 ± 2.458 | 3.706 ± 3.419 |
| DIABETES | PMRA | 6.223 ± 5.856 | 3.800 ± 3.366 | 6.467 ± 5.876 | 4.169 ± 3.740 | 5.314 ± 4.919 |
| DIABETES | PMDA | 5.832 ± 5.668 | 3.323 ± 3.024 | 6.386 ± 5.726 | 3.476 ± 3.220 | 4.950 ± 4.668 |
| DIABETES | OMNI2.5 | 2.662 ± 2.347 | 1.573 ± 1.448 | 2.940 ± 2.563 | 1.732 ± 1.588 | 2.413 ± 2.174 |
| DIABETES | OMNI5 | 2.383 ± 2.248 | 1.229 ± 1.166 | 2.378 ± 2.180 | 1.302 ± 1.256 | 1.967 ± 1.858 |

| | | | | | | |
|---|---|---|---|---|---|---|
| DIABETES | LPS_0.5 | 4.580 ± 4.436 | 3.453 ± 2.836 | 5.771 ± 5.135 | 3.958 ± 3.557 | 4.426 ± 4.247 |
| DIABETES | LPS_0.75 | 4.002 ± 3.777 | 3.119 ± 2.823 | 4.770 ± 4.188 | 3.241 ± 2.771 | 4.135 ± 3.874 |
| DIABETES | LPS_1.0 | 3.526 ± 3.307 | 2.393 ± 2.279 | 4.274 ± 4.049 | 2.942 ± 2.864 | 3.445 ± 3.201 |
| DIABETES | LPS_1.25 | 3.338 ± 3.265 | 2.444 ± 2.134 | 3.914 ± 3.502 | 2.677 ± 2.261 | 3.010 ± 2.822 |
| DIABETES | LPS_1.5 | 3.266 ± 3.240 | 1.978 ± 1.950 | 3.629 ± 3.250 | 2.567 ± 2.343 | 3.144 ± 2.938 |
| DIABETES | LPS_2.0 | 2.698 ± 2.544 | 2.021 ± 1.826 | 3.228 ± 2.920 | 2.239 ± 2.006 | 2.760 ± 2.583 |
| HEIGHT | GSA | 7.639 ± 6.802 | 4.024 ± 3.630 | 5.952 ± 5.233 | 3.688 ± 3.342 | 5.759 ± 5.077 |
| HEIGHT | JAPONICA | 6.483 ± 5.759 | 3.770 ± 3.584 | 4.056 ± 3.597 | 3.205 ± 2.879 | 4.501 ± 4.573 |
| HEIGHT | UKB_WCSG | 6.776 ± 6.017 | 2.924 ± 2.667 | 4.808 ± 4.290 | 2.207 ± 1.961 | 3.986 ± 3.690 |
| HEIGHT | CYTOSNP | 4.447 ± 4.069 | 2.981 ± 2.764 | 3.536 ± 3.220 | 2.340 ± 2.246 | 3.584 ± 3.456 |
| HEIGHT | PMRA | 6.329 ± 6.047 | 3.885 ± 3.710 | 5.036 ± 4.243 | 3.244 ± 2.877 | 4.753 ± 4.615 |
| HEIGHT | PMDA | 5.789 ± 4.944 | 3.620 ± 3.525 | 5.508 ± 5.089 | 2.856 ± 2.706 | 4.569 ± 4.351 |
| HEIGHT | OMNI2.5 | 2.460 ± 2.246 | 1.830 ± 1.682 | 2.329 ± 1.930 | 1.415 ± 1.289 | 2.228 ± 2.137 |
| HEIGHT | OMNI5 | 2.001 ± 1.866 | 1.217 ± 1.156 | 1.795 ± 1.661 | 0.972 ± 0.920 | 1.562 ± 1.437 |
| HEIGHT | LPS_0.5 | 4.913 ± 4.529 | 3.510 ± 3.148 | 5.206 ± 4.948 | 3.060 ± 2.743 | 4.719 ± 4.286 |
| HEIGHT | LPS_0.75 | 4.327 ± 4.075 | 3.215 ± 2.955 | 4.397 ± 3.882 | 2.894 ± 2.542 | 4.001 ± 3.838 |
| HEIGHT | LPS_1.0 | 3.794 ± 3.549 | 2.762 ± 2.439 | 4.191 ± 3.652 | 2.399 ± 2.185 | 3.525 ± 3.182 |
| HEIGHT | LPS_1.25 | 3.737 ± 3.332 | 2.805 ± 2.680 | 3.673 ± 3.316 | 2.253 ± 1.958 | 3.262 ± 3.310 |
| HEIGHT | LPS_1.5 | 3.559 ± 3.316 | 2.457 ± 2.263 | 3.336 ± 3.019 | 1.987 ± 1.845 | 3.159 ± 2.919 |
| HEIGHT | LPS_2.0 | 3.044 ± 2.741 | 2.269 ± 2.127 | 3.196 ± 2.817 | 1.957 ± 1.804 | 2.879 ± 2.638 |
| METABOLIC | GSA | 7.599 ± 6.769 | 4.156 ± 3.730 | 7.074 ± 6.270 | 4.318 ± 3.962 | 5.828 ± 5.151 |
| METABOLIC | JAPONICA | 6.168 ± 5.740 | 3.462 ± 3.282 | 4.240 ± 4.017 | 4.281 ± 3.863 | 4.696 ± 4.155 |
| METABOLIC | UKB_WCSG | 7.247 ± 6.807 | 2.952 ± 2.839 | 6.096 ± 5.153 | 2.937 ± 2.769 | 3.949 ± 3.493 |
| METABOLIC | CYTOSNP | 4.222 ± 3.856 | 2.525 ± 2.205 | 4.119 ± 3.611 | 2.994 ± 2.820 | 3.600 ± 3.128 |
| METABOLIC | PMRA | 5.963 ± | 4.055 ± | 6.393 ± | 3.974 ± | 5.030 ± |

| | | | | | | |
|---|---|---|---|---|---|---|
| | | 5.423 | 3.684 | 5.473 | 3.524 | 4.638 |
| METABOLIC | PMDA | 5.468 ± 4.899 | 3.332 ± 3.157 | 6.177 ± 5.653 | 3.794 ± 3.683 | 4.711 ± 4.066 |
| METABOLIC | OMNI2.5 | 2.464 ± 2.344 | 1.801 ± 1.699 | 2.686 ± 2.569 | 1.926 ± 1.852 | 2.319 ± 2.102 |
| METABOLIC | OMNI5 | 1.816 ± 1.717 | 1.284 ± 1.256 | 1.894 ± 1.760 | 1.173 ± 1.134 | 1.714 ± 1.597 |
| METABOLIC | LPS_0.5 | 4.495 ± 4.143 | 3.508 ± 3.286 | 5.988 ± 5.435 | 3.870 ± 3.647 | 4.726 ± 4.218 |
| METABOLIC | LPS_0.75 | 3.980 ± 3.715 | 2.842 ± 2.674 | 4.763 ± 4.423 | 3.252 ± 3.023 | 3.781 ± 3.385 |
| METABOLIC | LPS_1.0 | 3.427 ± 3.260 | 2.606 ± 2.443 | 4.293 ± 3.850 | 2.975 ± 2.688 | 3.503 ± 3.246 |
| METABOLIC | LPS_1.25 | 3.139 ± 2.911 | 2.329 ± 2.313 | 4.049 ± 3.783 | 2.654 ± 2.575 | 3.291 ± 2.962 |
| METABOLIC | LPS_1.5 | 2.819 ± 2.722 | 2.523 ± 2.352 | 3.741 ± 3.692 | 2.602 ± 2.376 | 3.019 ± 2.725 |
| METABOLIC | LPS_2.0 | 2.623 ± 2.605 | 2.023 ± 1.973 | 3.099 ± 2.929 | 2.183 ± 1.914 | 2.673 ± 2.465 |

Table S. 19 Mean absolute difference of percentile ranking between PGSs estimated from imputed genotyping data of eight genotyping arrays and six LPS coverages and PGS estimated from WGS in 5 different populations with PRsice p-value setting of 0.2

| Trait | Array/LPS | AFR | AMR | EAS | EUR | SAS |
|---|---|---|---|---|---|---|
| BMI | GSA | 7.692 ± 7.002 | 4.600 ± 4.081 | 7.153 ± 6.678 | 5.209 ± 4.644 | 5.891 ± 5.097 |
| BMI | JAPONICA | 6.737 ± 6.455 | 4.092 ± 3.782 | 4.606 ± 3.966 | 4.574 ± 4.069 | 4.984 ± 4.235 |
| BMI | UKB_WCSG | 7.384 ± 6.865 | 3.292 ± 2.857 | 6.106 ± 5.374 | 3.111 ± 2.922 | 4.151 ± 3.559 |
| BMI | CYTOSNP | 4.599 ± 4.478 | 2.982 ± 2.695 | 4.612 ± 3.864 | 3.694 ± 3.321 | 4.175 ± 3.501 |
| BMI | PMRA | 6.685 ± 6.329 | 4.028 ± 3.565 | 6.280 ± 5.949 | 4.781 ± 4.235 | 5.601 ± 4.994 |
| BMI | PMDA | 6.319 ± 5.911 | 3.741 ± 3.180 | 6.054 ± 5.542 | 4.443 ± 4.016 | 5.020 ± 4.165 |
| BMI | OMNI2.5 | 2.621 ± 2.546 | 1.951 ± 1.647 | 2.893 ± 2.446 | 2.042 ± 1.908 | 2.437 ± 2.027 |
| BMI | OMNI5 | 2.106 ± 1.994 | 1.290 ± 1.116 | 2.078 ± 1.862 | 1.368 ± 1.289 | 1.622 ± 1.468 |
| BMI | LPS_0.5 | 5.084 ± 5.093 | 3.858 ± 3.362 | 5.795 ± 5.140 | 4.525 ± 4.062 | 4.777 ± 4.239 |
| BMI | LPS_0.75 | 4.195 ± 4.008 | 3.459 ± 3.185 | 5.443 ± 4.839 | 3.708 ± 3.216 | 4.219 ± 3.742 |
| BMI | LPS_1.0 | 3.773 ± 3.561 | 2.802 ± 2.468 | 4.598 ± 4.042 | 3.222 ± 2.984 | 3.699 ± 3.216 |
| BMI | LPS_1.25 | 3.526 ± 3.244 | 2.780 ± 2.367 | 4.224 ± 3.915 | 3.179 ± 2.870 | 3.245 ± 3.010 |
| BMI | LPS_1.5 | 3.198 ± 3.080 | 2.486 ± 2.311 | 3.762 ± 3.305 | 2.731 ± 2.578 | 3.289 ± 2.947 |
| BMI | LPS_2.0 | 2.875 ± 2.642 | 2.334 ± 2.083 | 3.435 ± 2.923 | 2.396 ± 2.004 | 2.896 ± 2.466 |
| DIABETES | GSA | 7.466 ± 7.119 | 4.339 ± 3.966 | 7.723 ± 7.070 | 4.731 ± 4.380 | 5.692 ± 5.135 |
| DIABETES | JAPONICA | 6.611 ± 6.375 | 3.402 ± 2.945 | 4.416 ± 4.216 | 3.903 ± 3.584 | 4.598 ± 4.211 |
| DIABETES | UKB_WCSG | 7.000 ± 6.852 | 3.101 ± 2.843 | 6.441 ± 5.506 | 3.069 ± 2.769 | 4.558 ± 4.350 |
| DIABETES | CYTOSNP | 4.712 ± 4.752 | 2.407 ± 2.308 | 4.413 ± 3.989 | 2.779 ± 2.450 | 3.645 ± 3.373 |
| DIABETES | PMRA | 6.234 ± 5.796 | 3.682 ± 3.587 | 6.444 ± 5.648 | 3.996 ± 3.721 | 5.417 ± 4.928 |
| DIABETES | PMDA | 5.764 ± 5.489 | 3.384 ± 3.053 | 6.211 ± 5.688 | 3.543 ± 3.112 | 4.756 ± 4.515 |
| DIABETES | OMNI2.5 | 2.513 ± 2.318 | 1.525 ± 1.471 | 2.970 ± 2.596 | 1.732 ± 1.585 | 2.313 ± 2.126 |
| DIABETES | OMNI5 | 2.330 ± 2.189 | 1.238 ± 1.161 | 2.398 ± 2.185 | 1.307 ± 1.233 | 1.951 ± 1.809 |

| | | | | | | |
|---|---|---|---|---|---|---|
| DIABETES | LPS_0.5 | 4.583 ± 4.432 | 3.497 ± 3.119 | 5.887 ± 4.984 | 3.852 ± 3.482 | 4.501 ± 4.122 |
| DIABETES | LPS_0.75 | 3.989 ± 3.944 | 3.186 ± 2.855 | 4.757 ± 4.274 | 3.262 ± 2.859 | 4.079 ± 3.709 |
| DIABETES | LPS_1.0 | 3.738 ± 3.401 | 2.440 ± 2.310 | 4.353 ± 4.184 | 2.852 ± 2.677 | 3.359 ± 3.007 |
| DIABETES | LPS_1.25 | 3.399 ± 3.381 | 2.457 ± 2.177 | 3.832 ± 3.351 | 2.652 ± 2.342 | 3.004 ± 2.691 |
| DIABETES | LPS_1.5 | 3.404 ± 3.248 | 1.910 ± 2.036 | 3.693 ± 3.262 | 2.605 ± 2.300 | 3.080 ± 2.687 |
| DIABETES | LPS_2.0 | 2.804 ± 2.833 | 2.035 ± 1.877 | 3.320 ± 2.974 | 2.227 ± 1.986 | 2.652 ± 2.483 |
| HEIGHT | GSA | 7.859 ± 6.971 | 4.039 ± 3.540 | 5.945 ± 5.191 | 3.624 ± 3.221 | 5.846 ± 5.293 |
| HEIGHT | JAPONICA | 6.395 ± 5.697 | 3.633 ± 3.343 | 4.152 ± 3.582 | 3.201 ± 2.791 | 4.698 ± 4.615 |
| HEIGHT | UKB_WCSG | 6.829 ± 6.104 | 3.020 ± 2.649 | 4.837 ± 4.295 | 2.148 ± 1.928 | 3.982 ± 3.723 |
| HEIGHT | CYTOSNP | 4.477 ± 4.065 | 3.015 ± 2.624 | 3.723 ± 3.285 | 2.381 ± 2.266 | 3.577 ± 3.485 |
| HEIGHT | PMRA | 6.301 ± 6.067 | 3.960 ± 3.592 | 5.104 ± 4.148 | 3.087 ± 2.811 | 4.783 ± 4.747 |
| HEIGHT | PMDA | 5.764 ± 5.037 | 3.495 ± 3.292 | 5.521 ± 4.868 | 2.823 ± 2.739 | 4.585 ± 4.439 |
| HEIGHT | OMNI2.5 | 2.526 ± 2.320 | 1.899 ± 1.692 | 2.396 ± 2.048 | 1.418 ± 1.357 | 2.338 ± 2.185 |
| HEIGHT | OMNI5 | 2.089 ± 1.963 | 1.178 ± 1.086 | 1.875 ± 1.680 | 0.942 ± 0.919 | 1.558 ± 1.413 |
| HEIGHT | LPS_0.5 | 4.863 ± 4.434 | 3.360 ± 2.970 | 5.285 ± 4.917 | 3.085 ± 2.695 | 4.745 ± 4.480 |
| HEIGHT | LPS_0.75 | 4.382 ± 4.100 | 3.246 ± 2.789 | 4.460 ± 3.957 | 2.839 ± 2.477 | 4.020 ± 3.789 |
| HEIGHT | LPS_1.0 | 3.796 ± 3.501 | 2.823 ± 2.460 | 4.276 ± 3.765 | 2.383 ± 2.154 | 3.545 ± 3.336 |
| HEIGHT | LPS_1.25 | 3.646 ± 3.310 | 2.657 ± 2.500 | 3.807 ± 3.323 | 2.194 ± 1.965 | 3.321 ± 3.248 |
| HEIGHT | LPS_1.5 | 3.480 ± 3.363 | 2.396 ± 2.142 | 3.424 ± 3.056 | 1.939 ± 1.714 | 3.135 ± 3.015 |
| HEIGHT | LPS_2.0 | 3.001 ± 2.639 | 2.122 ± 1.895 | 3.244 ± 2.933 | 1.928 ± 1.713 | 2.895 ± 2.616 |
| METABOLIC | GSA | 7.622 ± 6.709 | 4.053 ± 3.627 | 7.052 ± 6.317 | 4.555 ± 4.014 | 5.931 ± 5.030 |
| METABOLIC | JAPONICA | 6.127 ± 5.673 | 3.362 ± 3.317 | 4.408 ± 4.030 | 4.340 ± 3.837 | 4.655 ± 4.222 |
| METABOLIC | UKB_WCSG | 7.194 ± 6.918 | 2.869 ± 2.673 | 6.019 ± 5.165 | 2.876 ± 2.671 | 3.902 ± 3.620 |
| METABOLIC | CYTOSNP | 4.181 ± 3.742 | 2.470 ± 2.310 | 4.203 ± 3.609 | 3.045 ± 2.946 | 3.499 ± 3.063 |
| METABOLIC | PMRA | 5.969 ± | 4.023 ± | 6.404 ± | 4.032 ± | 5.132 ± |

| | | | | | | |
|---|---|---|---|---|---|---|
| | | 5.816 | 3.792 | 5.558 | 3.582 | 4.754 |
| METABOLIC | PMDA | 5.478 ± 4.987 | 3.267 ± 3.162 | 6.072 ± 5.553 | 3.871 ± 3.723 | 4.631 ± 4.246 |
| METABOLIC | OMNI2.5 | 2.477 ± 2.317 | 1.762 ± 1.689 | 2.686 ± 2.497 | 1.961 ± 1.874 | 2.242 ± 2.016 |
| METABOLIC | OMNI5 | 1.882 ± 1.808 | 1.209 ± 1.189 | 1.851 ± 1.684 | 1.208 ± 1.125 | 1.611 ± 1.459 |
| METABOLIC | LPS_0.5 | 4.361 ± 4.071 | 3.352 ± 3.164 | 5.859 ± 5.169 | 3.962 ± 3.596 | 4.855 ± 4.213 |
| METABOLIC | LPS_0.75 | 4.044 ± 3.744 | 2.824 ± 2.633 | 4.846 ± 4.348 | 3.179 ± 2.960 | 3.898 ± 3.403 |
| METABOLIC | LPS_1.0 | 3.577 ± 3.340 | 2.521 ± 2.357 | 4.318 ± 3.908 | 3.131 ± 2.769 | 3.570 ± 3.103 |
| METABOLIC | LPS_1.25 | 3.182 ± 2.941 | 2.282 ± 2.272 | 3.943 ± 3.670 | 2.684 ± 2.605 | 3.378 ± 2.996 |
| METABOLIC | LPS_1.5 | 2.867 ± 2.755 | 2.350 ± 2.246 | 3.686 ± 3.465 | 2.605 ± 2.306 | 3.075 ± 2.705 |
| METABOLIC | LPS_2.0 | 2.626 ± 2.540 | 2.108 ± 1.976 | 3.076 ± 2.841 | 2.260 ± 2.013 | 2.707 ± 2.433 |

Table S. 20 Mean absolute difference of percentile ranking between PGSs estimated from imputed genotyping data of eight genotyping arrays and six LPS coverages and PGS estimated from WGS in 5 different populations with PRsice p-value setting of 0.3

| Trait | Array/LPS | AFR | AMR | EAS | EUR | SAS |
|---|---|---|---|---|---|---|
| BMI | GSA | 7.658 ± 6.875 | 4.623 ± 4.173 | 7.289 ± 6.801 | 5.437 ± 4.746 | 5.971 ± 5.267 |
| BMI | JAPONICA | 6.654 ± 6.232 | 4.153 ± 3.792 | 4.634 ± 4.028 | 4.611 ± 4.150 | 4.990 ± 4.207 |
| BMI | UKB_WCSG | 7.482 ± 6.908 | 3.427 ± 2.892 | 5.974 ± 5.420 | 3.130 ± 2.953 | 4.184 ± 3.691 |
| BMI | CYTOSNP | 4.626 ± 4.448 | 3.119 ± 2.784 | 4.725 ± 3.837 | 3.816 ± 3.417 | 4.217 ± 3.586 |
| BMI | PMRA | 6.654 ± 6.310 | 4.157 ± 3.705 | 6.373 ± 6.135 | 4.979 ± 4.557 | 5.609 ± 5.013 |
| BMI | PMDA | 6.309 ± 5.858 | 3.913 ± 3.245 | 6.294 ± 5.564 | 4.456 ± 4.103 | 5.041 ± 4.272 |
| BMI | OMNI2.5 | 2.525 ± 2.528 | 2.021 ± 1.630 | 2.968 ± 2.496 | 2.092 ± 1.992 | 2.417 ± 2.136 |
| BMI | OMNI5 | 2.132 ± 2.054 | 1.339 ± 1.139 | 2.115 ± 1.947 | 1.334 ± 1.354 | 1.671 ± 1.466 |
| BMI | LPS_0.5 | 5.040 ± 5.034 | 4.028 ± 3.458 | 6.035 ± 5.250 | 4.636 ± 4.225 | 4.908 ± 4.362 |
| BMI | LPS_0.75 | 4.173 ± 3.963 | 3.555 ± 3.247 | 5.528 ± 5.079 | 3.682 ± 3.282 | 4.331 ± 3.813 |
| BMI | LPS_1.0 | 3.749 ± 3.542 | 2.756 ± 2.537 | 4.766 ± 4.064 | 3.329 ± 3.074 | 3.697 ± 3.204 |
| BMI | LPS_1.25 | 3.491 ± 3.296 | 2.959 ± 2.512 | 4.249 ± 3.848 | 3.201 ± 3.020 | 3.352 ± 3.064 |
| BMI | LPS_1.5 | 3.104 ± 3.033 | 2.604 ± 2.333 | 3.819 ± 3.452 | 2.711 ± 2.644 | 3.255 ± 3.084 |
| BMI | LPS_2.0 | 2.899 ± 2.636 | 2.462 ± 2.224 | 3.599 ± 3.055 | 2.448 ± 2.113 | 2.879 ± 2.521 |
| DIABETES | GSA | 7.335 ± 6.928 | 4.209 ± 3.808 | 7.827 ± 7.331 | 4.537 ± 4.214 | 5.661 ± 4.870 |
| DIABETES | JAPONICA | 6.581 ± 6.521 | 3.457 ± 3.063 | 4.626 ± 4.337 | 4.046 ± 3.689 | 4.642 ± 4.185 |
| DIABETES | UKB_WCSG | 7.022 ± 6.811 | 3.245 ± 2.839 | 6.467 ± 5.821 | 3.017 ± 2.708 | 4.560 ± 4.315 |
| DIABETES | CYTOSNP | 4.691 ± 4.600 | 2.337 ± 2.300 | 4.587 ± 4.072 | 2.731 ± 2.395 | 3.724 ± 3.342 |
| DIABETES | PMRA | 6.350 ± 5.903 | 3.732 ± 3.731 | 6.742 ± 5.962 | 3.931 ± 3.644 | 5.525 ± 5.133 |
| DIABETES | PMDA | 5.880 ± 5.787 | 3.496 ± 3.118 | 6.441 ± 5.892 | 3.550 ± 3.038 | 4.751 ± 4.452 |
| DIABETES | OMNI2.5 | 2.578 ± 2.472 | 1.623 ± 1.441 | 3.053 ± 2.721 | 1.759 ± 1.599 | 2.396 ± 2.168 |
| DIABETES | OMNI5 | 2.305 ± 2.207 | 1.208 ± 1.158 | 2.422 ± 2.238 | 1.296 ± 1.216 | 1.998 ± 1.884 |

| | | | | | | |
|---|---|---|---|---|---|---|
| DIABETES | LPS_0.5 | 4.615 ± 4.475 | 3.528 ± 2.999 | 6.087 ± 5.135 | 3.714 ± 3.422 | 4.443 ± 4.194 |
| DIABETES | LPS_0.75 | 3.931 ± 4.007 | 3.166 ± 2.765 | 4.981 ± 4.491 | 3.137 ± 2.768 | 4.056 ± 3.773 |
| DIABETES | LPS_1.0 | 3.593 ± 3.442 | 2.408 ± 2.182 | 4.488 ± 4.247 | 2.763 ± 2.652 | 3.359 ± 3.070 |
| DIABETES | LPS_1.25 | 3.393 ± 3.277 | 2.471 ± 2.220 | 3.957 ± 3.478 | 2.578 ± 2.227 | 3.051 ± 2.864 |
| DIABETES | LPS_1.5 | 3.330 ± 3.197 | 2.040 ± 1.999 | 3.851 ± 3.549 | 2.454 ± 2.233 | 3.132 ± 2.746 |
| DIABETES | LPS_2.0 | 2.817 ± 2.793 | 2.103 ± 1.962 | 3.451 ± 3.193 | 2.100 ± 1.936 | 2.676 ± 2.531 |
| HEIGHT | GSA | 7.841 ± 6.974 | 4.044 ± 3.603 | 6.013 ± 5.257 | 3.588 ± 3.235 | 5.786 ± 5.129 |
| HEIGHT | JAPONICA | 6.377 ± 5.656 | 3.620 ± 3.354 | 4.120 ± 3.469 | 3.248 ± 2.886 | 4.654 ± 4.591 |
| HEIGHT | UKB_WCSG | 6.806 ± 6.116 | 2.930 ± 2.558 | 4.984 ± 4.377 | 2.081 ± 1.929 | 3.981 ± 3.627 |
| HEIGHT | CYTOSNP | 4.472 ± 4.185 | 3.026 ± 2.673 | 3.693 ± 3.313 | 2.358 ± 2.237 | 3.596 ± 3.407 |
| HEIGHT | PMRA | 6.326 ± 6.055 | 3.881 ± 3.545 | 5.249 ± 4.287 | 3.092 ± 2.792 | 4.751 ± 4.636 |
| HEIGHT | PMDA | 5.754 ± 4.995 | 3.457 ± 3.258 | 5.543 ± 4.933 | 2.807 ± 2.641 | 4.559 ± 4.352 |
| HEIGHT | OMNI2.5 | 2.519 ± 2.303 | 1.859 ± 1.703 | 2.370 ± 2.044 | 1.427 ± 1.314 | 2.304 ± 2.125 |
| HEIGHT | OMNI5 | 2.040 ± 1.931 | 1.125 ± 1.035 | 1.818 ± 1.702 | 0.944 ± 0.942 | 1.526 ± 1.420 |
| HEIGHT | LPS_0.5 | 4.852 ± 4.500 | 3.325 ± 2.883 | 5.280 ± 4.930 | 3.102 ± 2.709 | 4.756 ± 4.430 |
| HEIGHT | LPS_0.75 | 4.378 ± 4.064 | 3.179 ± 2.817 | 4.456 ± 4.105 | 2.817 ± 2.467 | 3.986 ± 3.743 |
| HEIGHT | LPS_1.0 | 3.767 ± 3.493 | 2.800 ± 2.397 | 4.321 ± 3.895 | 2.343 ± 2.163 | 3.576 ± 3.306 |
| HEIGHT | LPS_1.25 | 3.590 ± 3.276 | 2.736 ± 2.579 | 3.786 ± 3.357 | 2.188 ± 2.045 | 3.307 ± 3.194 |
| HEIGHT | LPS_1.5 | 3.517 ± 3.297 | 2.428 ± 2.117 | 3.442 ± 3.058 | 1.934 ± 1.759 | 3.117 ± 2.988 |
| HEIGHT | LPS_2.0 | 3.016 ± 2.708 | 2.213 ± 1.971 | 3.303 ± 2.935 | 1.879 ± 1.725 | 2.909 ± 2.646 |
| METABOLIC | GSA | 7.482 ± 6.585 | 3.998 ± 3.555 | 7.255 ± 6.608 | 4.565 ± 4.094 | 5.888 ± 5.248 |
| METABOLIC | JAPONICA | 6.195 ± 5.751 | 3.289 ± 3.193 | 4.679 ± 4.195 | 4.290 ± 3.712 | 4.783 ± 4.378 |
| METABOLIC | UKB_WCSG | 7.142 ± 6.786 | 2.874 ± 2.614 | 6.172 ± 5.298 | 2.854 ± 2.695 | 3.821 ± 3.592 |
| METABOLIC | CYTOSNP | 4.258 ± 3.905 | 2.536 ± 2.356 | 4.392 ± 3.747 | 3.116 ± 3.037 | 3.509 ± 2.996 |
| METABOLIC | PMRA | 5.921 ± | 4.043 ± | 6.450 ± | 4.043 ± | 5.195 ± |

| | | | | | | |
|---|---|---|---|---|---|---|
| | | 5.564 | 3.630 | 5.848 | 3.607 | 4.829 |
| METABOLIC | PMDA | 5.395 ± 5.001 | 3.222 ± 3.009 | 6.220 ± 5.748 | 3.868 ± 3.727 | 4.843 ± 4.381 |
| METABOLIC | OMNI2.5 | 2.443 ± 2.319 | 1.726 ± 1.670 | 2.787 ± 2.591 | 1.956 ± 1.894 | 2.249 ± 1.969 |
| METABOLIC | OMNI5 | 1.812 ± 1.705 | 1.193 ± 1.160 | 1.947 ± 1.815 | 1.225 ± 1.178 | 1.596 ± 1.450 |
| METABOLIC | LPS_0.5 | 4.373 ± 4.055 | 3.232 ± 2.937 | 5.970 ± 5.245 | 4.001 ± 3.636 | 4.835 ± 4.229 |
| METABOLIC | LPS_0.75 | 3.962 ± 3.636 | 2.809 ± 2.542 | 4.939 ± 4.613 | 3.337 ± 3.056 | 4.004 ± 3.510 |
| METABOLIC | LPS_1.0 | 3.554 ± 3.319 | 2.518 ± 2.298 | 4.469 ± 4.030 | 3.133 ± 2.782 | 3.676 ± 3.234 |
| METABOLIC | LPS_1.25 | 3.136 ± 2.922 | 2.261 ± 2.142 | 4.086 ± 3.781 | 2.729 ± 2.609 | 3.441 ± 2.914 |
| METABOLIC | LPS_1.5 | 2.846 ± 2.766 | 2.382 ± 2.173 | 3.740 ± 3.513 | 2.649 ± 2.423 | 3.070 ± 2.694 |
| METABOLIC | LPS_2.0 | 2.576 ± 2.472 | 2.090 ± 1.961 | 3.134 ± 2.872 | 2.288 ± 2.029 | 2.807 ± 2.578 |

Table S. 21 Mean absolute difference of percentile ranking between PGSs estimated from imputed genotyping data of eight genotyping arrays and six LPS coverages and PGS estimated from WGS in 5 different populations with PRsice p-value setting of 0.5

| Trait | Array/LPS | AFR | AMR | EAS | EUR | SAS |
|---|---|---|---|---|---|---|
| BMI | GSA | 7.676 ± 7.077 | 4.646 ± 4.127 | 7.376 ± 6.985 | 5.465 ± 4.926 | 6.040 ± 5.275 |
| BMI | JAPONICA | 6.841 ± 6.526 | 4.246 ± 3.885 | 4.814 ± 4.203 | 4.577 ± 4.154 | 4.825 ± 4.247 |
| BMI | UKB_WCSG | 7.428 ± 6.868 | 3.273 ± 2.837 | 6.004 ± 5.517 | 3.101 ± 2.944 | 4.177 ± 3.757 |
| BMI | CYTOSNP | 4.595 ± 4.501 | 3.123 ± 2.714 | 4.858 ± 3.983 | 3.849 ± 3.380 | 4.185 ± 3.526 |
| BMI | PMRA | 6.674 ± 6.343 | 4.077 ± 3.465 | 6.394 ± 6.292 | 4.931 ± 4.598 | 5.591 ± 4.970 |
| BMI | PMDA | 6.317 ± 5.951 | 3.888 ± 3.256 | 6.354 ± 5.697 | 4.441 ± 4.161 | 4.995 ± 4.248 |
| BMI | OMNI2.5 | 2.521 ± 2.458 | 2.034 ± 1.638 | 2.869 ± 2.556 | 2.166 ± 2.084 | 2.378 ± 2.128 |
| BMI | OMNI5 | 2.116 ± 2.005 | 1.391 ± 1.195 | 2.125 ± 1.885 | 1.387 ± 1.368 | 1.625 ± 1.483 |
| BMI | LPS_0.5 | 5.132 ± 4.949 | 4.070 ± 3.490 | 6.187 ± 5.419 | 4.636 ± 4.284 | 4.945 ± 4.424 |
| BMI | LPS_0.75 | 4.244 ± 4.143 | 3.623 ± 3.060 | 5.615 ± 5.113 | 3.759 ± 3.347 | 4.325 ± 3.728 |
| BMI | LPS_1.0 | 3.790 ± 3.575 | 2.847 ± 2.623 | 4.805 ± 4.149 | 3.388 ± 3.142 | 3.635 ± 3.340 |
| BMI | LPS_1.25 | 3.561 ± 3.426 | 2.844 ± 2.411 | 4.284 ± 3.861 | 3.193 ± 3.071 | 3.309 ± 3.061 |
| BMI | LPS_1.5 | 3.230 ± 3.165 | 2.492 ± 2.319 | 3.803 ± 3.516 | 2.799 ± 2.527 | 3.270 ± 2.978 |
| BMI | LPS_2.0 | 2.964 ± 2.720 | 2.406 ± 2.145 | 3.649 ± 2.999 | 2.534 ± 2.189 | 2.840 ± 2.531 |
| DIABETES | GSA | 7.353 ± 6.740 | 4.171 ± 3.753 | 7.812 ± 7.232 | 4.505 ± 4.066 | 5.696 ± 4.995 |
| DIABETES | JAPONICA | 6.602 ± 6.280 | 3.628 ± 3.274 | 4.719 ± 4.412 | 4.044 ± 3.832 | 4.839 ± 4.358 |
| DIABETES | UKB_WCSG | 7.083 ± 6.681 | 3.146 ± 2.885 | 6.622 ± 5.938 | 3.066 ± 2.763 | 4.524 ± 4.274 |
| DIABETES | CYTOSNP | 4.642 ± 4.485 | 2.470 ± 2.255 | 4.518 ± 4.015 | 2.755 ± 2.338 | 3.762 ± 3.589 |
| DIABETES | PMRA | 6.412 ± 5.816 | 3.971 ± 3.790 | 6.679 ± 5.833 | 4.056 ± 3.679 | 5.657 ± 4.969 |
| DIABETES | PMDA | 5.902 ± 5.620 | 3.636 ± 3.118 | 6.674 ± 5.829 | 3.721 ± 3.207 | 4.774 ± 4.655 |
| DIABETES | OMNI2.5 | 2.617 ± 2.351 | 1.658 ± 1.557 | 3.010 ± 2.668 | 1.744 ± 1.570 | 2.373 ± 2.240 |
| DIABETES | OMNI5 | 2.218 ± 2.083 | 1.232 ± 1.205 | 2.431 ± 2.192 | 1.261 ± 1.171 | 1.974 ± 1.883 |

| | | | | | | |
|---|---|---|---|---|---|---|
| DIABETES | LPS_0.5 | 4.647 ± 4.283 | 3.485 ± 2.982 | 6.120 ± 5.399 | 3.747 ± 3.450 | 4.417 ± 4.179 |
| DIABETES | LPS_0.75 | 4.035 ± 3.964 | 3.133 ± 2.867 | 5.050 ± 4.429 | 3.194 ± 2.871 | 4.075 ± 3.811 |
| DIABETES | LPS_1.0 | 3.622 ± 3.287 | 2.447 ± 2.242 | 4.503 ± 4.278 | 2.783 ± 2.650 | 3.424 ± 3.063 |
| DIABETES | LPS_1.25 | 3.435 ± 3.406 | 2.510 ± 2.202 | 3.921 ± 3.447 | 2.564 ± 2.193 | 3.058 ± 2.748 |
| DIABETES | LPS_1.5 | 3.385 ± 3.068 | 2.174 ± 2.142 | 3.825 ± 3.482 | 2.405 ± 2.174 | 3.250 ± 2.875 |
| DIABETES | LPS_2.0 | 2.758 ± 2.738 | 2.070 ± 1.887 | 3.685 ± 3.210 | 2.170 ± 1.986 | 2.759 ± 2.594 |
| HEIGHT | GSA | 7.849 ± 7.013 | 4.124 ± 3.635 | 6.064 ± 5.136 | 3.570 ± 3.252 | 5.803 ± 5.202 |
| HEIGHT | JAPONICA | 6.300 ± 5.671 | 3.698 ± 3.401 | 4.175 ± 3.510 | 3.153 ± 2.791 | 4.552 ± 4.490 |
| HEIGHT | UKB_WCSG | 6.746 ± 6.042 | 2.928 ± 2.635 | 4.986 ± 4.415 | 2.087 ± 1.942 | 3.963 ± 3.639 |
| HEIGHT | CYTOSNP | 4.440 ± 4.175 | 3.022 ± 2.773 | 3.703 ± 3.219 | 2.318 ± 2.231 | 3.631 ± 3.345 |
| HEIGHT | PMRA | 6.223 ± 6.046 | 3.881 ± 3.628 | 5.238 ± 4.280 | 3.117 ± 2.802 | 4.794 ± 4.615 |
| HEIGHT | PMDA | 5.656 ± 5.029 | 3.457 ± 3.261 | 5.560 ± 4.966 | 2.797 ± 2.609 | 4.570 ± 4.398 |
| HEIGHT | OMNI2.5 | 2.515 ± 2.262 | 1.885 ± 1.707 | 2.373 ± 2.049 | 1.419 ± 1.272 | 2.249 ± 2.094 |
| HEIGHT | OMNI5 | 2.060 ± 1.929 | 1.177 ± 1.100 | 1.895 ± 1.730 | 0.931 ± 0.940 | 1.523 ± 1.370 |
| HEIGHT | LPS_0.5 | 4.919 ± 4.513 | 3.278 ± 2.887 | 5.248 ± 4.878 | 3.035 ± 2.736 | 4.761 ± 4.394 |
| HEIGHT | LPS_0.75 | 4.353 ± 3.989 | 3.163 ± 2.852 | 4.504 ± 4.097 | 2.844 ± 2.466 | 4.012 ± 3.787 |
| HEIGHT | LPS_1.0 | 3.850 ± 3.504 | 2.821 ± 2.398 | 4.308 ± 3.782 | 2.323 ± 2.183 | 3.669 ± 3.423 |
| HEIGHT | LPS_1.25 | 3.601 ± 3.260 | 2.692 ± 2.511 | 3.843 ± 3.273 | 2.217 ± 2.069 | 3.351 ± 3.282 |
| HEIGHT | LPS_1.5 | 3.468 ± 3.323 | 2.430 ± 2.154 | 3.442 ± 3.067 | 1.951 ± 1.778 | 3.152 ± 2.934 |
| HEIGHT | LPS_2.0 | 3.040 ± 2.727 | 2.264 ± 2.054 | 3.307 ± 2.972 | 1.937 ± 1.755 | 2.913 ± 2.691 |
| METABOLIC | GSA | 7.394 ± 6.633 | 3.985 ± 3.388 | 7.344 ± 6.662 | 4.561 ± 4.026 | 5.825 ± 5.175 |
| METABOLIC | JAPONICA | 6.156 ± 5.831 | 3.176 ± 3.100 | 4.732 ± 4.261 | 4.154 ± 3.720 | 4.852 ± 4.451 |
| METABOLIC | UKB_WCSG | 7.076 ± 6.754 | 2.781 ± 2.610 | 6.206 ± 5.368 | 2.864 ± 2.683 | 3.769 ± 3.478 |
| METABOLIC | CYTOSNP | 4.275 ± 3.982 | 2.483 ± 2.295 | 4.598 ± 3.931 | 3.130 ± 2.994 | 3.631 ± 3.048 |
| METABOLIC | PMRA | 5.870 ± | 3.922 ± | 6.502 ± | 4.100 ± | 5.202 ± |

| | | | | | | |
|---|---|---|---|---|---|---|
| | | 5.552 | 3.594 | 5.852 | 3.678 | 4.742 |
| METABOLIC | PMDA | 5.357 ± 4.941 | 3.144 ± 2.918 | 6.226 ± 5.863 | 3.783 ± 3.667 | 4.843 ± 4.336 |
| METABOLIC | OMNI2.5 | 2.414 ± 2.358 | 1.663 ± 1.644 | 2.775 ± 2.612 | 1.881 ± 1.821 | 2.296 ± 2.058 |
| METABOLIC | OMNI5 | 1.879 ± 1.787 | 1.154 ± 1.114 | 2.005 ± 1.810 | 1.186 ± 1.139 | 1.649 ± 1.551 |
| METABOLIC | LPS_0.5 | 4.381 ± 4.196 | 3.217 ± 3.087 | 6.094 ± 5.391 | 3.942 ± 3.595 | 4.840 ± 4.280 |
| METABOLIC | LPS_0.75 | 3.980 ± 3.691 | 2.789 ± 2.616 | 5.015 ± 4.565 | 3.220 ± 2.909 | 4.062 ± 3.616 |
| METABOLIC | LPS_1.0 | 3.541 ± 3.345 | 2.437 ± 2.231 | 4.603 ± 4.077 | 3.020 ± 2.831 | 3.647 ± 3.162 |
| METABOLIC | LPS_1.25 | 3.187 ± 3.040 | 2.330 ± 2.222 | 4.130 ± 3.861 | 2.669 ± 2.534 | 3.489 ± 3.019 |
| METABOLIC | LPS_1.5 | 2.898 ± 2.856 | 2.246 ± 2.157 | 3.802 ± 3.565 | 2.665 ± 2.431 | 3.096 ± 2.805 |
| METABOLIC | LPS_2.0 | 2.564 ± 2.500 | 2.095 ± 1.911 | 3.257 ± 2.952 | 2.202 ± 2.014 | 2.866 ± 2.625 |

Table S. 22 Mean absolute difference of percentile ranking between PGSs estimated from imputed genotyping data of eight genotyping arrays and six LPS coverages and PGS estimated from WGS in 5 different populations with PRsice p-value setting of 1

| Trait | Array/LPS | AFR | AMR | EAS | EUR | SAS |
|---|---|---|---|---|---|---|
| BMI | GSA | 7.680 ± 6.987 | 4.621 ± 3.999 | 7.455 ± 6.995 | 5.459 ± 4.807 | 6.070 ± 5.291 |
| BMI | JAPONICA | 6.778 ± 6.574 | 4.216 ± 3.847 | 4.780 ± 4.145 | 4.626 ± 4.221 | 4.796 ± 4.313 |
| BMI | UKB_WCSG | 7.424 ± 6.814 | 3.223 ± 2.839 | 6.005 ± 5.499 | 3.108 ± 2.960 | 4.287 ± 3.686 |
| BMI | CYTOSNP | 4.642 ± 4.570 | 2.998 ± 2.653 | 4.960 ± 3.964 | 3.839 ± 3.315 | 4.107 ± 3.566 |
| BMI | PMRA | 6.681 ± 6.432 | 3.981 ± 3.372 | 6.369 ± 6.270 | 4.905 ± 4.575 | 5.586 ± 4.901 |
| BMI | PMDA | 6.249 ± 5.923 | 3.793 ± 3.259 | 6.290 ± 5.728 | 4.410 ± 4.161 | 5.119 ± 4.275 |
| BMI | OMNI2.5 | 2.494 ± 2.414 | 1.916 ± 1.620 | 2.945 ± 2.554 | 2.148 ± 2.063 | 2.457 ± 2.119 |
| BMI | OMNI5 | 2.125 ± 1.998 | 1.361 ± 1.260 | 2.133 ± 1.898 | 1.360 ± 1.371 | 1.673 ± 1.482 |
| BMI | LPS_0.5 | 5.090 ± 4.938 | 3.899 ± 3.350 | 6.190 ± 5.476 | 4.605 ± 4.251 | 5.016 ± 4.404 |
| BMI | LPS_0.75 | 4.262 ± 4.181 | 3.547 ± 3.106 | 5.680 ± 5.068 | 3.786 ± 3.322 | 4.238 ± 3.727 |
| BMI | LPS_1.0 | 3.920 ± 3.637 | 2.855 ± 2.576 | 4.829 ± 4.183 | 3.416 ± 3.096 | 3.687 ± 3.340 |
| BMI | LPS_1.25 | 3.613 ± 3.498 | 2.790 ± 2.477 | 4.293 ± 3.825 | 3.156 ± 3.104 | 3.350 ± 3.079 |
| BMI | LPS_1.5 | 3.305 ± 3.225 | 2.536 ± 2.214 | 3.871 ± 3.498 | 2.785 ± 2.541 | 3.303 ± 3.019 |
| BMI | LPS_2.0 | 2.996 ± 2.757 | 2.404 ± 2.115 | 3.596 ± 2.968 | 2.545 ± 2.166 | 2.911 ± 2.506 |
| DIABETES | GSA | 7.397 ± 6.849 | 4.111 ± 3.654 | 7.841 ± 7.319 | 4.528 ± 4.122 | 5.634 ± 4.951 |
| DIABETES | JAPONICA | 6.614 ± 6.274 | 3.741 ± 3.444 | 4.811 ± 4.371 | 4.083 ± 3.897 | 4.892 ± 4.315 |
| DIABETES | UKB_WCSG | 7.208 ± 6.785 | 3.246 ± 2.997 | 6.603 ± 5.897 | 3.073 ± 2.841 | 4.527 ± 4.262 |
| DIABETES | CYTOSNP | 4.716 ± 4.501 | 2.568 ± 2.380 | 4.670 ± 4.130 | 2.746 ± 2.425 | 3.740 ± 3.467 |
| DIABETES | PMRA | 6.456 ± 5.851 | 3.922 ± 3.765 | 6.765 ± 5.830 | 4.130 ± 3.660 | 5.565 ± 4.967 |
| DIABETES | PMDA | 5.923 ± 5.657 | 3.634 ± 3.138 | 6.602 ± 5.851 | 3.657 ± 3.222 | 4.790 ± 4.573 |
| DIABETES | OMNI2.5 | 2.627 ± 2.404 | 1.664 ± 1.519 | 3.058 ± 2.707 | 1.749 ± 1.665 | 2.348 ± 2.211 |
| DIABETES | OMNI5 | 2.232 ± 2.125 | 1.242 ± 1.247 | 2.458 ± 2.231 | 1.271 ± 1.198 | 2.001 ± 1.924 |

| | | | | | | |
|---|---|---|---|---|---|---|
| DIABETES | LPS_0.5 | 4.653 ± 4.321 | 3.563 ± 3.110 | 6.102 ± 5.390 | 3.728 ± 3.523 | 4.411 ± 4.214 |
| DIABETES | LPS_0.75 | 3.979 ± 3.944 | 3.144 ± 2.996 | 5.010 ± 4.443 | 3.180 ± 2.878 | 4.198 ± 3.799 |
| DIABETES | LPS_1.0 | 3.620 ± 3.286 | 2.555 ± 2.337 | 4.500 ± 4.262 | 2.867 ± 2.716 | 3.424 ± 3.097 |
| DIABETES | LPS_1.25 | 3.380 ± 3.375 | 2.548 ± 2.298 | 3.941 ± 3.491 | 2.566 ± 2.271 | 3.091 ± 2.752 |
| DIABETES | LPS_1.5 | 3.340 ± 3.039 | 2.159 ± 2.015 | 3.938 ± 3.513 | 2.436 ± 2.305 | 3.241 ± 2.953 |
| DIABETES | LPS_2.0 | 2.795 ± 2.752 | 2.103 ± 1.965 | 3.635 ± 3.194 | 2.271 ± 2.091 | 2.780 ± 2.649 |
| HEIGHT | GSA | 7.835 ± 7.004 | 4.150 ± 3.749 | 6.026 ± 5.136 | 3.576 ± 3.262 | 5.795 ± 5.238 |
| HEIGHT | JAPONICA | 6.258 ± 5.640 | 3.702 ± 3.443 | 4.239 ± 3.591 | 3.153 ± 2.783 | 4.540 ± 4.520 |
| HEIGHT | UKB_WCSG | 6.751 ± 6.042 | 2.981 ± 2.588 | 5.009 ± 4.405 | 2.085 ± 1.950 | 3.980 ± 3.625 |
| HEIGHT | CYTOSNP | 4.465 ± 4.227 | 3.074 ± 2.885 | 3.750 ± 3.235 | 2.350 ± 2.239 | 3.599 ± 3.344 |
| HEIGHT | PMRA | 6.318 ± 6.108 | 3.961 ± 3.592 | 5.189 ± 4.266 | 3.102 ± 2.842 | 4.775 ± 4.639 |
| HEIGHT | PMDA | 5.655 ± 5.023 | 3.466 ± 3.366 | 5.582 ± 4.963 | 2.775 ± 2.583 | 4.537 ± 4.457 |
| HEIGHT | OMNI2.5 | 2.484 ± 2.262 | 1.948 ± 1.797 | 2.390 ± 1.982 | 1.424 ± 1.299 | 2.349 ± 2.157 |
| HEIGHT | OMNI5 | 2.043 ± 1.888 | 1.200 ± 1.152 | 1.871 ± 1.695 | 0.953 ± 0.910 | 1.524 ± 1.398 |
| HEIGHT | LPS_0.5 | 4.870 ± 4.477 | 3.288 ± 2.987 | 5.263 ± 4.980 | 2.984 ± 2.773 | 4.739 ± 4.392 |
| HEIGHT | LPS_0.75 | 4.304 ± 3.922 | 3.219 ± 2.927 | 4.455 ± 4.094 | 2.823 ± 2.478 | 4.002 ± 3.791 |
| HEIGHT | LPS_1.0 | 3.856 ± 3.526 | 2.848 ± 2.495 | 4.326 ± 3.771 | 2.332 ± 2.152 | 3.619 ± 3.375 |
| HEIGHT | LPS_1.25 | 3.582 ± 3.210 | 2.751 ± 2.621 | 3.828 ± 3.258 | 2.217 ± 2.064 | 3.313 ± 3.207 |
| HEIGHT | LPS_1.5 | 3.438 ± 3.302 | 2.434 ± 2.174 | 3.486 ± 3.064 | 1.965 ± 1.752 | 3.176 ± 2.935 |
| HEIGHT | LPS_2.0 | 3.066 ± 2.689 | 2.251 ± 2.131 | 3.325 ± 2.973 | 1.948 ± 1.741 | 2.901 ± 2.667 |
| METABOLIC | GSA | 7.273 ± 6.589 | 3.927 ± 3.311 | 7.379 ± 6.677 | 4.554 ± 4.032 | 5.718 ± 5.135 |
| METABOLIC | JAPONICA | 6.095 ± 5.734 | 3.247 ± 3.086 | 4.748 ± 4.219 | 4.236 ± 3.754 | 4.869 ± 4.409 |
| METABOLIC | UKB_WCSG | 7.105 ± 6.747 | 2.696 ± 2.534 | 6.221 ± 5.326 | 2.854 ± 2.756 | 3.749 ± 3.395 |
| METABOLIC | CYTOSNP | 4.274 ± 4.000 | 2.460 ± 2.170 | 4.532 ± 3.921 | 3.160 ± 3.063 | 3.608 ± 3.071 |
| METABOLIC | PMRA | 5.841 ± | 3.827 ± | 6.593 ± | 4.177 ± | 5.157 ± |

| | | | | | | |
|---|---|---|---|---|---|---|
| | | 5.444 | 3.471 | 5.876 | 3.682 | 4.712 |
| METABOLIC | PMDA | 5.293 ± 4.887 | 3.133 ± 2.828 | 6.244 ± 5.897 | 3.870 ± 3.699 | 4.874 ± 4.295 |
| METABOLIC | OMNI2.5 | 2.427 ± 2.329 | 1.711 ± 1.636 | 2.772 ± 2.554 | 1.953 ± 1.819 | 2.305 ± 2.083 |
| METABOLIC | OMNI5 | 1.854 ± 1.701 | 1.126 ± 1.062 | 1.976 ± 1.756 | 1.228 ± 1.192 | 1.628 ± 1.544 |
| METABOLIC | LPS_0.5 | 4.362 ± 4.157 | 3.206 ± 3.026 | 6.170 ± 5.316 | 4.001 ± 3.632 | 4.810 ± 4.345 |
| METABOLIC | LPS_0.75 | 3.987 ± 3.644 | 2.771 ± 2.524 | 5.104 ± 4.636 | 3.206 ± 2.896 | 4.060 ± 3.628 |
| METABOLIC | LPS_1.0 | 3.551 ± 3.261 | 2.462 ± 2.258 | 4.541 ± 4.140 | 3.024 ± 2.844 | 3.634 ± 3.170 |
| METABOLIC | LPS_1.25 | 3.180 ± 3.012 | 2.211 ± 2.155 | 4.165 ± 3.938 | 2.632 ± 2.566 | 3.417 ± 2.890 |
| METABOLIC | LPS_1.5 | 2.805 ± 2.722 | 2.242 ± 2.019 | 3.884 ± 3.560 | 2.676 ± 2.455 | 3.056 ± 2.717 |
| METABOLIC | LPS_2.0 | 2.578 ± 2.464 | 2.061 ± 1.836 | 3.252 ± 2.990 | 2.226 ± 1.990 | 2.790 ± 2.540 |
